# Lifespan Normative Modeling of Brain Microstructure

**DOI:** 10.1101/2024.12.15.628527

**Authors:** Julio E. Villalón-Reina, Alyssa H. Zhu, Leila Nabulsi, Sophia I. Thomopoulos, Clara A. Moreau, Yixue Feng, Tamoghna Chattopadhyay, Sebastian Benavidez, Leila Kushan, John P. John, Himanshu Joshi, Iyad Ba Gari, Katherine E. Lawrence, Talia M. Nir, Neda Jahanshad, Carrie E. Bearden, Seyed Mostafa Kia, Andre F. Marquand, the Alzheimer’s Disease Neuroimaging Initiative, Paul M. Thompson

## Abstract

Normative models of brain metrics based on large populations could be extremely valuable for detecting brain abnormalities in patients with a variety of disorders, including degenerative, psychiatric and neurodevelopmental conditions, but no such models exist for the brain’s white matter (WM) microstructure. Here we present a large-scale normative model of brain WM microstructure – based on 19 international diffusion MRI datasets covering almost the entire lifespan (totaling N=54,583 individuals; age: 4-91 years). We extracted regional diffusion tensor imaging (DTI) metrics using a standardized analysis and quality control protocol and used hierarchical Bayesian regression (HBR) to model the statistical distribution of derived WM metrics as a function of age and sex. We extracted the average lifespan trajectories and corresponding centile curves for each WM region. We illustrate the utility of the method by applying it to detect and visualize profiles of WM microstructural deviations in a variety of contexts: in mild cognitive impairment, Alzheimer’s disease, and 22q11.2 deletion syndrome – a neurogenetic condition that markedly increases risk for schizophrenia. The resulting large-scale model provides a common reference to identify disease effects on the brain’s microstructure in individuals or groups, and to compare disorders, and discover factors affecting WM abnormalities. The derived normative models are a valuable resource publicly available to the community, adaptable and extendable to future datasets as the available data expands.

## 1. Introduction

Large-scale international initiatives have greatly increased the availability of brain imaging data, driving the development of advanced statistical tools such as normative^1^ and generative models^2^ to study brain measures across life, health, and disease. Normative modeling (NM) is a statistical technique that estimates the normative distribution, i.e., the centiles of variation, of a biological measure in a reference population, given explanatory or clinical variables such as age and sex. The resulting model can be used to gauge and track brain abnormalities and factors that influence them. NM is sensitive to individual differences and offers multivariate metrics of abnormality, extending the standard case-control maps of group differences. This has sparked broader interest in NM for comprehensive studies of neurological and psychiatric conditions, and to support personalized medicine^3^.

NM has previously been applied to anatomical brain scans to create lifespan trajectories for structural brain metrics. Rutherford et al.^4^ used the Predictive Clinical Neuroscience toolkit^1^ (https://pcntoolkit.readthedocs.io) to chart the lifespan trajectories for a range of structural brain measures, available at the PCNPortal^5^ (https://pcnportal.dccn.nl/). Bethlehem et al.^6^ aggregated structural MRI scans from over 100 studies, creating lifespan ‘brain charts’ from 101,457 human participants from birth to 100 years of age. Ge et al.^7^ applied algorithms including multivariable fractional polynomial regression, warped Bayesian linear regression and hierarchical Bayesian regression to model brain morphometry in 37,407 healthy individuals from 86 international sites as part of the ENIGMA-Lifespan project, creating CentileBrain (https://centilebrain.org). Similar approaches have been extended to infant brain data, most recently by the ENIGMA Origins group^8^.

So far, we lack lifespan normative models of the brain’s white matter (WM) microstructure, which is sensitive to complex maturation, aging, and disease processes not detectable with standard structural MRI. Diffusion magnetic resonance imaging (dMRI) is a non-invasive technique that is uniquely sensitive to the local microstructural environment of the brain^9^. This method can assess processes occurring in living brain tissue at the micrometer scale (∼10 µm), surpassing the voxel resolution of current clinical scanners (1-3 mm), and is therefore sensitive to cellular-level structures that restrict water diffusion in tissue, such as cellular membranes of neurons and glia, and myelin. Over the past two decades, dMRI has been instrumental in the study of white WM, providing a comprehensive set of metrics that are sensitive to changes across the lifespan and a wide array of neurological and psychiatric diseases^9,10^. While dMRI models can be applied to study gray matter using higher-order dMRI models and advanced multishell and multidimensional acquisition protocols, we focus on diffusion tensor imaging (DTI), arguably the most widely used dMRI model for studying the WM. On the other hand, several microstructural properties of the gray matter (e.g. water exchange between the intra– and the extra-cellular space; non-Gaussian diffusion along neuronal and glial processes, and the signal contribution from soma) limit the applicability of DTI to study the gray matter. DTI has been thoroughly studied and histologically validated in the WM^11,12^, and is the initial stepping stone for studying WM in neurological and psychiatric conditions.

Creating a universal normative model for dMRI metrics is challenging due to the influence of scanning protocol parameters, such as voxel size, the number of diffusion gradient directions, *b*-values (diffusion weightings), and field strengths^13–16^. Here we address this challenge using hierarchical Bayesian regression (HBR)^17^, an approach designed to model site, scanner and cohort effects in multi-site data. To incorporate sufficient dMRI data for NM, we pooled dMRI data from diverse international studies, accommodating data collected with different acquisition protocols. As some datasets focus on development and others on aging, we merged data from different age ranges to build a normative model covering the lifespan (4-91 years).

We built normative models for the most widely used dMRI brain metrics – fractional anisotropy (FA), and mean, radial, and axial diffusivity (MD/RD/AD) – based on 54,583 individuals scanned in 19 public neuroimaging studies. We then leveraged the estimated normative models for 21 deep WM regions to identify WM abnormalities in two age-related conditions: dementia, mild cognitive impairment (MCI) for senescence phase and a neurogenetic disorder, 22q11.2 deletion syndrome (22qDel) for the development phase. In this anomaly detection scenario, we first show how the deviation scores from the reference population norm can identify abnormal microstructure in mild cognitive impairment (MCI), dementia, and 22qDel.

Lastly, to better understand the lifespan charts, we tested two hypotheses proposed by the theory of retrogenesis^18,19^. First, we tested the so-called “last-in first-out” hypothesis, which suggests that the early-maturing cerebral structures are somewhat spared from neurodegeneration in old age and tend to degenerate last in late life, relative to late-maturing structures. By contrast, the “gain-predicts-loss” hypothesis argues that the neural tissues tend to show correlations between their rate of tissue maturation in early life and degeneration in later life, with fast developing structures showing the most rapid decline.

## 2. Results

### 2.1. Lifespan trajectories of brain microstructural metrics

We fitted normative models to four diffusion tensor imaging (DTI) metrics (FA, MD, AD, RD) from 21 deep WM regions and the Global-WM (a single metric including the entire WM) based on 19 worldwide cohorts after rigorous quality control (see **Table S1**). Normative models were estimated for 22 WM regions for each DTI metric using a generative model that uses 10 repetitions of 80%-20% train-test splits –with different random seeds– of all datasets but excluding the clinical population. **Figures 1 and 2** show the results of these sampling procedures for the Global-WM. The posterior distributions for the parameters of the HBR model are estimated from the training data using the No-U-Turn Sampler (NUTS)^20^. For inference on the test data, we sample from the posterior predictive distribution, which represents the distribution over new observations given the estimated parameters. To derive the normative ranges, we sampled from the posterior predictive distribution across the full age range for both males and females (see Methods section). In data from 19 sites, we see that each site has a different intercept (**Figure 1B**). This is expected, as the mean values of DTI metrics depend on the scanning parameters, such as voxel size and *b*-value, yielding a different distribution of diffusivity and anisotropy values per site. In **Figure 1D** we show the 2nd, 10th, 25th, 75th and 90th and 98th percentile curves, corresponding to Z=土0.68, 土1.29 and 土2, respectively. For the Global-WM, the FA showed an inverted U-shaped trajectory, with a peak at 29 years of age. The three diffusivity measures (MD, AD, and RD) show a U-shaped trajectory, with minima later in life: at 37 years for RD, 43 years for MD, and 53 years for AD (**Figure 2**). **Figures S1**, **S2**, **S3** and **S4** show the lifespan trajectories and percentile curves for each DTI metric for the 21 regions besides the Global-WM.

**Figure 1.**
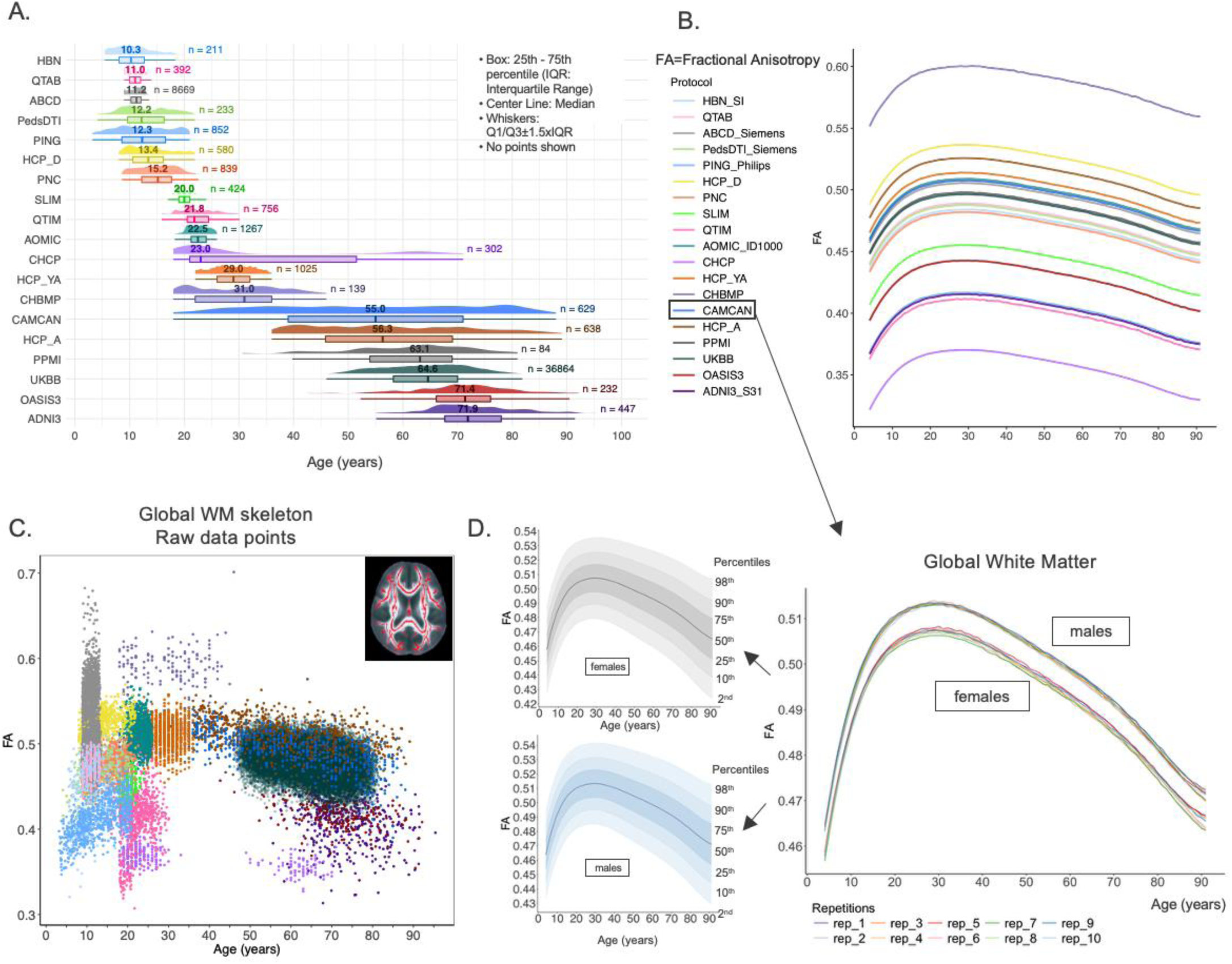
Normative models of brain microstructure. **A)** Age distribution of the 19 datasets used in the experiments. Median ages are shown above the boxplots, and total numbers of subjects are listed to the right. For clinical samples (HBN, ADNI3, OASIS3 and PPMI), only the control subjects are shown (as they were the only ones used to define the normative model). The color code for the 19 datasets is the same for panels A, B and C. **B)** Shows the lifespan trajectory for the global white matter fractional anisotropy metric (Global-WM FA) as a mean across males and females (these are separated in D). Each curve shows the result of generating synthetic predictions for each of the 19 scanning protocols by sampling from the predictive posterior distribution. Thus, each curve has a different intercept (random intercept option used for the hierarchical Bayesian Regression). **Note:** HBN_SI, ABCD_Siemens, PedsDTI_Siemens, PING_Philips, AOMIC_ID1000 and ADNI3_S31 are subsamples of HBN, ABCD, PedsDTI, PING, AOMIC and ADNI3, respectively. **C)** shows the raw data points of FA (Fractional Anisotropy) for the Global-WM,where each cohort is shown in a different color. **D)** shows the generated predictions for the site CAMCAN for the Global-WM FA (Fractional Anisotropy) metric, for each of the 10 HBR iterations. The reference charts can be made separately for women and for men by modeling sex as a batch effect. To the left, the lifespan trajectory of FA (Fractional Anisotropy) for the Global-WM after averaging the 10 iterations for each sex separately. It also shows centile curves for Z=土0.68, 土1.29 and 土2. Note: When generating data from one site and for all 10 repetitions, we noticed that for ages greater than 91 years and under 4 years there is instability across the repetitions, likely caused by the scarcity of subjects in those age ranges. We show the main trajectory of FA (Fractional Anisotropy) between ages 4 and 91 years. Source data are provided as a Source Data file.

**Figure 2.**
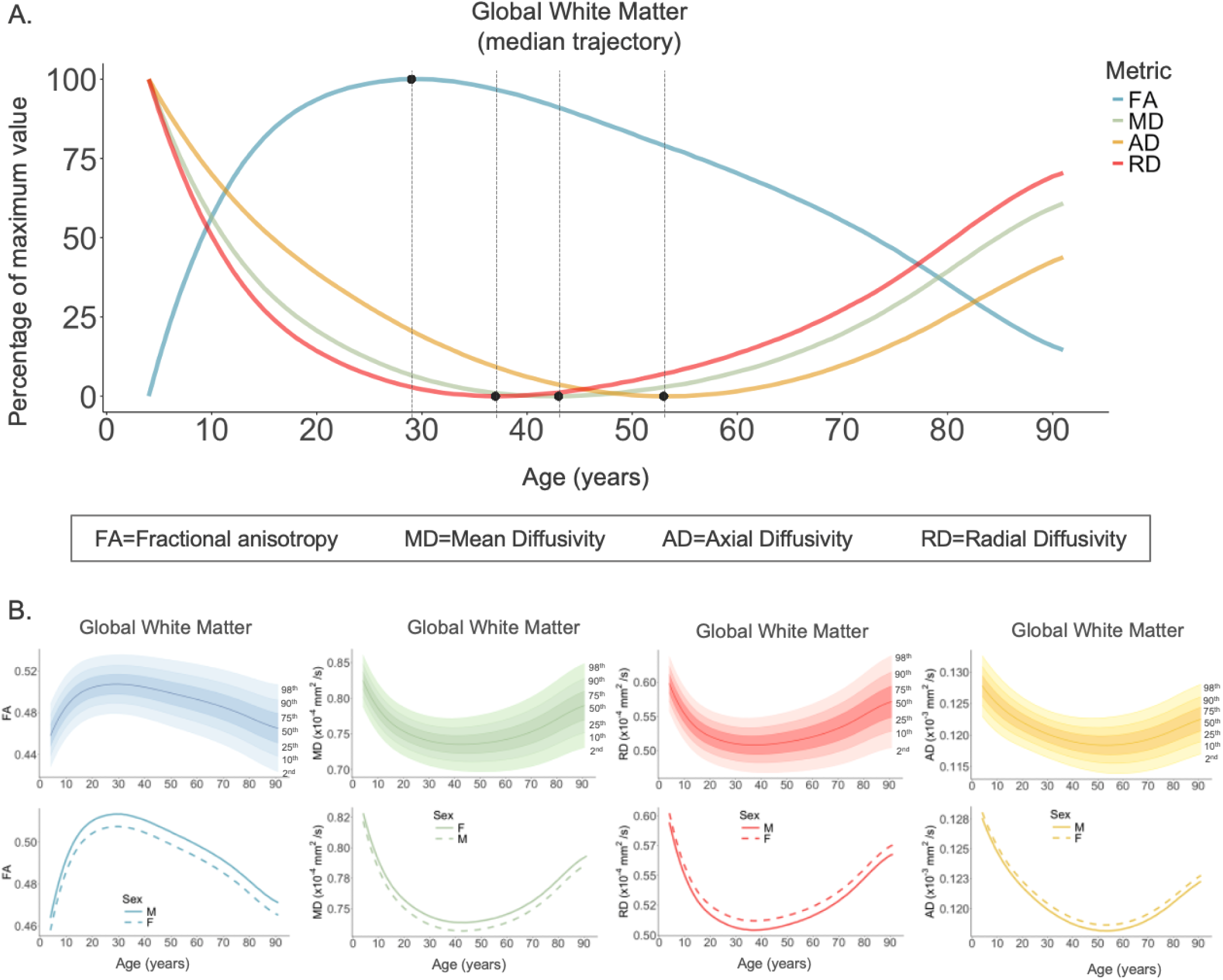
Normative models of the overall white matter (Global-WM) and peak ages per ROI. **A)** Age-dependent charts of the four DTI metrics (FA, MD, AD, RD) across the lifespan (age 4-91 years). In order to fit them in the same plot we adjusted the y-axis as a percentage of the maximum value for each metric. The dashed vertical lines indicate the age of maximum FA and the minimum age for MD, AD, RD. **B)** We show the centile curves for FA, MD, AD and RD for the Global-WM (females). The bottom row shows the trajectories for males and females separately. Source data are provided as a Source Data file.

For those regions that had a U-shape trajectory, FA peaked between 16 years (GCC) and 39 years (FX), MD reached minima between age 30 (PTR) and 51 years (UNC), while RD showed a pattern similar to that of MD with minima as early as 27 years (PTR) to 44 years (UNC). AD showed the greatest variability across the regions among all metrics; 14 out of 22 regions showed a U-shaped pattern for which minimum AD values occurred between 42 and 55 years. For AD, some regions had a quasi-linear trajectory from ages 4 to 80 years (CGH, FXST), and three showed an initial increase up to age ∼20 to reach a plateau or later increase again at a later age (ALIC, CGC, CST). The average age of maturation (peak/minimum) of the DTI metrics differed. Pairwise comparisons of peak/minimum ages across regions were significantly different: FA to MD (ψ=-13.0, p(corrected)=0.01, 95% CI=[-17.3, –9.1]), FA to AD (ψ=-20.1, p(corrected)=0.01, 95% CI=[-27.1, – 10.1]), FA to RD (ψ=-7.1, p(corrected)=0.01, 95% CI=[-11.2, –4.9]), MD to AD (ψ=-6.8, p(corrected)=0.05, 95% CI=[-10.1, –3.1]), MD to RD (ψ=5.1, p(corrected)=0.008, 95% CI=[3.6, 6.7]), AD to RD (ψ=12.6, p(corrected)=0.02, 95% CI=[8.1, 16.6]) (**Table S2**). On average, FA reached maturity at the youngest age (trimmed mean=29.2 years), followed in order by RD (trimmed mean=35.8 years), MD (trimmed mean=41.7 years) and AD (trimmed mean=48.2 years). **Figure 3** shows the peak/minimum ages for FA, MD, AD, and RD for 21 WM regions.

**Figure 3.**
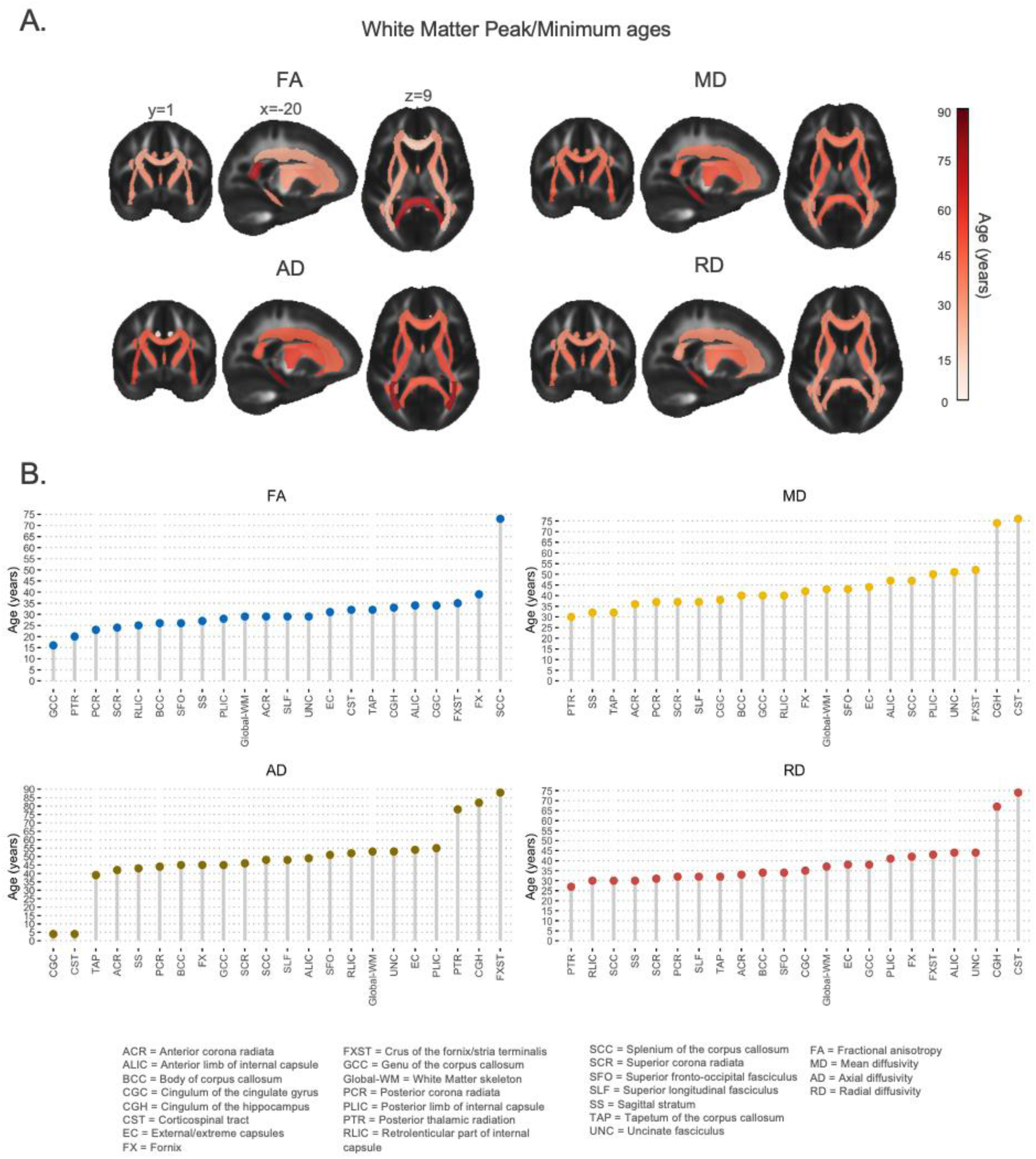
White matter peak/minimum ages. A) Peak ages for the 21 white matter regions of interest for all four DTI metrics. B) Age at peak or minimum for each DTI metric, organized from youngest to oldest age of maturation. Source data are provided as a Source Data file.

A key question for the practical utility of normative models is how well the models fit the data. To test model accuracy, we ran three diagnostic metrics for all 22 WM regions: the standardized mean squared error (SMSE) that assesses the goodness of fit and central tendency, the mean standardized log loss (MSLL)^20^ that evaluates both the mean and standard deviation of the predicted distribution, and Pearson’s correlation (Rho) between the observed and the predicted values. An SMSE value lower than 1 indicates a better fit, a more negative MSLL value and a Rho value closer to 1 indicate a better performance. SMSE scores across all regions and across all DTI metrics were below 0.78 (median=0.57; sd=0.11) and MSLL scores were below –0.14 (median=-0.31; sd=0.09), and Rho ranged from 0.46 to 0.86 (median=0.64; sd=0.08). Consequently, an SMSE value of 0.5 means that the model’s predictions are better than simply predicting the mean and that the hierarchical structure of HBR is helping the model generalize better than a non-hierarchical or naive model. An MSLL below zero (0) indicates that the model performs better than the baseline and Rho above 0.5 indicates a moderate to strong linear relationship between the model’s predictions and the actual outcomes. This, in the context of HBR, is a good sign that the partial pooling and group-level structure are helping the model generalize well across different levels of the hierarchy. AD had on average the best SMSE, MSLL and Rho scores (medians=0.47, –0.39, 0.72, respectively), followed, in order, by FA, MD and RD. In terms of the WM regions, the Global-WM had the best SMSE, MSLL and Rho scores for FA and AD, whereas the fornix (FX) had the best scores for MD and RD. A few regions stand out with lower performances, i.e., the tapetum (TAP) (for FA and RD), the corticospinal tract CST (for MD and AD), the sagittal stratum (SS) (for MD and RD) and the uncinate fasciculus (UNC) (for AD). See results in **Figure 4B** and Figures **S5-8**. **Figures S9-12** show the diagnostic metrics by site. Importantly, we emphasize that these metrics do not necessarily have to be consistent across different sites. For instance, when comparing UKBB and ABCD, Rho is higher in UKBB due to a greater variance to account for, given that the age range spans about 30 years (50 to 80 years). In contrast, in ABCD, explaining as much variance is nearly impossible since the age range is much smaller (8 to 13 years). Essentially, these metrics are not comparable across sites unless there is certainty that they originate from the same distribution, which is not the case. In none of the cases did we find any significant differences between the diagnostic scores (SMSE, Rho, MSLL) from the HBR model from the current manuscript and the HBR models with prior ComBat harmonization (see **Tables S3-S4** for details). Lastly, we did not find evidence for residual site effects on the Z-scores of the four metrics (FA, MD, RD, AD) (see **Tables S5-S8**).

**Figure 4.**
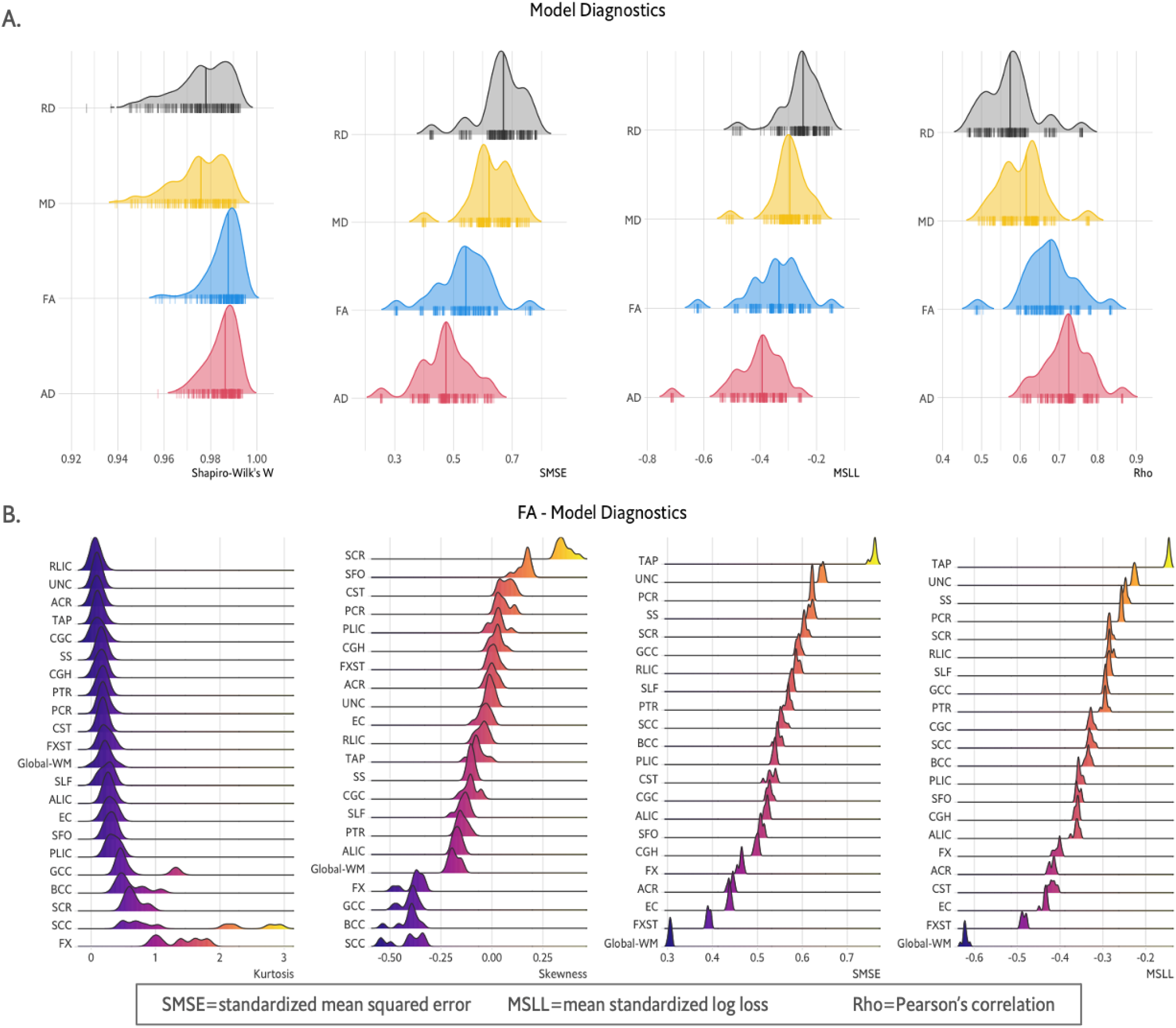
Hierarchical Bayesian Regression: model diagnostics. Here we show how well the models fit the data, as a function of age. **A)** We show Shapiro-Wilk’s W statistic, the standardized mean squared error (SMSE), the mean standardized log loss (MSLL) and Pearson’s correlation (Rho) between the observed and the fitted values for all four DTI metrics. All WM regions are lumped into one density plot for each DTI metric. The line inside the density plot indicates the median. Axial diffusivity (AD) had on average the lowest SMSE (median=0.47) and MSLL (median=-0.39) and highest Rho (median=0.72) of all DTI metrics. RD had on average the highest SMSE (median=0.67) and MSLL (median=-0.24) and lowest Rho (median=0.57). FA had the highest W (median=0.98) and MD the lowest (median=0.97). **B)** We show excess kurtosis, skewness, SMSE, MSLL per ROI for fractional anisotropy (FA). Skewness and kurtosis are shown to contextualize the results of the Shapiro-Wilk normality test. We show that most regions had a mesokurtic distribution (kurtosis < 1.0) with the exception of the FX and the SCC. Most regions had a skewness between –0.25 and 0.25, except for the SCR (positively skewed) and the SCC, BCC, GCC and FX (negatively skewed). The regions with the best scores for SMSE and MSLL are the Global-WM, the FXST and EC. ROI-wise plots for the other metrics (MD, AD, RD) are found in the supplements (Fig. S6, S7, S8). Source data are provided as a Source Data file.

Additionally, to test the reliability of the estimated centiles we calculated Shapiro-Wilk’s W, the kurtosis and the skewness of the estimated Z-scores (**Figure 4**). We verified how well the estimated centiles followed the normal distribution. All W scores across all four DTI metrics were above 0.97 – here 1.0 is a perfect score and indicative of normality. Furthermore, we investigated the distribution of the estimated Z-scores. In **Figure 4B**, the lowest scoring WM regions for kurtosis, from the RLIC to the PLIC, all 18 regions have excess kurtosis values between 0 and 1 (mesokurtic). The FX showed higher kurtosis (between 1 and 10) for all four metrics (FA, MD, AD and RD), and the TAP showed values between 1 and 2 for MD and AD. This may be due to the low SNR of those regions, caused by partial voluming with CSF-filled structures in the neighborhood (i.e., the lateral ventricles), and by the misalignments during registration caused by the low anatomical resolution around those regions. Interestingly, the FX also showed high skewness across DTI metrics, negative for FA, positive for the MD, RD and AD, despite showing the best goodness of fit metrics for MD and RD. The TAP showed high positive skewness for MD, RD, AD, following the FX (see **Figures S5-8, S13-16**). Finally, we were not able to predict the site from the Z-scores using a leave-one-out strategy with Support Vector Machines, assessed by using accuracy, balanced accuracy and macro F1. This suggests that the deviation Z-scores derived from our original HBR model are handling unwanted site effects as effectively as a prior ComBat harmonization before running HBR (see Methods section).

### 2.2. Retrogenesis hypotheses

We did not find evidence for the ‘gain-predicts-loss’ hypothesis (**Figures S17, S18**). We found evidence supporting the ‘last-in, first-out’ hypothesis, for FA, MD and AD, but not RD; we found significant correlations between the age of peak and the percentage of change post-maturation (**Figure 5**) —85-91 years— for FA (*r*=0.21, *p_spin*=0.02, α=0.05), —75-84 years— for MD (*r*=0.42, *p_spin*=0.04, α=0.05) and AD (*r*=0.24, *p_spin*=0.02, α=0.05), suggesting that WM tracts with a later age of maturation decline faster (**Figure S19-S22**). We also found a positive association between the percent change from age 4 to 14 years (**Figure 5**) and the age of peak for FA —maximum FA—(*r*=0.03, *p_spin*=0.02, α=0.05) and MD —minimum MD— (*r*=0.32, *p_spin*=0.01, α=0.05). That is, the greater the early percent change from age 4 to 14 years (**S19, S20**), the younger the age of maturity. Our findings suggest that for FA and MD, those WM regions that develop faster reach maturity earlier and decline slower, whereas those regions with a slower development, tend to peak later and decline faster. Finally, we did not find differences in the age of peak between the three groups of WM regions: commissural, association and projection fibers.

**Figure 5.**
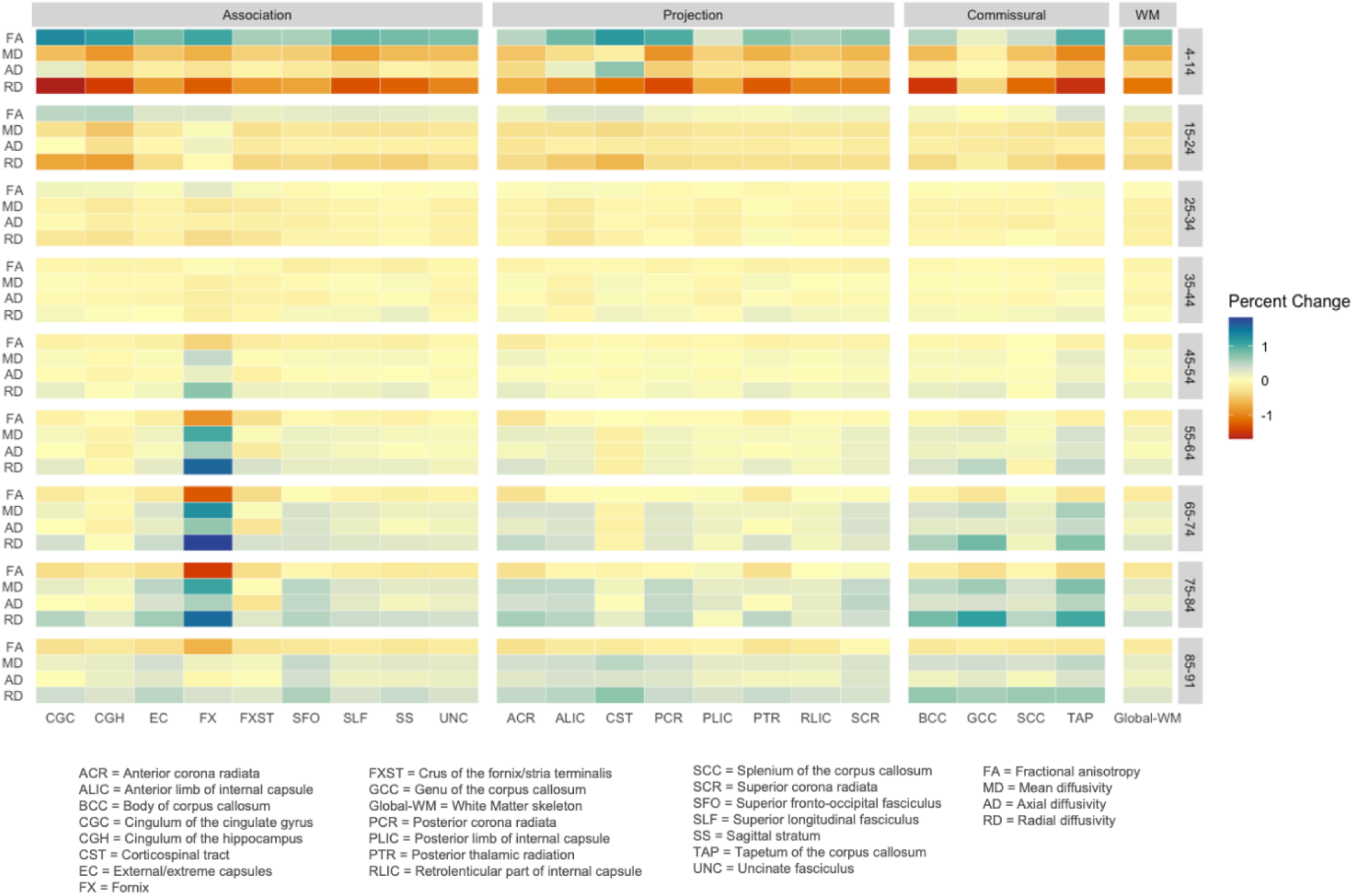
White matter changes across the lifespan. Percent change of DTI metrics every 10 years across the lifespan (4-91 years). regions are grouped by type: association, projection, and commissural fibers and Global-WM. Source data are provided as a Source Data file.

### 2.3. Anomaly detection

As detailed in the Methods, we next set out to find significant deviations from the reference model for each ROI per metric across 10 experimental repetitions. We did this for two datasets, for which 80% of healthy controls were used in the model estimation, i.e., ADNI3 and OASIS3. The anomaly detection focused on the participants with dementia or MCI. The Z-scores were first converted to abnormality scores, i.e., the probability of each metric being normal, based on a modified normal cumulative distribution function^17^. We use these predicted probabilities of abnormalities together with an ROC analysis to identify the DTI metrics and WM regions that are most informative for the brain disease under study. We calculated the area under the ROC curve (AUC) for classification of the patients (AlzD or MCI) versus the 20% controls in the test set for each of the 10 experimental repetitions. Here, we report the regions per metric with significantly above the chance performance (AUC>0.5) that passed multiple comparisons correction (across metrics) in 9 out of 10 repetitions to consider deviations significant and stable; for dementia, we found the highest classification accuracy for the CGH for RD (AUC=0.74), for MD (AUC=0.7) and FA (AUC=0.65). For axial diffusivity in dementia, the BCC had the highest accuracy (AUC=0.67). For MCI, the AUCs were lower, as expected, than in dementia: values ranged from 0.55 to 0.6 (**Tables S9 & S10**). The best performance was observed for MD in the SCC (AUC=0.6), for AD in the BCC (AUC=0.67), and in the FX for FA and MD (AUC=0.58/0.59, respectively). Extreme deviations (|Z|>2) were mostly in the same direction for dementia and MCI; most subjects showed positive deviations (Z>2) for the diffusivity measures (MD, RD, AD) and negative deviations for FA (**Tables S11 & S12**). In MCI and dementia, the metric with the largest number of extreme deviations was RD (**Figure 6**). 37% of dementia participants had positive extreme deviations for RD in the CGH, and 25% in the SCC; 17% of subjects with MCI had positive extreme deviations in the SCC and 11% of subjects in the CGH. For FA, negative extreme deviations were mostly found in the CGH for dementia (20%) and the FX (21%) and in the SCC for MCI (13%).

**Figure 6.**
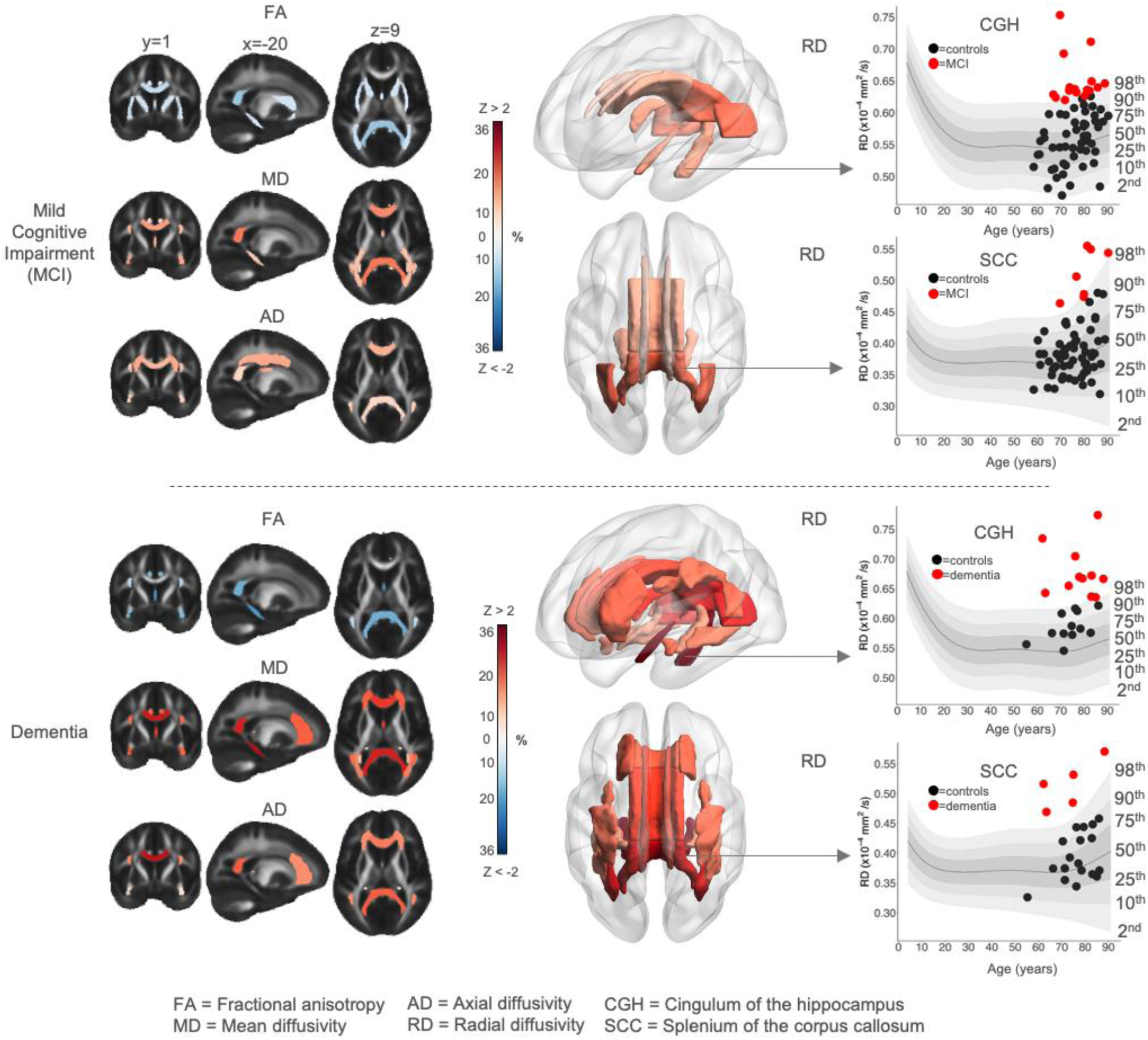
Anomaly detection in elderly individuals with mild cognitive impairment (MCI) and dementia (ADNI3 and OASIS3 datasets). The WM regions shown have AUC > 0.5 (i.e., the models in these regions significantly discriminate the cases from controls) and passed multiple comparisons testing while controlling the false discovery rate (FDR). The extreme deviations (|Z|>2) are shown for FA, MD, AD in 2D, and for RD in 3D. The regions are color-coded based on the percentage of participants with extreme deviations –-blue scale for negative deviations, red scale for positive deviations. The metric with the greatest proportion of extreme deviations is RD, with up to 36% of positive extreme deviations for the **CGH** in dementia patients. As expected, the extreme deviations for FA are in the opposite direction than for diffusivity metrics, i.e., MD, AD, RD. The centile curves on the right are specifically for the ADNI3-S55 dataset. Dots displayed on the centile curves denote the ADNI3-S55 patients. The dots in red are those with Z>2. **CGH**=cingulum of the hippocampus; **SCC**=splenium of the corpus callosum. 3D visualizations were done with BrainNet Viewer^85^. Source data are provided as a Source Data file.

### 2.4. Model adaptation

Next, we adapted the large-scale trained reference model to healthy controls from two new independent datasets. We assume that these two datasets were not available during the development phase of the reference model to simulate a real-world scenario where the model needs to be adapted to local data in the deployment stage. The NIMHANS elderly cohort has MCI and dementia participants and age-matched healthy controls; and the UCLA neurogenetic dataset has 22qDel participants and age-matched healthy controls (**Table 1**). For each dataset we used 50% of the healthy controls for recalibrating the model. Thereafter, the remaining 50% of healthy controls and the patients were used for anomaly detection, as described in the Methods (sections 9 and 10). For the NIMHANS cohort, we found a greater severity –assessed by the percentage of subjects with extreme deviations– for MCI compared to dementia (**Figure 7A & 7B**). Additionally, for MCI we found subjects with extreme positive deviations for MD, AD and RD, whereas for dementia, all four DTI metrics showed regions with significant extreme deviations; negative deviations for FA and positive deviations for diffusivities (**Tables S13-16).**

**Figure 7.**
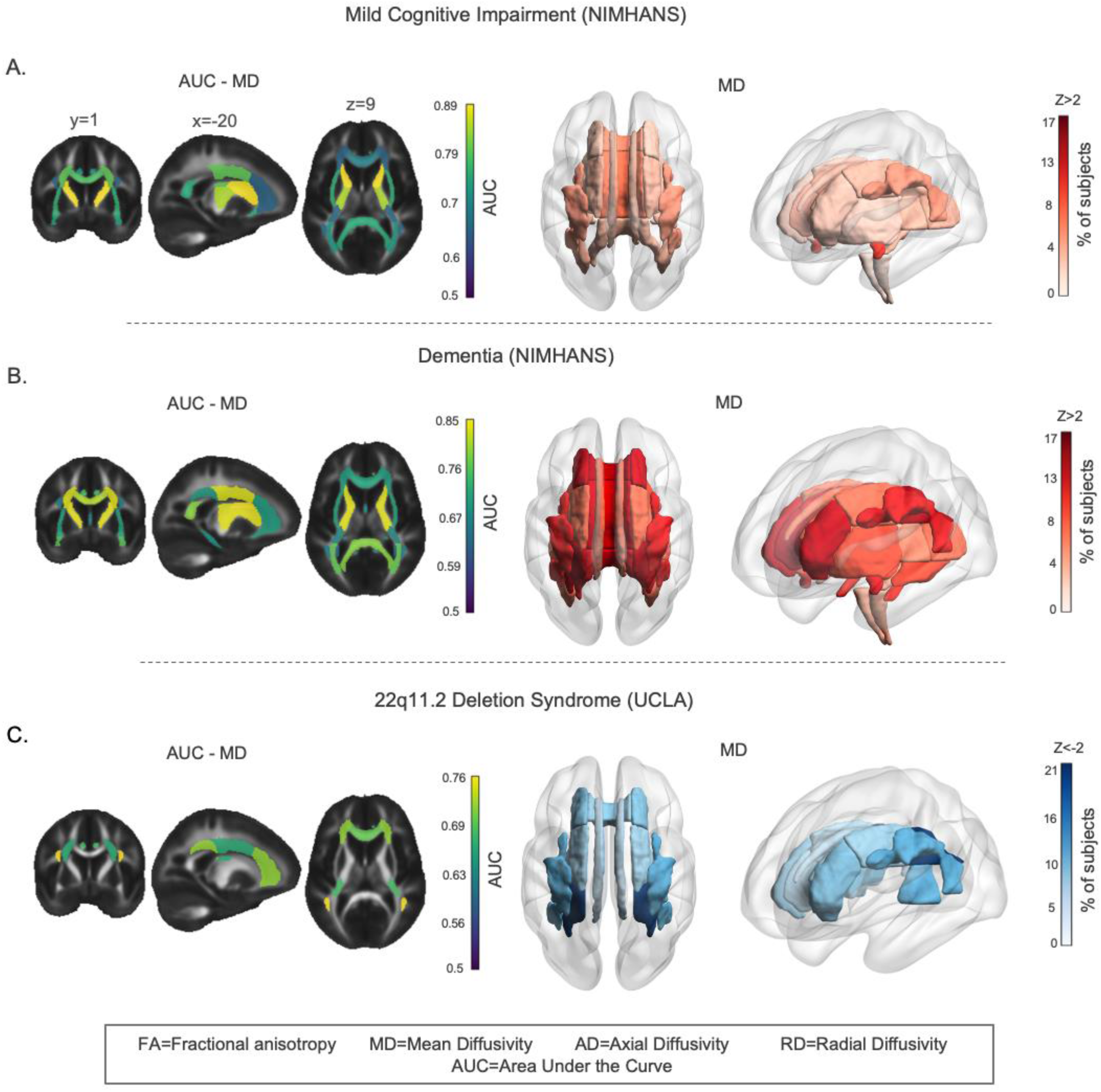
Results of the model adaptation experiment. Anomaly detection for patients with Mild Cognitive Impairment (MCI) and dementia in the NIMHANS dataset and patients with 22q11.2 Deletion Syndrome (22qDel) in the UCLA dataset. The colored ROIs are those that passed FDR correction, with AUC>0.5 when tested against the test sample of healthy controls (50%) from the same dataset (UCLA or NIMHANS). We show the AUCs per ROI and the percentage of extreme deviations for MD (Z>2 & Z<-2). **A)** Discriminative results for the MCI patients from the NIMHANS dataset. **B)** Discriminative ROIs for the dementia patients from the NIMHANS dataset. **C)** Discriminative ROIs for 22qDel. 3D visualizations were done with BrainNet Viewer^85^. Source data are provided as a Source Data file.

**Table 1.** Demographics (age and sex) of the datasets used for hierarchical Bayesian regression (HBR) model estimation and model adaptation (NIMHANS and UCLA). The N for each group reflects the participants after quality control and removal of outliers, as explained in the methods section. The hyphen (-) in the Controls/Cases column indicates that the study was population-based. (SD=standard deviation). MCI=Mild Cognitive Impairment.

| Study/Dataset | Controls/<br>Cases | Age Range | Median Age (SD) | N (Males %) | Country |
| --- | --- | --- | --- | --- | --- |
| ABCD | - | 8 - 13 | 11.2 (1.1) | 8669 (52) | United States |
| AOMIC | - | 18 - 26 | 22.5 (1.7) | 1267 (46) | The Netherlands |
| CAMCAN | - | 18 - 88 | 55 (18.5) | 629 (49) | United Kingdom |
| CHBMP | - | 18 - 46 | 31 (7.7) | 139 (89) | Cuba |
| CHCP | - | 18 - 71 | 23 (17.7) | 302 (44) | China |
| HCP-A | - | 36 - 89 | 56.3 (13.9) | 638 (42) | United States |
| HCP-D | - | 8 - 21 | 13.4 (3.6) | 580 (48) | United States |
| HCP-YA | - | 22 - 36 | 29 (3.6) | 1025 (45) | United States |
| PedsDTI | - | 4 - 21 | 12.2 (4.2) | 233 (48) | United States |
| PING | - | 3 - 21 | 12.3 (4.8) | 852 (52) | United States |
| PNC | - | 8 - 22 | 15.2 (3.3) | 839 (45) | United States |
| QTAB | - | 9 - 14 | 11 (1.2) | 392 (51) | Australia |
| QTIM | - | 15 - 30 | 21.8 (2.9) | 756 (37) | Australia |
| SLIM | - | 17 - 24 | 20 (1.0) | 424 (44) | China |
| UKBB | - | 46 - 81 | 64.6 (7.5) | 36864 (46) | United Kingdom |
| ADNI3 | Controls | 55 - 95 | 71.9 (7.5) | 447 (41) | United States |
|  | MCI | 55 - 93 | 75.4 (8.0) | 214 (57) |  |
|  | Dementia | 55 - 95 | 78 (8.8) | 69 (59) |  |
| HBN | Controls | 5 - 19 | 10.3 (3.6) | 211 (53) | United States |
| OASIS3 | Controls | 42 - 92 | 71.4 (8.4) | 232 (48) | United States |
|  | MCI | 62 - 87 | 76.8 (7.5) | 11 (36) |  |
|  | Dementia | 65 - 82 | 76.1 (5.1) | 12 (41) |  |
| PPMI | Controls | 30 - 80 | 63.1 (10.5) | 84 (63) | United States |
| NIMHANS | Controls | 51 - 81 | 65.0 (7.5) | 118 (50) | India |
|  | MCI | 52 - 88 | 68.0 (7.4) | 89 (65) |  |
|  | Dementia | 51 - 86 | 70.5 (7.4) | 90 (58) |  |
| UCLA | Controls | 7 - 31 | 14.2 (5.2) | 67 (47) | United States |
|  | 22q11.2 Del. S. | 6 - 27 | 14.0 (6.1) | 80 (46) |  |

For 22qDel, we found significant effects in multiple regions for MD and AD. MD showed more regions with extreme deviations (9 of 21 regions) than AD (5 of 21 regions). Interestingly, the entire corona radiata showed negative extreme deviations (ACR, SCR, PCR). For both DTI metrics, extreme deviations were negative, except for the PLIC in AD, where the extreme deviations were positive. (**Figure 7C; Tables S17 & S18**).

## 3. Discussion

We used HBR to create normative models of the brain’s WM microstructure across the lifespan, leveraging a diverse, large-scale international dataset of diffusion MRI scans. In addition to revealing lifespan trends for the most widely used metrics of brain microstructure, we showed how the model is useful for investigating neurological, and neurogenetic conditions. Reference models were created to quantify the distributional properties of FA, MD, AD, and RD throughout life. By mapping their normal range of variation, and projecting clinical data into this normative range, we detected patterns of deviation from this range for dementia, MCI, and 22qDel. Such large-scale normative models offer transdiagnostic clinical value; unlike group differences in case-control analyses, normative modeling yields individual profiles of anomalies, which are agnostic to the clinical labels and accommodate the well-supported premise that not all individuals with the same disease deviate in the same brain regions and in the same direction^1,3^. Mapping the extent and scope of individual-level deviations can complement the aggregated group difference effects reported in large-scale case-control studies, but may also flag anomaly patterns that would be washed out in group comparisons of cases and controls. Prior work on normative modeling of diffusion MRI metrics has been done by Chamberland et al.^21^, Elad et al.^22^, and Cirstian et al.^23^. Chamberland et al. included not just DTI metrics but also higher order metrics. They used bundles extracted from tractograms and used a rather small sample of subjects.

As expected, for most regions, we found an inverted U-shaped lifespan trajectory for FA, increasing during maturation and declining after a peak. Diffusivity metrics exhibited a U-shape lifespan trajectory, with an initial decrease during maturation and an increase later in life. FA typically reached maturity (i.e., a maximum) at a younger age, followed by RD and MD, whereas AD reached maturity (i.e., a minimum) at age ∼40 or ∼50 years, depending on the ROI. This is in line with prior cross-sectional and longitudinal studies of age-dependent changes in WM DTI metrics^24–30^. Even though most regions in this study showed a U-shaped or inverted U-shaped trajectory across the lifespan, particularly for FA, MD, and RD, this was not the case for all WM regions. AD was the metric with the most regions deviating from this U-shaped trajectory–six out of 22 regions–with some regions showing a generally linear trend, followed by FA, whereas RD and MD were the most consistent in terms of displaying the inverted U-shape. Future normative modeling efforts focusing on these few WM regions may consider fitting age with linear models.

Relatively few studies have tested the retrogenesis hypothesis on WM DTI metrics^18,19,28,31–34^, but such an analysis is now feasible as large dMRI datasets are available from childhood to senescence. Retrogenesis was first proposed in the context of Alzheimer’s disease, where functional and cognitive decline followed a pattern that was to some extent the reverse of the normal human developmental sequence^35^. This hypothesis has been termed ‘last-in first-out’. It was later hypothesized that structural and functional neuroimaging phenotypes might follow the same ‘last-in first-out’ trend^36^, with some evidence supporting this for WM DTI metrics in Alzheimer’s disease^37,38^. The underlying hypothesis for the WM is that the early myelinating fibers, such as the internal capsules, cerebral peduncles, and corticospinal tracts, may be less vulnerable to degeneration than the late myelinating fibers, such as the inferior and superior longitudinal fasciculi, anterior centrum semiovale, corpus callosum, and hippocampus. Nonetheless, the empirical evidence for this theory for DTI metrics has been moderate^33^ or absent^18,19,25^. Instead, we found significant correlations between the age at maturation for FA and the percentage of change post-maturation (85-91 years), and the age at maturation for MD&AD and the post maturation change (75-84 years), suggesting that WM tracts with a later age of maturation may decline faster. We also found a positive association between the percent change of FA and MD from age 4 to 14 years and the age of peak (i.e., the maximum FA and the minimum MD) across regions, suggesting that the fastest and earliest maturing regions may have a less pronounced slope from peak to age 91 years and thus effectively degenerate slower. Nevertheless, approaches using piecewise linear fits with three defined sections of the lifespan: maturation, plateau and degeneration^18^, may be of interest for future experiments.

Another related retrogenesis theory is the ‘gain-predicts-loss’ hypothesis. It suggests that the rate of tissue maturation during development is inversely correlated with the rate of degeneration at an older age. This hypothesis has been tested on lifespan samples using either quadratic fits or by correlating the pre-maturation slope to the post-maturation slope, assuming some degree of symmetry on either side of the maturation peak. Even so, some studies found no such symmetry in DTI lifespan trajectories^18,34^. Others have found moderate or strong evidence^28,32^, especially for FA and RD. We did not find sufficient evidence for the ‘gain-predicts-loss’ hypothesis for any of the DTI metrics.

The post-analysis of the anomaly detection experiments identified extreme deviations (|Z|>2) in both directions, positive and negative–a hallmark of heterogeneity within the clinical group. In MCI and dementia, most deviations for MD and RD were positive, indicating higher WM diffusivity; however, for some regions, a smaller set of participants showed negative deviations (lower diffusivity). This was more prevalent in MCI than in dementia, and more prevalent for FA and AD than for MD and RD. FA may indeed be higher in the early stages of MCI than in healthy controls, and may even reflect recent evidence of Alzheimer’s disease subtypes: typical, limbic-predominant, and hippocampal-sparing Alzheimer’s disease^39^. In the future, normative models may help to identify the disease subtype to which an individual belongs to, given the availability of other biomarkers such as tau, amyloid, and APOE, although this is beyond the scope of the current manuscript.

Our anomaly detection framework was also able to gauge disease severity in new subjects by adapting the normative models to an unseen sample. For the NIMHANS dataset, we found some WM regions with extreme deviations in MCI, whereas all regions had significantly anomalous deviations for dementia, especially for MD. The observed differences in the MCI and dementia patterns between the ADNI3/OASIS3 and NIMHANS samples may be partially explained by the differences in ancestry (predominantly white Caucasian North Americans in the ADNI3/OASIS3 datasets vs. Indian ancestry participants in the NIMHANS dataset), recruitment protocols or sample sizes.

We also found a similar phenomenon for 22qDel: for some regions, a proportion of 22qDel participants also had deviations in the opposite direction to the majority of subjects; this aspect would be missed using approaches that averaged multi-subject data prior to group comparisons. In particular, for 22qDel, even though the largest case-control studies show lower diffusivity for almost the entire WM^40^, we identified some regions for which a small proportion of extreme deviations was in the opposite direction–that is, higher diffusivity (MD and AD). This may reflect biological differences within the sample of cases, such as the length of the 22q11.2 locus deletion. Another reason may be that many factors influence DTI, such as neurite density, axonal dispersion, and crossing fibers; higher-order diffusion models and advanced data acquisitions may be needed to disentangle these effects.

The question remains about the generalizability of the reference models across ethnoracial groups. We included samples from diverse countries for the training phase of the normative models (i.e., The Netherlands, the United Kingdom, Australia, the United States, China and Cuba). Additionally, we ran model adaptation to an independent dataset from India and a dataset from the United States, to understand whether or not the genetic ancestry affects the pretrained model’s performance. One strength of our study is the diverse origins of the datasets; we did not include the ethnoracial background in our normative model. A detailed analysis of the ethnoracial background of all the datasets included in our study would be challenging as many countries limit or even prohibit the collection of ethnoracial data. We could estimate the ethnoracial identity by the country of origin of each sample, but that would be an incomplete and inaccurate analysis. Country-level grouping may not adequately reflect genetic heterogeneity and to adequately address this, we ought to explicitly model ancestry or include genomic data when available. However, even if samples do record race or ethnicity, it is at best a poor proxy for ancestry. Even so, the mathematical principles of model adaptation, outlined in the current paper, mean that the model is adaptable to any new datasets across the world. This aspect of normative models has been studied by Rutherford et al.^41^ and Ge at al.^42^, where the authors concluded that regardless of the ethnoracial background, the performance of pretrained race-neutral normative models was high, with mean absolute errors being less than 10% across ethnoracial groups. Even if there were differences in the distributions of brain metrics at new sites and scanners, the model adaptation step is designed to handle it.

Regarding the adaptation of the normative models to unseen datasets during training (UCLA and NIMHANS), we included more than 25 control subjects per new site in order to accurately calibrate the model to the data from the new sites. This number has been shown to yield good performance metrics on the test sets by Gaiser et al.^43^. Accordingly, the number of controls used for model adaptation were: UCLA_Prisma=27, UCLA_Trio=36, NIMHANS_Siemens=88, NIMHANS_Philips=30. (see **Table S19** for specific acquisition parameters). These are average typical sample sizes for clinical studies, where different scanners are often used within one institution. Our approach allows for model adaptation in such extreme cases of small data as shown by Kia et al^17^.

One limitation of this study is that the ENIGMA-DTI protocol has not been optimized for very early childhood (3-5 years), so models based on infant data^8^ may be more appropriate for this age range. Additionally, some of the regions are noisier than others, which may lead to greater noise or measurement error for some regions due to registration misalignments, which are common in volumetric/skeleton based registrations. A future refinement, to address issues with tract-to-tract registration, is to develop normative models based on tractometry, and evaluate their improved calibration versus any increase in resources required to run such a method. Another source of variability between regions is the anatomy of the tract itself. Some tracts are thinner and/or shorter than others, which in turn leads to difficulties with the alignment to an atlas. Some tracts are very thin, such as the Fornix (FX) and the Tapetum (TAP), and short, like the Uncinate (UNC), which after skeletonization may be subject to more misalignments during registrations. For instance, the close anatomical neighborhood of many tracts, such as the FX, the TAP and much of the corpus callosum (splenium, body and genu) consists mostly of the lateral ventricles filled with cerebrospinal fluid (CSF). This proximity to large CSF-filled structures results in higher partial volume effects around the WM tracts, thus reducing the SNR for the affected tracts. This will consequently have an impact in any regression model trying to fit a model to the data points derived from those tracts. This could be avoided in future work by directly registering the WM bundles in the streamline space, as suggested by Chandio et al.^44^.

WM regions differ from each other in terms of their goodness of fit, primarily because they differ in the strength of association with age. Some have strong age related effects, whilst others have less effect, at least across the age range we measure. Even though some WM regions showed bad scores for metrics of goodness of fit, i.e., SMSE, MSLL and Rho, those same regions often showed good calibration metrics for the centiles. For instance, the TAP and UNC for FA had high SMSE, high MSLL, and low Rho, meaning that their goodness of fit was not optimal, but the calibration metrics of the Z-scores were within the average of all the other regions. Interestingly, the FX, and the corpus callosum (SCC, BCC, GCC) showed the worst metrics of Z-score calibration (Kurtosis, Skewness, and SW), but the goodness of fit scores are within normal average scores.

Future studies will include a richer set of microstructural measures, including Standard Model Imaging^45^ and Neurite Orientation Dispersion and Density Imaging^46^. Although our results presented are based on DTI, the reference models are still highly sensitive to WM microstructural deviations from the norm in psychiatric and neurological diseases. Additionally, the single-shell scanning protocols are, to this day, the most commonly available on most MRI scanners. More advanced multi-compartmental models beyond DTI will require multi-shell dMRI data, which is still limited in clinical populations, but worthy of examination in future. Finally, more advanced distributional models may also be beneficial for normative modeling, such as high-order non-Gaussian distributions modeled using nonlinear probability warping^47^, SHASH function models^48^ and high-dimensional variational autoencoders that can model the joint distribution of multiple features^49,50^.

## Methods

The study was approved by the University of Southern California IRB (Proposal #HS-24-00535). There was no human data collection as part of the study, only secondary analysis of deidentified anonymized data collected as part of prior approved studies.

### 1. Datasets analyzed

**Table S19** summarizes the diffusion MRI acquisition protocols for all datasets. We included the following public datasets with various age ranges to cover the lifespan: the Adolescent Brain Cognitive Development (ABCD)^51^, the Amsterdam Open MRI Collection (AOMIC)^52^, the Cambridge Centre for Ageing and Neuroscience (CAMCAN)^53^, the Cuban Human Brain Mapping Project (CHBMP)^54^, the Chinese Human Connectome Project (CHCP)^55^, the Human Connectome Project Aging & Development (HCP_A & HCP_D)^56^, the Human Connectome Project Young Adult (HCP_YA)^57^, the DTI component of the NIH MRI Study of Normal Brain Development (PedsDTI)^58^, the Pediatric Imaging, Neurocognition, and Genetics data repository (PING)^59^, the Philadelphia Neurodevelopmental Cohort (PNC)^60^, the Queensland Twin Adolescent Brain (QTAB)^61^, the Queensland Twin Imaging dataset (QTIM)^62^, the Southwest University Longitudinal Imaging Multimodal (SLIM) study^63^, and the UK BioBank (UKBB)^64^.

Several clinical datasets were also included in the analyses: the third phase of the Alzheimer’s Disease Neuroimaging Initiative (ADNI3)^65,66^, the third release of the Open Access Series of Imaging Studies (OASIS3)^67^, the Parkinson’s Progression Markers Initiative (PPMI)^68^, and the Healthy Brain Network (HBN)^69^. Importantly, only the healthy controls (HC) of the latter four datasets were added to the pool of training data to build the normative models, yielding a total of 19 public datasets. **Figure 1A** summarizes the age distribution for the 19 datasets used for training. In contrast, the patients in ADNI3 and OASIS3 were included as test data for the anomaly detection experiment. ADNI3 and OASIS3 included patients with mild cognitive impairment (MCI) and dementia. Two additional datasets were used for the model adaptation experiments: the National Institute of Mental Health and Neuro Sciences (NIMHANS) dataset^70^ and the 22q11.2 copy number variant (CNV) dataset from the University of California Los Angeles (UCLA)^71,72^. **Table 1** summarizes the demographics of all datasets analyzed.

### 2. Preprocessing of diffusion MRI scans

Three studies provided preprocessed dMRI images through their official project websites: HCP_YA, ABCD and AOMIC. Two studies provided precomputed DTI maps (PedsDTI and UKBB). The remaining datasets were preprocessed in-house. In all cases, preprocessing included correction for eddy currents, movement, and EPI-induced susceptibility distortions, even though the protocols were not fully aligned. A detailed description of the in-house preprocessing performed on each dataset may be found in the Supplementary Methods section. Importantly, tensor fitting was performed on the datasets with single-shell dMRI data or on the lowest non-zero shell available, if multi-shell data were provided. DTI maps (FA, MD, RD and AD) were computed with DiPy’s tensor fitting function^73^ for the following datasets: AOMIC, CAMCAN, CHBMP, CHCP, HCP_A, HCP_D, HCP_YA, NIMHANS, PNC, QTAB and UCLA. FSL’s dtifit^74^ was used to fit the diffusion tensor for ABCD, ADNI3, HBN, OASIS3, PING, PPMI, QTIM, SLIM. All subjects’ FA maps were linearly registered with FSL’s flirt and nonlinearly registered to the ENIGMA-FA template with ANTs’ diffeomorphic non-linear registration algorithm^74,75^. For QTIM, non-linear registration was performed using the 3DMI algorithm^76^. The combined linear and non-linear transformations were applied to the MD, RD and AD maps. Mean DTI metrics were extracted from 21 bilateral regions of interest (regions) from the Johns Hopkins University WM atlas (JHU-WM)^77^ using the ENIGMA-DTI protocol^78^. The mean for each DTI metric across the whole skeleton (Global-WM) was also extracted for each participant, yielding a total of 22 regions.

### 3. Quality control

All study participants had information on age and sex at the time of the imaging session along with a usable dMRI scan. ADNI3, OASIS3, PPMI, CAMCAN and HBN were manually checked for head movement, FOV (field of view) related artifacts and other dMRI acquisition artifacts and preprocessing errors caused by poorly tuned software tools. For all other datasets used for training, outliers were detected with an anomaly detection algorithm called isolation forests^79^ implemented in Scikit-learn^80^, as detailed in the Supplementary Methods. Briefly, isolation forests were run on each ROI for each DTI metric, separately, to identify a subset of outlier subjects for each DTI metric. Subsequently, outlier subjects that were duplicated in at least two DTI metrics were removed from the sample. Aggregating the subjects left after this procedure plus the subjects from the datasets that were manually checked, a total of 54,583 subjects remained for training. The goal of using an automatic tool such as the isolation forests is to automate the quality check procedure and reduce the effects of inter-rater variability which arise when relying solely on manual QC. For a comparison between manual quality checked and automatic quality checked Z-scores see **Figures S23-S26** and Tables **S20-S23**. One may opt to train a set of initial normative models to find the outliers, remove them, and then retrain the normative models without the outliers^81^.

### 4. White matter regions assessed

Twenty-two JHU-WM regions of interest were assessed in this study (**Table S1**), including the following: ACR=anterior corona radiata, ALIC=anterior limb of internal capsule, BCC=body of corpus callosum, CGC=cingulum of the cingulate gyrus, CGH=cingulum of the hippocampus, CST=corticospinal tract, EC=external/extreme capsules, FX=fornix, FXST=crus of the fornix/stria terminalis, GCC=genu of the corpus callosum, PCR=posterior corona radiata, PLIC=posterior limb of internal capsule, PTR=posterior thalamic radiation, RLIC=retrolenticular part of internal capsule, SCC=splenium of the corpus callosum, SCR=superior corona radiata, SFO=superior fronto-occipital fasciculus, SLF=superior longitudinal fasciculus, SS=sagittal stratum, TAP=tapetum of the corpus callosum, UNC=uncinate fasciculus, and the Global-WM=mean across the entire WM skeleton of the ENIGMA-DTI template.

### 5. Normative Modeling with HBR

The HBR NM framework is thoroughly described by Kia et al.^17^, but here we summarize the basic principles.

Let 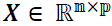 be a matrix of *n* the number of participants and *p* covariates. We added age and sex as the covariates for the present study and we modeled the non-linear effect of age with a cubic b-spline basis set expansion with five evenly spaced knots. Here, we denote the dependent variable as *y* ∈ ℝ. In its simplest form, NM assumes a Gaussian distribution over *y*, i.e., *y* ∼ *N*(μ, σ^2^), and it aims to find a parametric or non-parametric form for μ and σ given ***X***. In its parametric form, μ and σ are respectively parameterized on *f*_μ_(***X***, θ_μ_) and 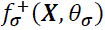, where θ_μ_ and θ_σ_ are the parameters of *f*_μ_ and 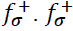 is a non-negative function used to fit the standard deviation of the noise ϵ_*i*_. Consequently, in a multi-site scenario, a separate set of model parameters could be estimated for each site, or batch, *i*, as follows:

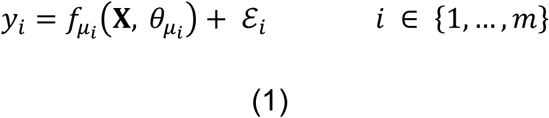

However, an assumption in HBR is that θ_μ*i*_ (and θ_σ*i*_) across different batches come from the same joint prior distribution that functions as a regularizer and prevents overfitting on small batches. HBR is described as a partial pooling strategy in the multi-site scenario, i.e., similar to the no-pooling scenario, in HBR, the parameters for *f*_μ*i*_ and 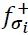 are estimated separately for each batch. Then, in the NM framework, the deviations from the norm can be quantified as Z-scores for each subject in the *i*_*th*_ batch:

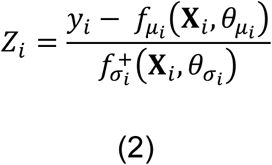

The harmonization of Z-scores happens at this stage where the Z-scores are adjusted for additive and multiplicative batch effects using the estimated *f*_μ*i*_ and 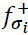 for each batch. In fact, the model does not yield a set of harmonized data, i.e., with the batch variability removed. Instead, it preserves the sources of biological variability that may correlate with the batch effects. Consequently, the frequent statistical dependence in multi-site neuroimaging projects that occurs between age and site (see **Figure 1A**), caused by different inclusion criteria across cohorts, may be better tackled with HBR. When removing the site effect with harmonization methods such as ComBat^82^ some of the biological variance correlated with age and site may be inadvertently removed^17^.

### 6. Regression models

We identified 37 dMRI acquisition protocols across the 19 datasets (see **Table S19**). Some of the 19 datasets had subsamples acquired at multiple locations, often at different time points, at scanners from different vendors, ultimately modifying the acquisition protocols and leading to differences in the numbers of gradient directions, number of *b*-zero volumes, *b*-values, voxel sizes, etc. Essentially, these subsamples constituted a slightly different protocol within each dataset. Consequently, we used the *dMRI protocol* as the batch effect when training the normative model with HBR. HBR models these site-dependent parameters in a hyperprior, and the model hyperparameters are adapted to incoming data from each site. Subsequently, we ran an HBR model for each DTI metric across all regions that included sex as a second grouping variable in the HBR setting, and age as a fixed independent variable. In this study, we do not expect the biological sources of variance under investigation (e.g., age or sex) to differ systematically across sites. We ran the training and testing of the HBR normative models using the PCNtoolkit package (v29) and Python 3.9 (Details in Supplementary Methods).

### 7. Model Diagnostics

We evaluated the goodness of fit of the HBR models, using the standardized mean squared error (SMSE) and the mean standardized log loss (MSLL) for all four DTI metrics. The SMSE assesses the goodness of fit and central tendency, for which a value lower than 1 indicates a good predictive performance. We also used the mean standardized log loss (MSLL)^20^ which evaluates both the mean and standard deviation of the predicted distribution; a more negative value indicates a better performance. Finally, we included Pearson’s correlation coefficient (Rho) between the observed and predicted WM measures. We also evaluated the calibration of the centiles; that is, if the modeled distribution matches the observed distribution, the Z-scores assigned to a new sample of controls drawn from the same distribution should be normally distributed. First, we tested the normality of the estimated Z-scores of the test data using the Shapiro-Wilk test^75^. We used the test statistic (W) to quantify the degree to which the observed distribution deviates from the assumed normal distribution. Values less than one (1) indicate deviation from normality. Secondly, we calculated the skewness and the kurtosis of the estimated Z-scores to further investigate their distributions. We ran all validation metrics (i.e., SMSE, MSLL, Rho, W, skewness and kurtosis) for each ROI across all 10 experimental iterations. The results are shown in **Figure 4** and **Figures S5-8**. In addition to the model fit evaluations shown in Figure 4 for each WM region, we ran evaluation metrics for each site separately for each DTI metric (see supplementary **Figures S9-12**).

In order to have a basis for comparison to other harmonization methods, we ran ComBat-GAM^83^ to harmonize the raw data across sites. We then ran HBR with the following two models:

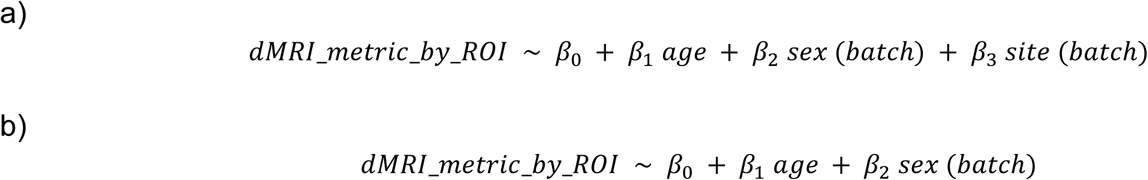

Model a is the same model that we originally ran but with previously harmonized data. Model b assumes that site differences are already removed with harmonization and hence no hierarchical batch effect should be required for the variable site. We tested the site-effects on the Z-scores derived from the three models, the original model from the manuscript and models a and b explained above.

To test whether the validation scores were significantly different or not between our HBR model (without prior ComBat) and the models with prior ComBat harmonization, we ran a Weighted Least Squares regression (WLS) with ROI (n=22) and the number of repetitions (n=10) as fixed effects, weighted by the number of subjects per site and clustering the standard errors by site. This model gives a weighted global mean difference (the intercept) of the validation scores between models by adjusting for ROI and repetition average shifts via fixed effects.

Specifically, for each DTI metric (FA, MD, RD, AD) we ran the following WLS models:

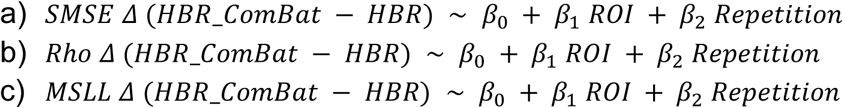

For inference on the intercept (β0) we used cluster-robust (‘sandwich’) standard errors clustered at the site level which allows for arbitrary correlation among ROIs and repetitions within each site, ensuring a non-biased p-value.

In order to test for site effects at the ROI level, we performed the following mixed model linear regression for each ROI, separately:

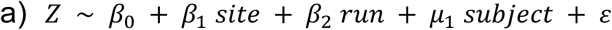

– where site (37 sites) and run (10 runs) are modeled as fixed effects and subject is modeled as a random effect. Each subject has multiple ROIs, so the subject random intercept absorbs the within-subject correlation. The runs contain different subsets of subjects because they are derived from random train/test splits, so run should be a fixed effect. We used a joint Wald test to evaluate whether to reject the hypothesis that all site coefficients were equal to zero. We corrected the resulting p-values across ROIs using FDR.

By using a leave-one-out strategy, we aimed to predict with support vector machines (SVMs) the site at test time for the Global-WM. We specified a linear kernel, we enabled probability estimates and we adjusted the weights inversely proportional to the class frequencies (i.e. the number of subjects per site) in the input data, which gives less weight to larger sites and more weight to smaller sites.

### 8. Plotting of the age trajectories

It is customary for the distribution estimated in a Bayesian model to be computed by sampling from the posterior predictive distribution. To generate samples from the posterior predictive distribution, we use the posterior samples {θ_*i*_} and generate new data points 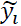 from the likelihood function *p*( *y* ∣ θ_*i*_). To this end, for each posterior sample θ_*i*_, a corresponding sample 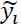 from the predictive distribution conditioned on θ_*i*_ is drawn. We generated these samples for the 10 repetitions of the HBR experiment (each with a different 80%-20% train-test split). To model the trajectory of each of the DTI metrics for each WM

ROI, with respect to age and for males and females separately, we used the generative features of the hierarchical Bayesian model. For each DTI metric per ROI we generated samples for the age range 4 to 91 years and for both sexes 20 times and for one site only. We picked CAMCAN as the site for the simulations of the Global-WM (**Figures 1D and 2**). After generating these samples, we averaged the predicted data across the 20 iterations to generate a smooth curve. In **Figure 1B** we generated samples for 19 sites for the Global-WM to display the site differences.

### 9. Anomaly detection on the reference cohort

Training and testing data sets were created with an 80% to 20% sample split, stratified by site, based on the total number of 54,583 participants. To achieve stability, we repeated the same procedure 10 times by using Scikit-learn’s ‘train_test_split’ function with different random seeds, thus ensuring a different split each time. To ensure robustness against data perturbations, we ran HBR 10 times on each of the 80% sample splits (n=43,666). Z-scores were re-calculated on each of the 10 test sets (20% sample split; n=10,917). Z-scores were also calculated for the Alzheimer’s dementia and MCI cases from ADNI3 and OASIS3. We calculated the probabilities of abnormality *P*(*Z*), i.e., the abnormality scores, from the Z-scores for both controls (20% sample split) and clinical samples (dementia and MCI) using the following formula^17^:

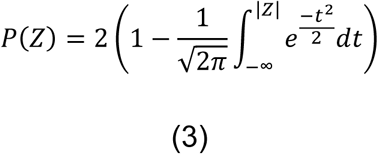

ROI-wise areas under the ROC curves (AUCs) were calculated to determine the classification accuracy of the computed deviations. Given the normal distribution of z-scores and without any assumption on the direction of abnormal samples (left or right tail), the extreme deviations from the norm can be quantified in the form of a probability by computing the area of the shaded region of the cumulative distribution function at |z|. If there are no deviations, then the probability of abnormality is 0 and it is closer to 1 as |z| increases. This is a monotone transform because, as |z| increases, the tail probability of being an abnormal sample increases monotonically (strictly increasing), which does not affect the ROC analysis. Subsequently, we performed permutation tests with 1,000 random samples to derive permutation *p*-values for each DTI metric per ROI, and applied a false discovery rate (FDR) correction on each DTI metric separately across regions to identify those that showed significant group differences. The regions that showed significance in 9 out of the 10 experimental iterations were retained^4,17^. For each clinical group, i.e., dementia and MCI, we summarized the individual deviations within each group by first separating them into positive and negative deviations, counting how many subjects had an extreme deviation (defined as Z>2 for positive deviations, Z<-2 for negative deviations) at a given ROI, and then dividing by the group size (see **Figure 6**).

### 10. Model adaptation

Following the development phase, we aimed to deploy the trained normative models by transferring the estimated hyperparameters to new datasets. This deployment strategy, called model adaptation, allows us to re-infer the parameters of the model at local sites. We used the HBR model adaptation function called ‘transfer’ available in the PCNtoolkit v29^17^. Here, we took 50% of the healthy control subjects from the new sites to adapt the parameters of the reference model to the local model. Then the Z-scores for the remaining 50% healthy controls and the cases of the new incorporated sites are computed as described above in Methods section 5. We tested the model adaptation scenario on two cohorts with pathologies affecting the brain at two different stages of the lifespan, i.e., UCLA and NIMHANS (see **Table 1**). The UCLA dataset includes people with a rare neurogenetic condition who carry chromosome 22q11.2 CNVs (copy number variants; 22qDel) that affect brain development and have a higher risk for future psychiatric conditions such as neurodevelopmental delay, autism spectrum disorder, attention deficit hyperactivity disorder and schizophrenia. The NIMHANS dataset has patients with MCI and Alzheimer’s dementia, both affecting the brain in later stages of life. Once the normative models were adapted to these new sites, we summarized the individual deviations within each group (22qDel, MCI and dementia) by first separating them into positive and negative deviations, counting how many subjects had extreme deviations at a given ROI, and then dividing by the group size (see **Figure 7**).

### 11. Retrogenesis of WM Regions

We tested two retrogenesis hypotheses that have been proposed for drawing general principles from the lifespan charts of brain metrics. Here we use the term peak for all DTI metrics, that is, for the maximum of FA and for the minimum of MD, AD, RD. To test the “gain-predicts-loss” hypothesis, we ran Spearman correlations between the percent change from age 4 years to the peak and the percent change between the peak and age 91 years. By doing so, we are testing correlations between metrics derived from the two sides of the curve, the maturation and the decline during aging. According to the “gain-predicts-loss” hypothesis both sides of the curve should have slopes that are correlated, i.e., the rate of senescence should depend on the rate of maturation. We assessed statistical significance with spin permutation tests with 1,000 rotations using the ENIGMA-toolbox^84^ to take into account the spatial correlation structure, slightly modified to fit the permutations across the WM regions in the JHU atlas. The other hypothesis, the “last-in first-out” hypothesis, assumes that the cortical regions of higher complexity (supporting higher order functions) have a slower maturation rate and may be more sensitive to degeneration after reaching their maturational peak. By extension, the WM tracts connecting these more complex cortical regions should also have a slower maturation rate. We first calculated the percent change for each DTI metric every 10 years across the lifespan for each ROI (4-14,15-24, 25-34, 35-44, 45-54, 55-64, 65-74, 75-84, 85-91 years; **Figure 5**). We then assessed associations between the percent change from age 4-14 years, 75-84 years and 85-91 years with the age of peak, by also using spin permutation tests. If the “last in, first-out” were true, a higher percent change during development should be correlated with a younger age of peak, and a higher percent change in senescence should be correlated with an older age of peak.

## Data Availability

The estimated normative models’ data generated in this study have been deposited in the PCNPortal^5^ database under accession code [https://pcnportal.dccn.nl/]. Source data are provided with this paper.

The data generated in this study, such as source data files containing summary statistics underlying each figure (e.g., group means, standard errors, regression coefficients, model outputs, and aggregated metrics) are provided in the Supplementary Information and Source Data files.

We fully support the journal’s commitment to transparency and reproducibility. However, several datasets used in this study (ABCD, UK Biobank, ADNI) are governed by Data Use Agreements that prohibit redistribution of individual participant-level data, including publication as supplementary .csv files and/or sharing on online repositories (e.g. Github).

Researchers wishing to reproduce analyses using the underlying individual-level data may obtain access directly through the respective data access procedures:

- ABCD Study: https://nda.nih.gov/abcd
- UK Biobank: https://www.ukbiobank.ac.uk/enable-your-research
- ADNI: https://adni.loni.usc.edu/data-samples/access-data/

## Code Availability

Our analysis code and full description of the normative modeling pipeline are publicly available to ensure reproducibility. The code will be available in this repository: https://github.com/villalonreina/ENIGMA-DTI-Normative-Modeling, which is also forked here: https://github.com/ENIGMA-git/ENIGMA-DTI-Normative-Modeling

## The Alzheimer’s Disease Neuroimaging Initiative (ADNI)

Data used in preparation of this article were obtained from the Alzheimer’s Disease Neuroimaging Initiative (ADNI) database (adni.loni.usc.edu). As such, the investigators within the ADNI contributed to the design and implementation of ADNI and/or provided data but most of them did not participate in analysis or writing of this report. The consortium members who qualified for authorship are: Paul M. Thompson, Neda Jahanshad, Talia M. Nir and Sophia Thomopoulus. A complete listing of ADNI investigators may be found at: http://adni.loni.usc.edu/wp-content/uploads/how_to_apply/ADNI_Acknowledgement_List.pdf”

## Acknowledgments

We thank Emily Laltoo for her contributions to data processing for the study.

**ABCD**. Some of the data used in preparing this article were obtained from the Adolescent Brain Cognitive Development SM (ABCD) Study (https://abcdstudy.org), held in the NIMH Data Archive (NDA). This is a multisite, longitudinal study designed to recruit more than 10,000 children aged 9-10 and follow them over 10 years into early adulthood. The ABCD Study® is supported by the National Institutes of Health and additional federal partners under award numbers U01DA041048, U01DA050989, U01DA051016, U01DA041022, U01DA051018, U01DA051037, U01DA050987, U01DA041174, U01DA041106, U01DA041117, U01DA041028, U01DA041134, U01DA050988, U01DA051039, U01DA041156, U01DA041025, U01DA041120, U01DA051038, U01DA041148, U01DA041093, U01DA041089, U24DA041123, U24DA041147. A full list of supporters is available at https://abcdstudy.org/federal-partners.html. A listing of participating sites and a complete listing of the study investigators can be found at https://abcdstudy.org/consortium_members/. ABCD consortium investigators designed and implemented the study and/or provided data but did not necessarily participate in the analysis or writing of this report. This manuscript reflects the views of the authors and may not reflect the opinions or views of the NIH or ABCD consortium investigators. The ABCD data repository grows and changes over time. The ABCD data used in this report came from 10.15154/1523041. DOIs can be found at https://dx.doi.org/10.15154/1523041.

**ADNI**. Data used in the preparation of this article were obtained from the Alzheimer’s Disease Neuroimaging Initiative (ADNI) database (adni.loni.usc.edu). The ADNI was launched in 2003 as a public-private partnership, led by Principal Investigator Michael W. Weiner, MD. The primary goal of ADNI has been to test whether serial magnetic resonance imaging (MRI), positron emission tomography (PET), other biological markers, and clinical and neuropsychological assessment can be combined to measure the progression of mild cognitive impairment (MCI) and early Alzheimer’s disease (AD). For up-to-date information, see www.adni-info.org. Data collection and sharing for the ADNI project was funded by the Alzheimer’s Disease Neuroimaging Initiative (ADNI) (National Institutes of Health Grant U01 AG024904) and DOD ADNI (Department of Defense award number W81XWH-12-2-0012). ADNI is funded by the National Institute on Aging, the National Institute of Biomedical Imaging and Bioengineering, and through generous contributions from the following: AbbVie, Alzheimer’s Association; Alzheimer’s Drug Discovery Foundation; Araclon Biotech; BioClinica, Inc.; Biogen; Bristol-Myers Squibb Company; CereSpir, Inc.; Cogstate; Eisai Inc.; Elan Pharmaceuticals, Inc.; Eli Lilly and Company; EuroImmun; F. Hoffmann-La Roche Ltd and its affiliated company Genentech, Inc.; Fujirebio; GE Healthcare; IXICO Ltd.; Janssen Alzheimer Immunotherapy Research & Development, LLC.; Johnson & Johnson Pharmaceutical Research & Development LLC.; Lumosity; Lundbeck; Merck & Co., Inc.;Meso Scale Diagnostics, LLC.; NeuroRx Research; Neurotrack Technologies; Novartis Pharmaceuticals Corporation; Pfizer Inc.; Piramal Imaging; Servier; Takeda Pharmaceutical Company; and Transition Therapeutics. The Canadian Institutes of Health Research is providing funds to support ADNI clinical sites in Canada. Private sector contributions are facilitated by the Foundation for the National Institutes of Health (www.fnih.org). The grantee organization is the Northern California Institute for Research and Education, and the study is coordinated by the Alzheimer’s Therapeutic Research Institute at the University of Southern California. ADNI data are disseminated by the Laboratory for Neuro Imaging at the University of Southern California.

**AOMIC**. The Amsterdam Open MRI Collection (AOMIC) is a collection of three datasets with multimodal (3T) MRI data. All raw/preprocessed data is publicly available from the Openneuro data sharing platform: ID1000: https://openneuro.org/datasets/ds003097 PIOP1: https://openneuro.org/datasets/ds002785 PIOP2: https://openneuro.org/datasets/ds002790

**CAMCAN**. The Cambridge Centre for Ageing and Neuroscience (CAMCAN) was supported by the UK Biotechnology and Biological Sciences Research Council (grant number BB/H008217/1), together with support from the UK Medical Research Council Cognition & Brain Sciences Unit (CBU) and University of Cambridge, UK. We are grateful to the CAMCAN respondents and their primary care teams in Cambridge for their participation in the CAMCAN study. We also thank colleagues at the MRC Cognition and Brain Sciences Unit MEG and MRI facilities for their assistance.

**CHCP**. Data were provided [in part] by the Chinese Human Connectome Project (CHCP, PI: Jia-Hong Gao) funded by the Beijing Municipal Science & Technology Commission, Chinese Institute for Brain Research (Beijing), National Natural Science Foundation of China, and the Ministry of Science and Technology of China.

**CHBMP**. The Cuban Human Brain Mapping Project (CHBMP) repository is an open multimodal neuroimaging and cognitive dataset available for registered users on the LORIS repository which is part of the MNI neuroinformatics ecosystem at: https://chbmp-open.loris.ca/

**HBN**. Healthy Brain Network Biobank **(HBN)** data. The dataset is described in Nature Scientific Data: Alexander, L. et al. An open resource for transdiagnostic research in pediatric mental health and learning disorders. Scientific Data 4, 170181 (2017). https://doi.org/10.1038/sdata.2017.181

**HCP-Aging.** Research reported in this publication was supported by the National Institute of Mental Health under grant U01MH109589 and by the McDonnell Center for Systems Neuroscience, at Washington University in St. Louis. The **HCP-Development** 2.0 Release data analyzed in this report came from DOI: 10.15154/1520708. The HCP-Aging 2.0 Release data used in this report came from DOI: 10.15154/1520707.

**HCP-YA** data were provided [in part] by the Human Connectome Project, WU-Minn Consortium (Principal Investigators: David Van Essen and Kamil Ugurbil; U54 MH091657) funded by the 16 NIH Institutes and Centers that support the NIH Blueprint for Neuroscience Research; and by the McDonnell Center for Systems Neuroscience at Washington University.

**PedsDTI**. Data used in the preparation of this article were obtained from the **Pediatric MRI Data Repository created by the NIH MRI Study of normal brain development (PedsDTI)**. This is a multi-site, longitudinal study of typically developing children, from ages newborn through young adulthood, conducted by the Brain Development Cooperative Group and supported by the National Institute of Child Health and Human Development, the National Institute on Drug Abuse, the National Institute of Mental Health, and the National Institute of Neurological Disorders and Stroke (Contract #s N01-HD02-3343, N01-MH9-0002, and N01-NS-9-2314, N01-NS-9-2315, N01-NS-9-2316, N01-NS-9-2317, N01-NS-9-2319 and N01-NS-9-2320). Disclaimer: The views herein do not necessarily represent the official views of the National Institute of Child Health and Human Development, the National Institute on Drug Abuse, the National Institute of Mental Health, the National Institute of Neurological Disorders and Stroke, the NIH, the US Department of Health and Human Services, or any other agency of the US Government.

**OASIS-3.** Data were provided [in part] by OASIS-3: Longitudinal Multimodal Neuroimaging: Principal Investigators: T. Benzinger, D. Marcus, J. Morris; NIH P30 AG066444, P50 AG00561, P30 NS09857781, P01 AG026276, P01 AG003991, R01 AG043434, UL1 TR000448, R01 EB009352. AV-45 doses were provided by Avid Radiopharmaceuticals, a wholly owned subsidiary of Eli Lilly.

**PING**. Data collection and subsequent dataset for this project were obtained from the Pediatric Imaging, Neurocognition and Genetics Study (PING), National Institutes of Health Grant RC2DA029475. PING is funded by the National Institute on Drug Abuse and the Eunice Kennedy Shriver National Institute of Child Health & Human Development. PING data are disseminated by the PING Coordinating Center at the Center for Human Development, University of California, San Diego, as detailed in Jernigan et al (2016).

**PNC**. The Philadelphia Neurodevelopmental Cohort (PNC) data are available at: https://www.med.upenn.edu/bbl/philadelphianeurodevelopmentalcohort.html

**PPMI**. Data used in the preparation of this article were obtained on August 25, 2022, from the Parkinson’s Progression Markers Initiative (PPM**I**) database (www.ppmi-info.org/access-data-specimens/download-data), RRID:SCR 006431. For up-to-date information on the study, visit www.ppmi-info.org. PPMI – a public-private partnership – is funded by the Michael J. Fox Foundation for Parkinson’s Research and funding partners, including 4D Pharma, Abbvie, AcureX, Allergan, Amathus Therapeutics, Aligning Science Across Parkinson’s, AskBio, Avid Radiopharmaceuticals, BIAL, BioArctic, Biogen, Biohaven, BioLegend, BlueRock Therapeutics, Bristol-Myers Squibb, Calico Labs, Capsida Biotherapeutics, Celgene, Cerevel Therapeutics, Coave Therapeutics, DaCapo Brainscience, Denali, Edmond J. Safra Foundation, Eli Lilly, Gain Therapeutics, GE HealthCare, Genentech, GSK, Golub Capital, Handl Therapeutics, Insitro, Janssen Neuroscience, Jazz Pharmaceuticals, Lundbeck, Merck, Meso Scale Discovery, Mission Therapeutics, Neurocrine Biosciences, Neuropore, Pfizer, Piramal, Prevail Therapeutics, Roche, Sanofi, Servier, Sun Pharma Advanced Research Company, Takeda, Teva, UCB, Vanqua Bio, Verily, Voyager Therapeutics, the Weston Family Foundation and Yumanity Therapeutics.

**QTAB**. The Queensland Twin Adolescent Brain Project (QTAB) project resource was produced as a result of i) the goodwill and contribution of 422 twin/triplet participants and their parents, ii) funding from the National Health and Medical Research Council, Australia (APP1078756) and the Queensland Brain Institute, University of Queensland, iii) access to several key resources, including the Centre for Advanced Imaging, the Human Studies Unit, Institute of Molecular Bioscience, and the Queensland Cyber Infrastructure Foundation, at the University of Queensland, local and national twin registries at the QIMR Berghofer Medical Research Institute and Twin Research Australia, as well as the many assessments made available by researchers worldwide, and iv) was established with the purpose of promoting the conduct of health-related research in adolescence.

**QTIM**. The Queensland Twin IMaging (QTIM) study is forever grateful to the twins and siblings for their willingness to participate in our studies. We thank Marlene Grace and Ann Eldridge for participant recruitment; Kerrie McAloney for study co-ordination; Kori Johnson, Aaron Quiggle, Natalie Garden, Matthew Meredith, Peter Hobden, Kate Borg, Aiman Al Najjar and Anita Burns for data acquisition; David Butler and Daniel Park for IT support. The QTIM study was supported by the National Institute of Child Health and Human Development (R01 HD050735) and the National Health and Medical Research Council (496682, 1009064).

**SLIM.** The Southwest University Longitudinal Imaging Multimodal (SLIM) Brain Data Repository was supported by the National Natural Science Foundation of China (31271087; 31470981; 31571137; 31500885), National Outstanding young people plan the Program for the Top Young Talents by Chongqing, the Fundamental Research Funds for the Central Universities (SWU1509383,SWU1509451), Natural Science Foundation of Chongqing (cstc2015jcyjA10106), Fok Ying Tung Education Foundation (151023), General Financial Grant from the China Postdoctoral Science Foundation (2015M572423, 2015M580767), Special Funds from the Chongqing Postdoctoral Science Foundation (Xm2015037), Key research for Humanities and social sciences of Ministry of Education(14JJD880009).

**UKBB**. This research was conducted using data from UK Biobank, a major biomedical database, under application number 11559. UK Biobank is generously supported by its founding funders the Wellcome Trust and UK Medical Research Council, as well as the Department of Health, Scottish Government, the Northwest Regional Development Agency, British Heart Foundation and Cancer Research UK.

## Author contributions Statement

J.E.V.R. contributed with Conceptualization, Data Curation, Formal Analysis, Investigation, Writing

– Original Draft Preparation, Investigation, Software, Visualization

A.H.Z. contributed with Writing – Review & Editing, Data Curation, Project Administration

S.B. contributed with Data Curation

C.A.M. contributed with Writing – Review & Editing

Y.F. contributed with Data Curation

T.C. contributed with Data Curation

L.N. contributed with Data Curation

L.K. contributed with Data Curation, Project Administration, Project Administration

C.E.B. contributed with Funding Acquisition, Writing – Review & Editing, Resources

J.P.J. contributed with Funding Acquisition, Resources – Review & Editing

H.J. contributed with Data Curation

S.I.T. contributed with Data Curation, Project Administration

I.B.G. contributed with maintaining the computational framework

K.E.L. contributed with Data Curation, Writing – Review & Editing

T.M.N. contributed with Data Curation, Writing – Review & Editing

N.J. contributed with Funding Acquisition, Writing – Review & Editing, Resources, Supervision

S.M.K. contributed with Methodology, Software, Writing – Review & Editing, Supervision, Software

A.F.M. contributed with Methodology, Writing – Review & Editing

P.M.T. contributed with Conceptualization, Funding Acquisition, Supervision, Writing – Review & Editing, Resources

