## Supplements for "Lifespan Normative Modeling of Brain Microstructure"

### Supplements (Article: Lifespan Normative Modeling of Brain Microstructure)

**Figure S1**.

**Trajectories and centile curves of Fractional Anisotropy (FA).** Normative models from age 4 to 91 years for all 21 white matter regions of interest. The trajectories represent the average of the simulated data across the 10 different train_test data splits.

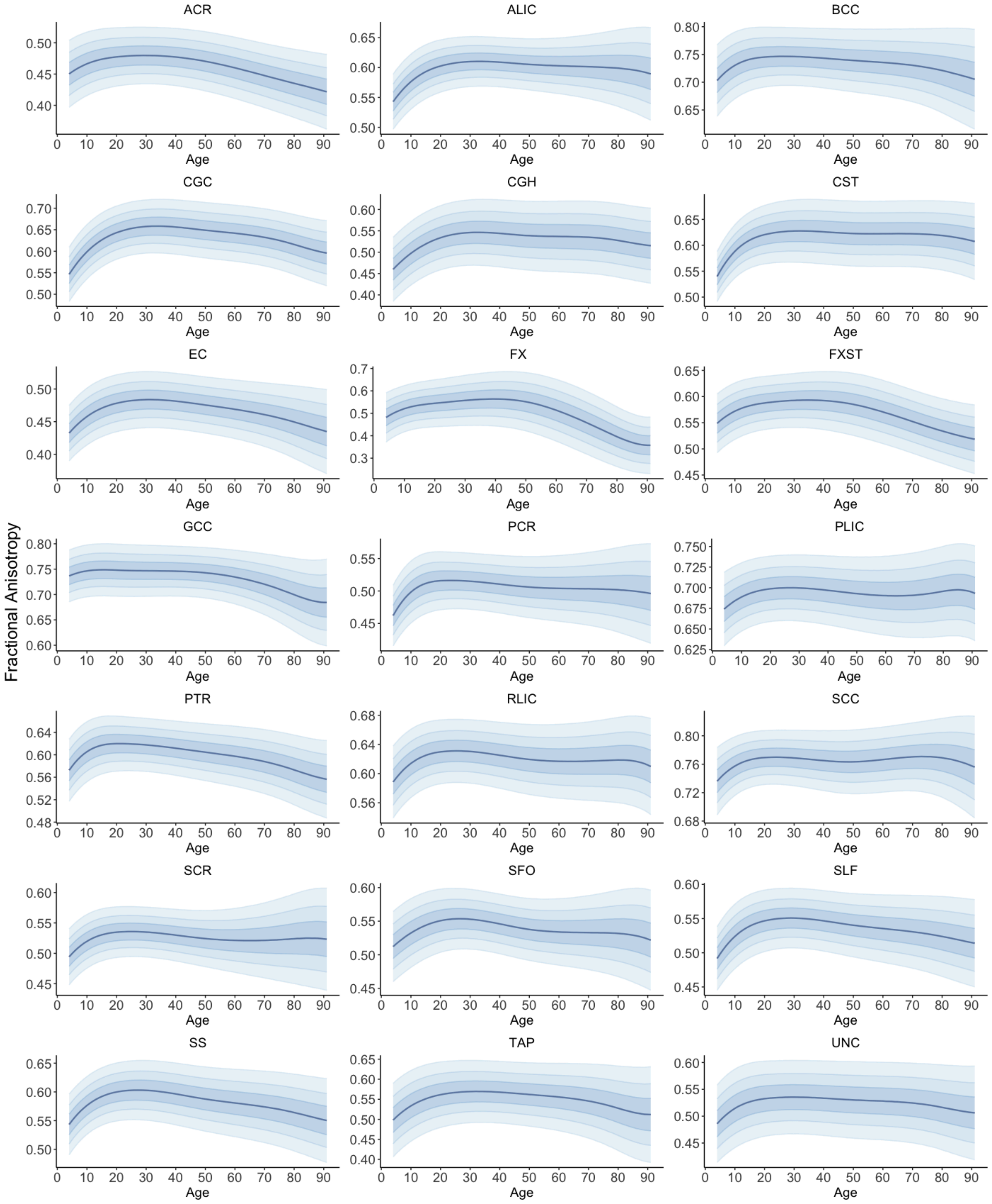

Figure S2.

**Trajectories and centiles of mean diffusivity (MD).** Normative models from age 4 to 91 years for all 21 white matter regions of interest. The trajectories represent the average of the simulated data across the 10 different train_test data splits.

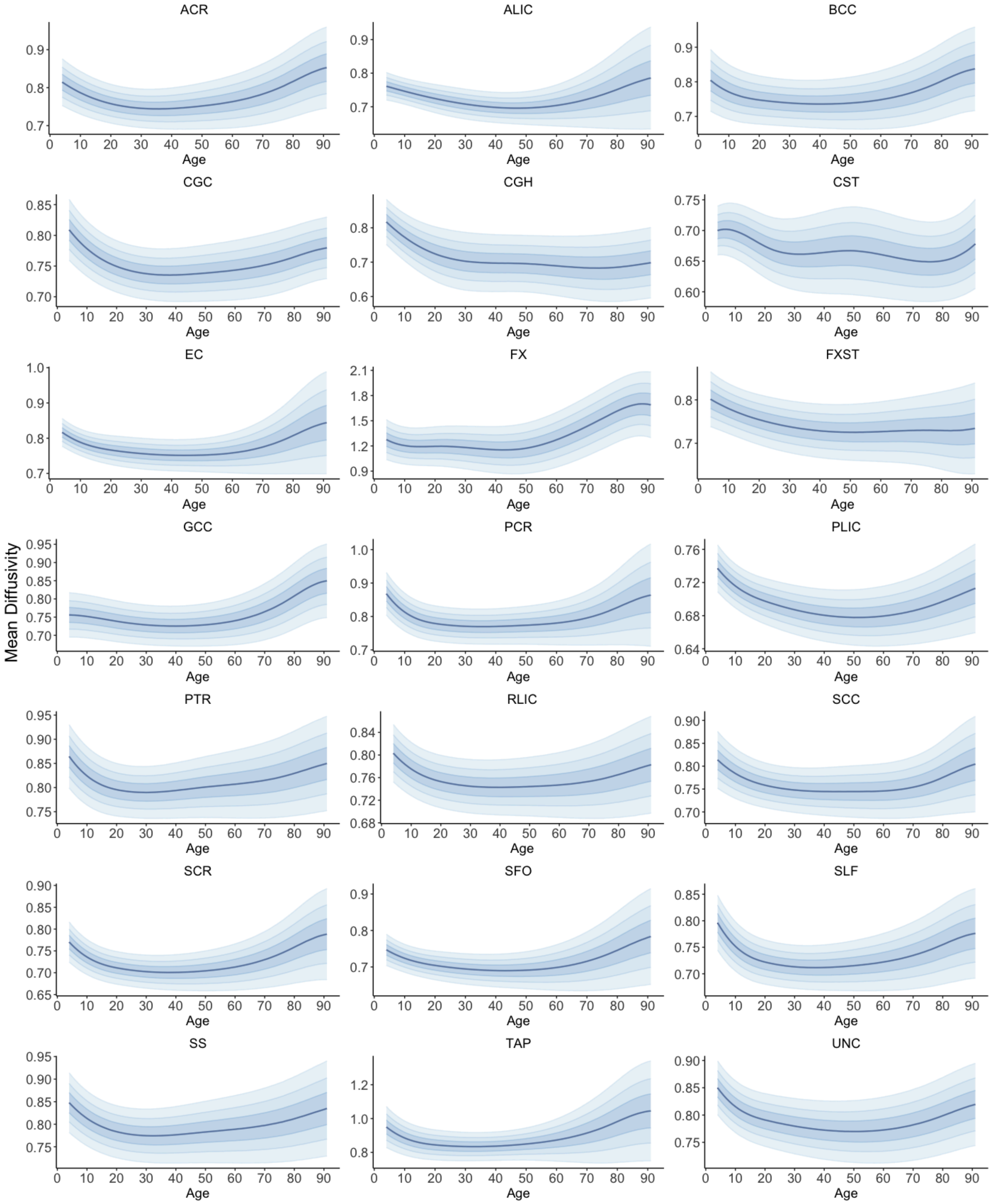

Figure S3.

**Trajectories and centiles of axial diffusivity (AD).** Normative models from age 4 to 91 years for all 21 white matter regions of interest. The trajectories represent the average of the simulated data across the 10 different train_test data splits.

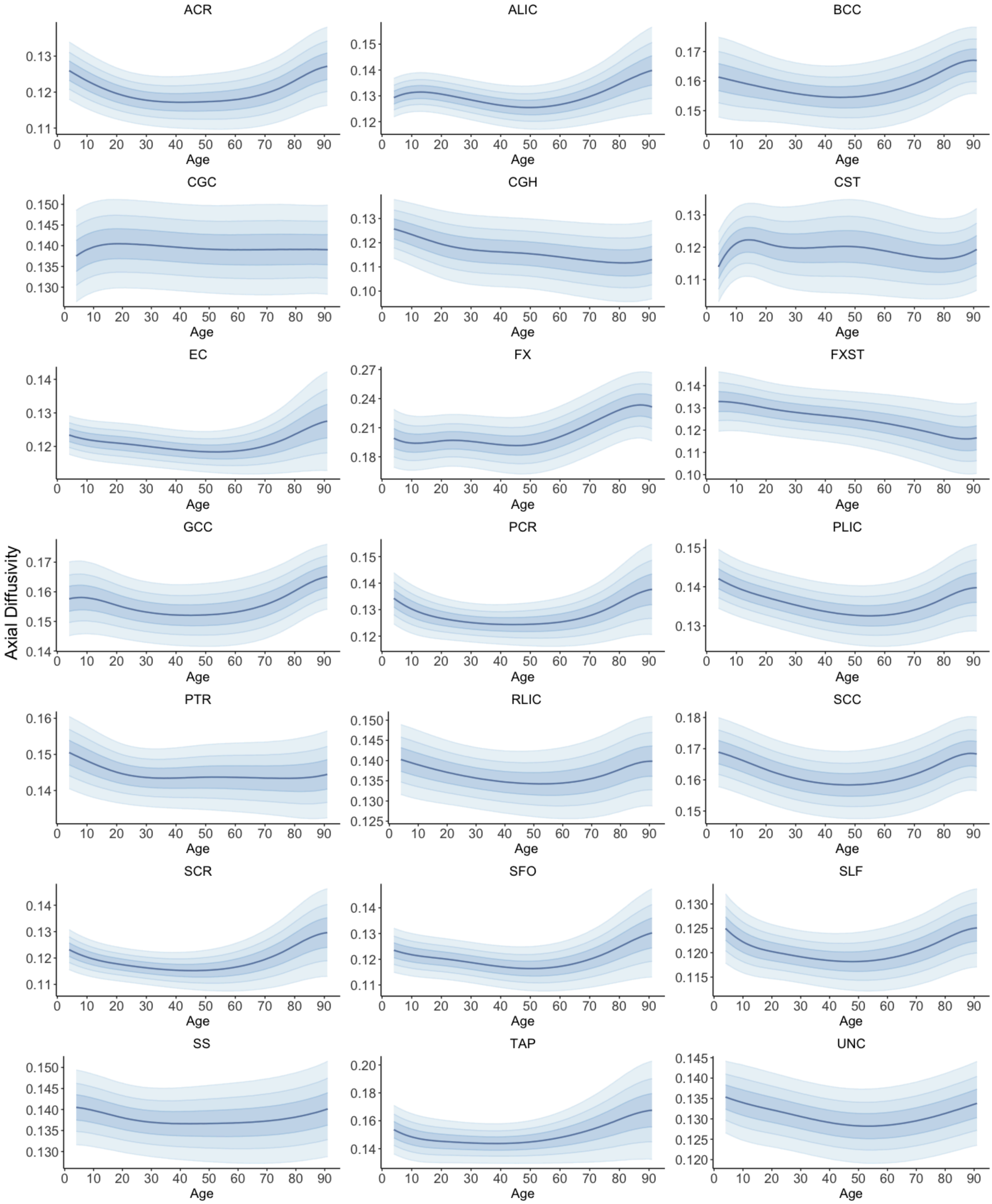

Figure S4.

**Trajectories and centiles of radial diffusivity (RD)**. Normative models from age 4 to 91 years for all 21 white matter regions of interest. The trajectories represent the average of the simulated data across the 10 different train_test data splits.

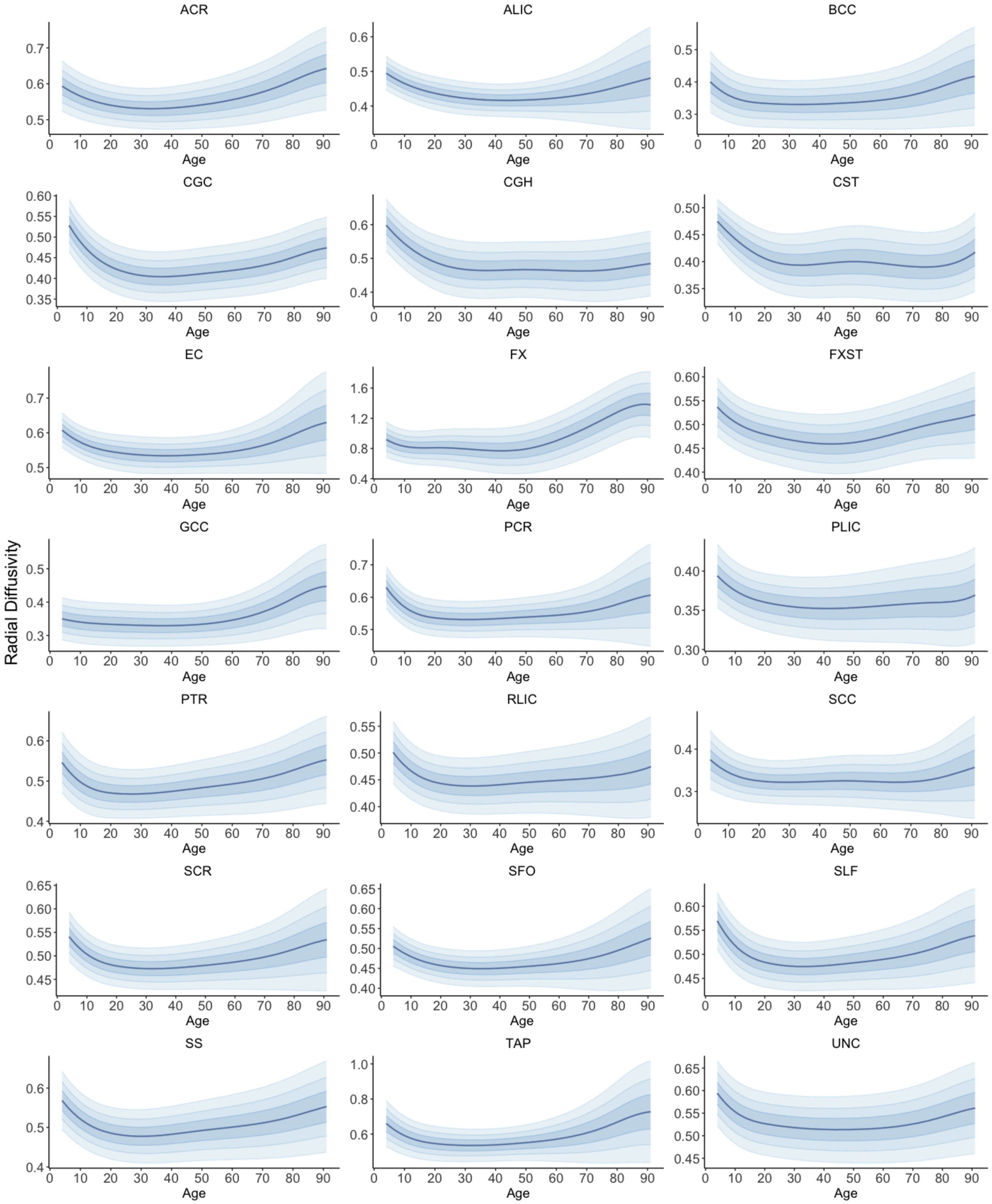

Figure S5.

**Model Diagnostics.** Model diagnostics metrics (Shapiro Wilk's *W*, kurtosis, skewness, SMSE, MSLL, Rho) for Fractional Anisotropy (FA) for each ROI (22 ROIs, 10 experimental repetitions).

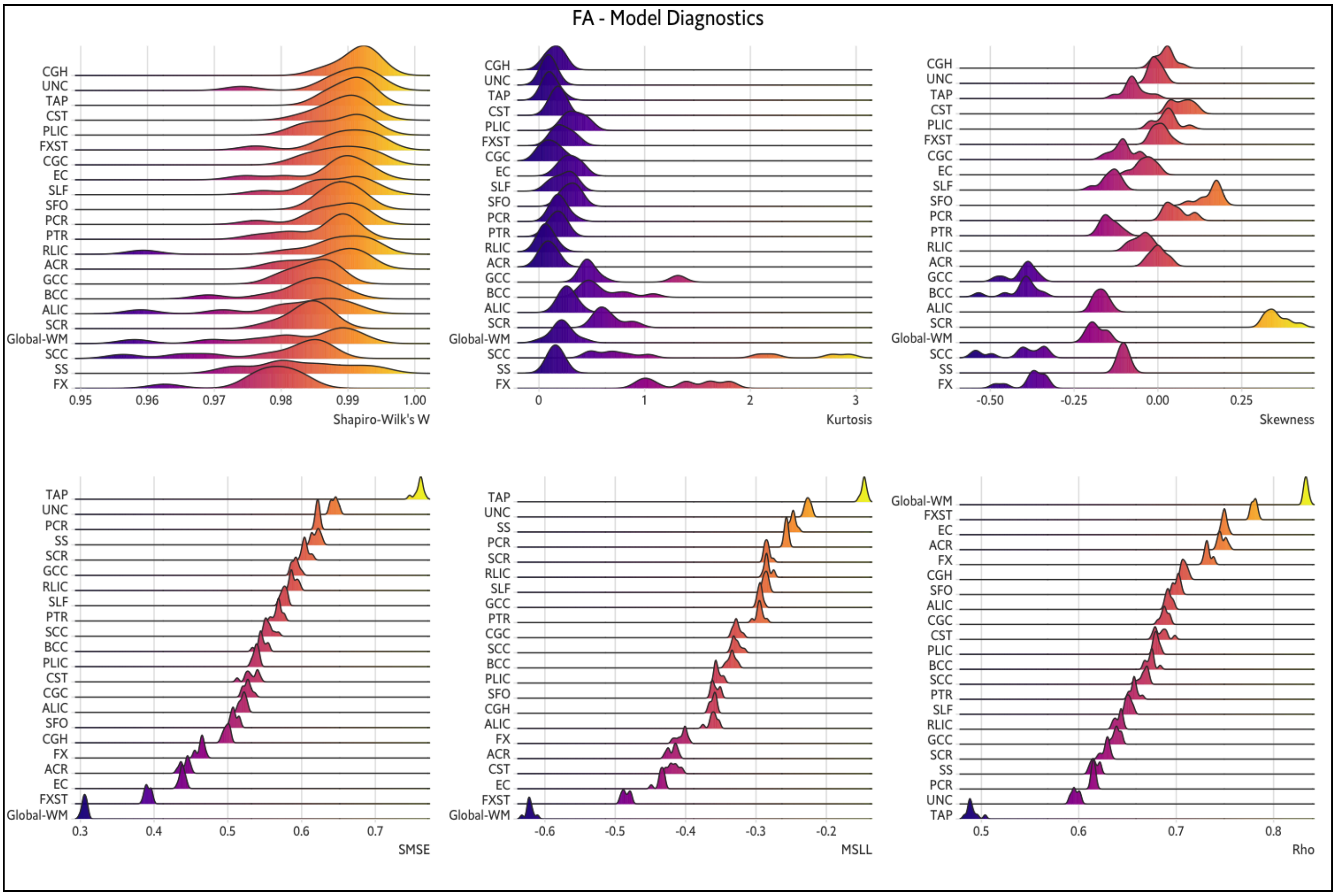

Figure S6.

**Model Diagnostics.** Model diagnostics metrics (Shapiro Wilk's *W*, kurtosis, skewness, SMSE, MSLL, Rho) for Mean Diffusivity (MD) for each ROI (22 ROIs, 10 experimental repetitions).

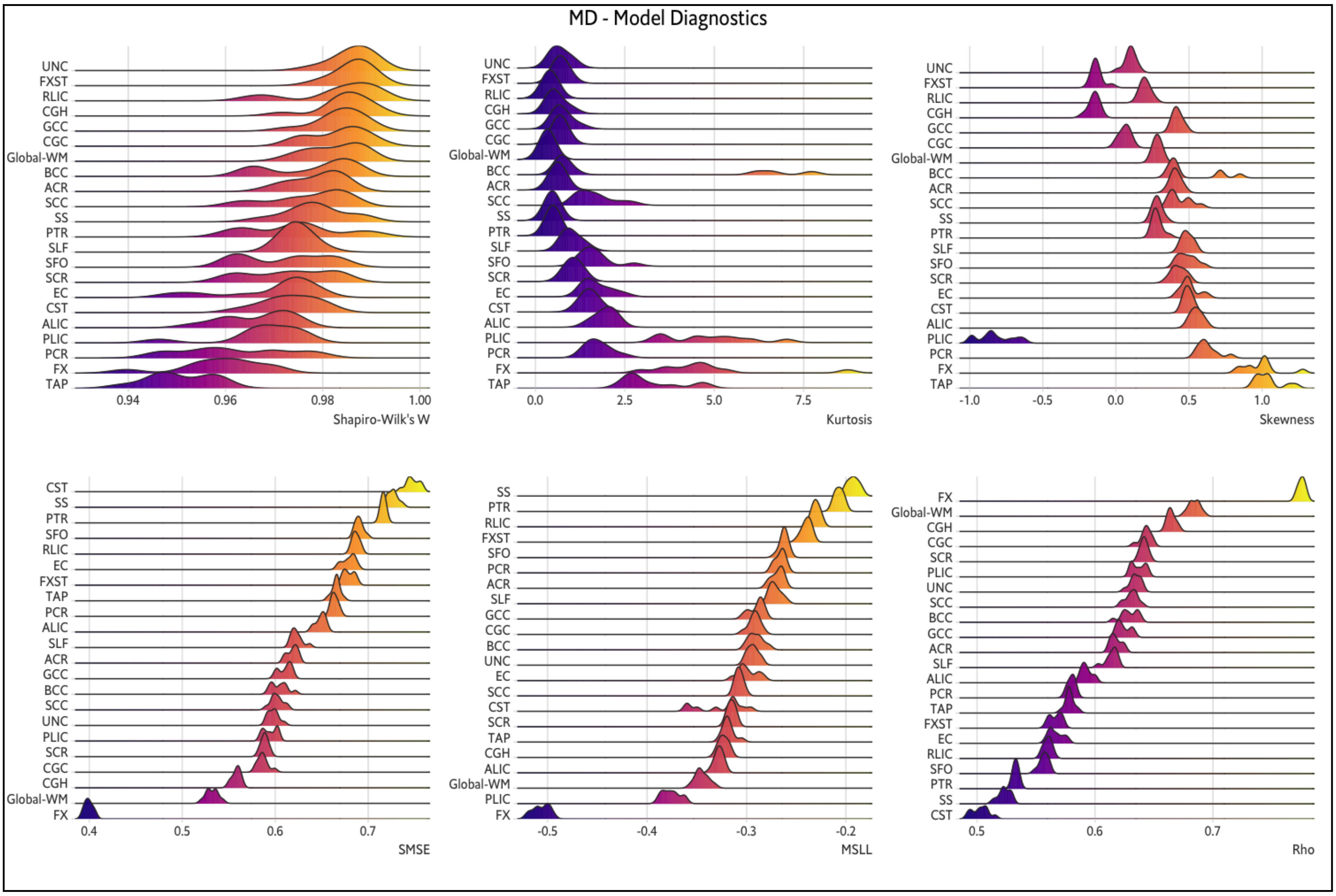

Figure S7.

**Model Diagnostics.** Model diagnostics metrics (Shapiro Wilk's *W*, kurtosis, skewness, SMSE, MSLL, Rho) for Axial Diffusivity (AD) for each ROI (22 ROIs, 10 experimental repetitions).

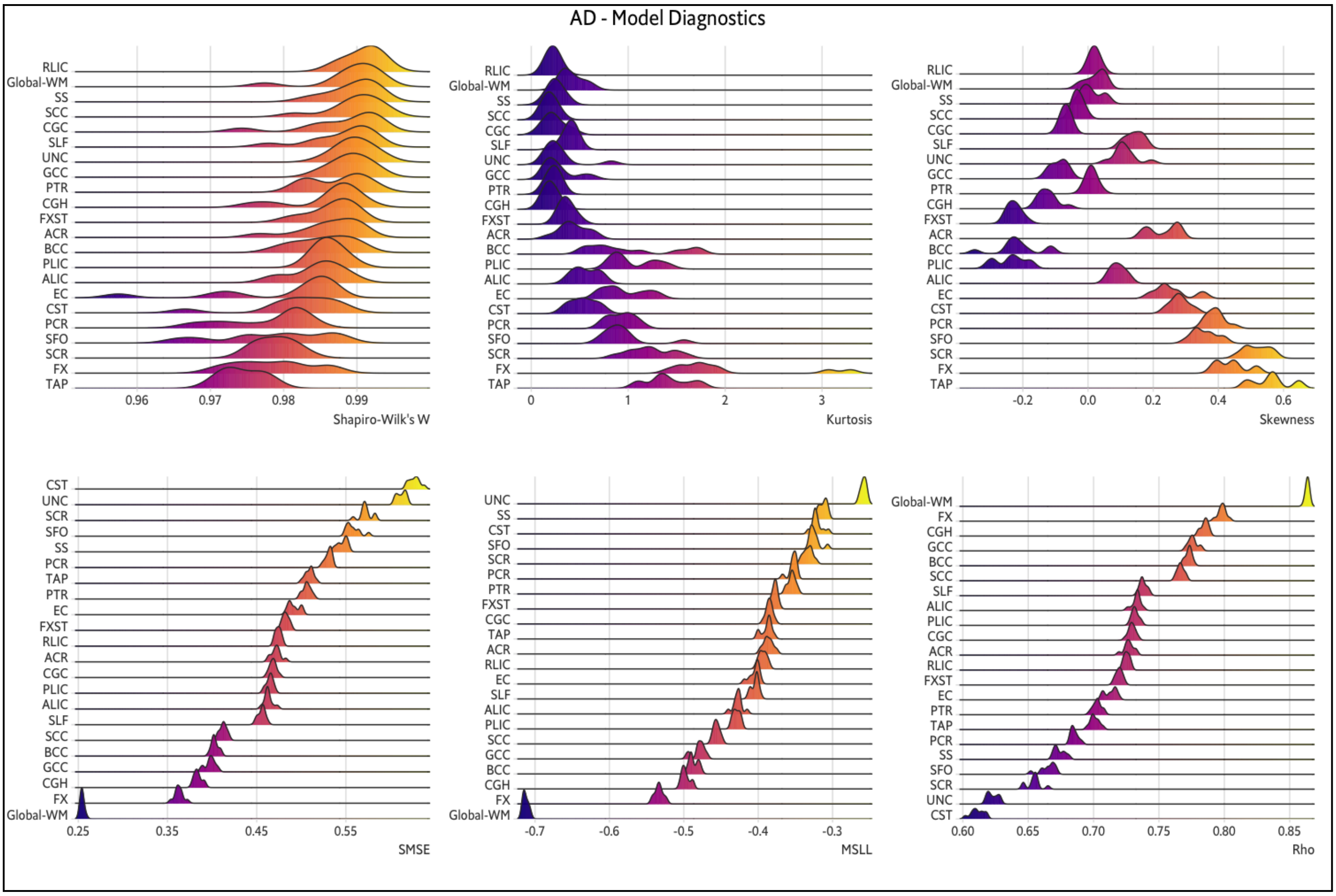

Figure S8.

**Model Diagnostics.** Model diagnostics metrics (Shapiro Wilk's *W*, kurtosis, skewness, SMSE, MSLL, Rho) for Radial Diffusivity (RD) for each ROI (22 ROIs, 10 experimental repetitions).

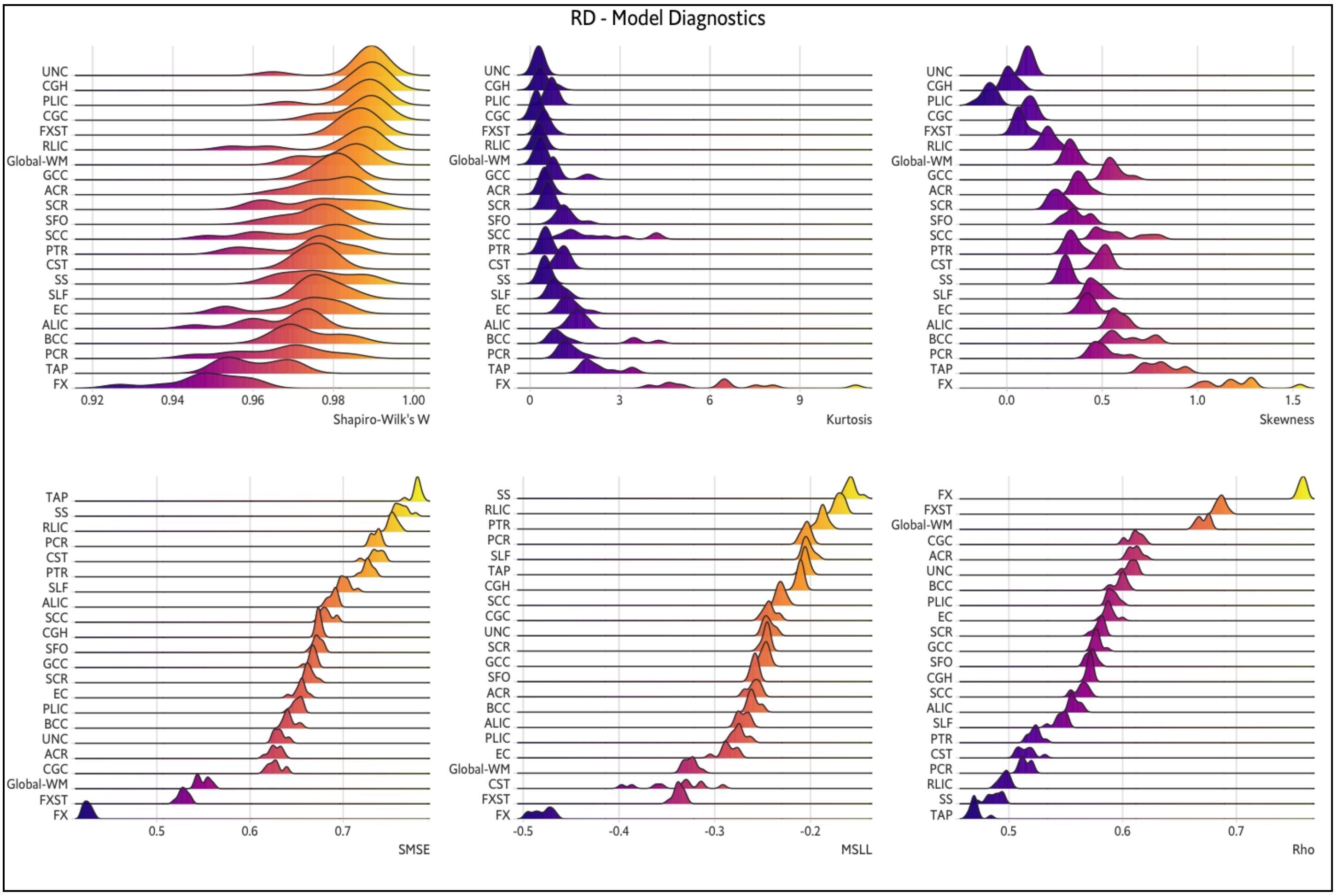

Figure S9.

**Model Diagnostics per site.** Model diagnostics metrics (SMSE, MSLL, Rho) for Fractional Anisotropy (FA) per site (37 sites, 10 experimental iterations).

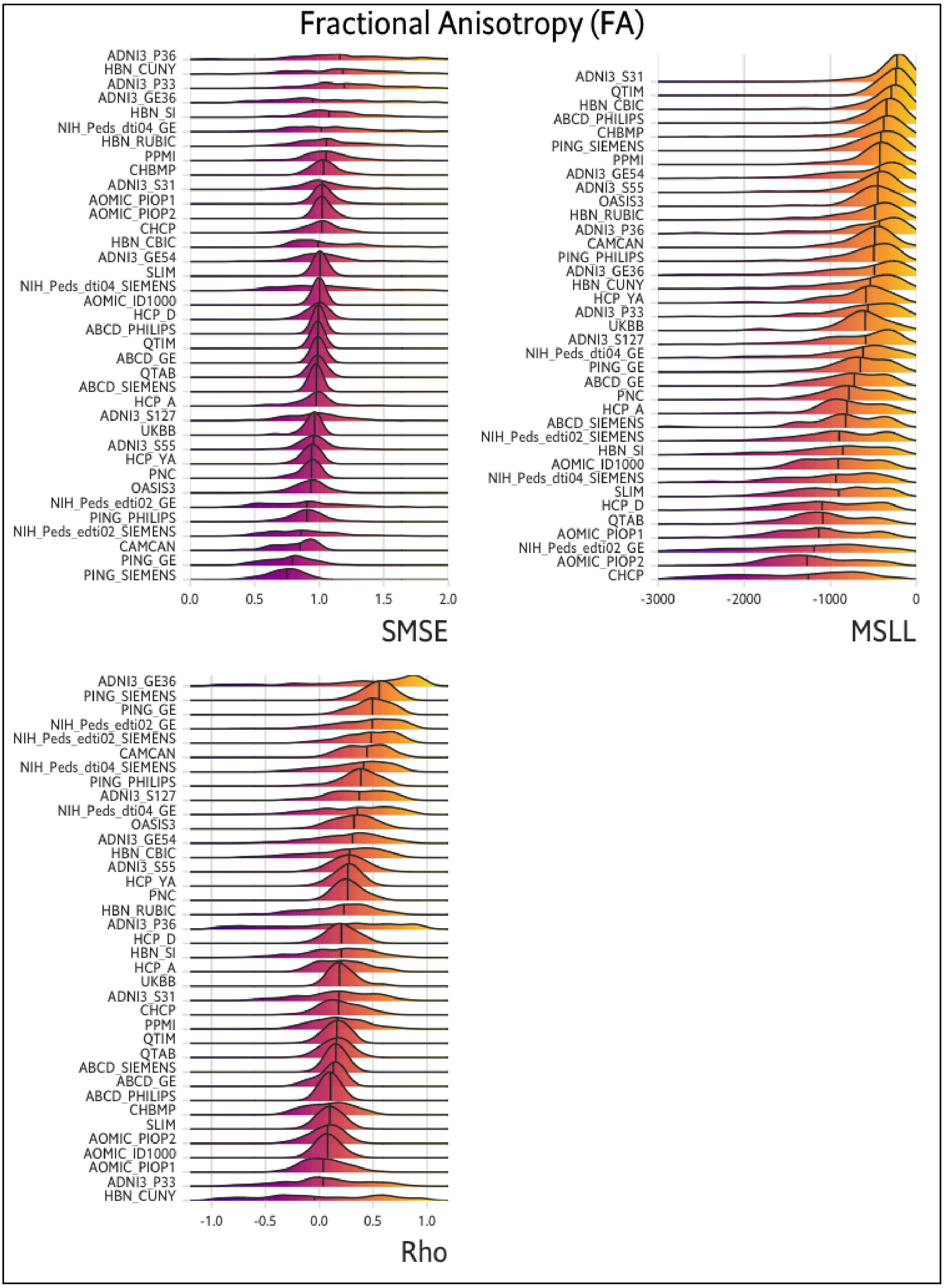

Figure S10

**Model Diagnostics per site.** Model diagnostics metrics (SMSE, MSLL, Rho) for Mean Diffusivity (MD) per site. (37 sites, 10 experimental iterations).

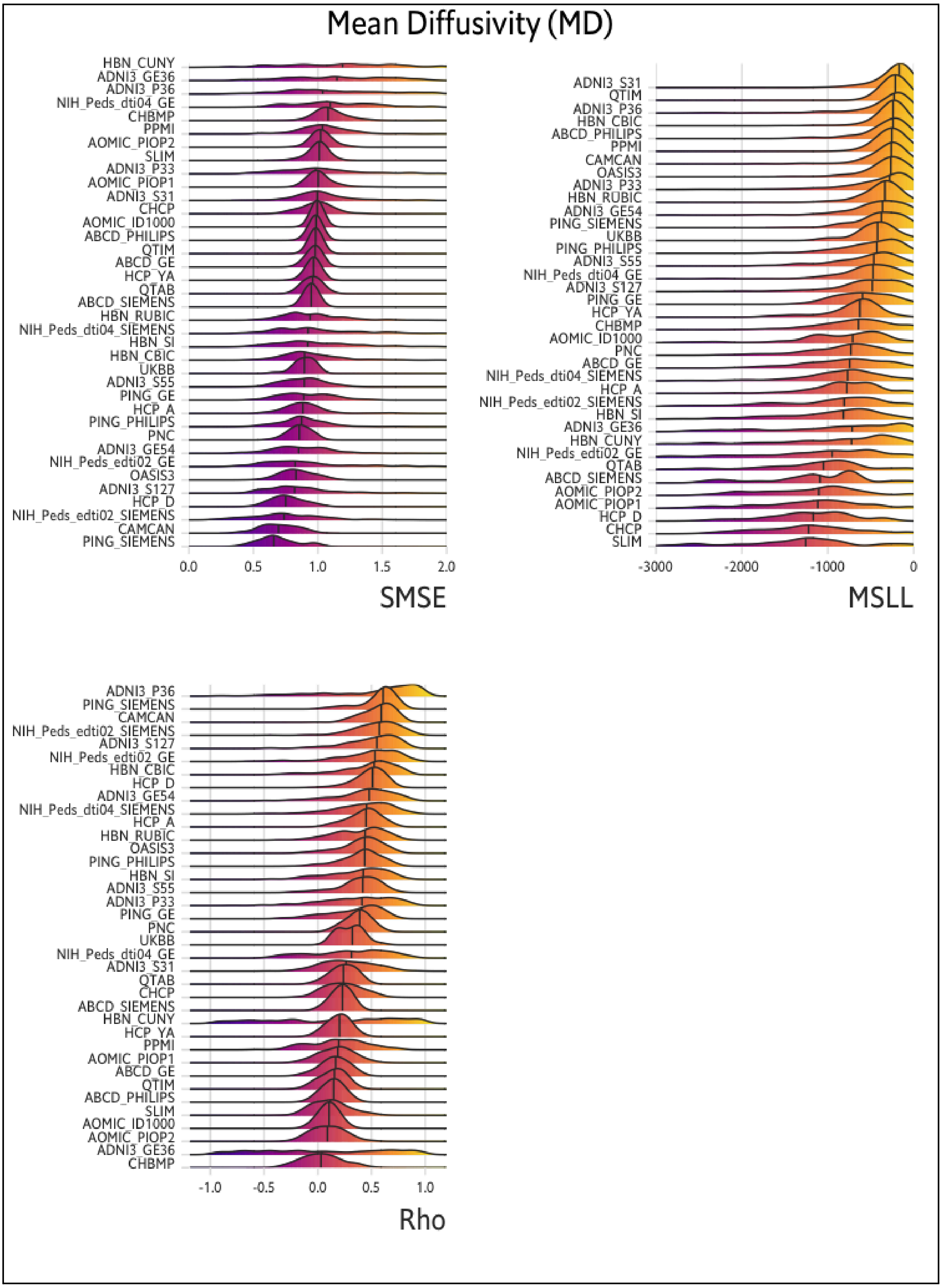

Figure S11.

**Model Diagnostics per site.** Model diagnostics metrics (SMSE, MSLL, Rho) for Axial Diffusivity (AD) per site. (37 sites, 10 experimental iterations).

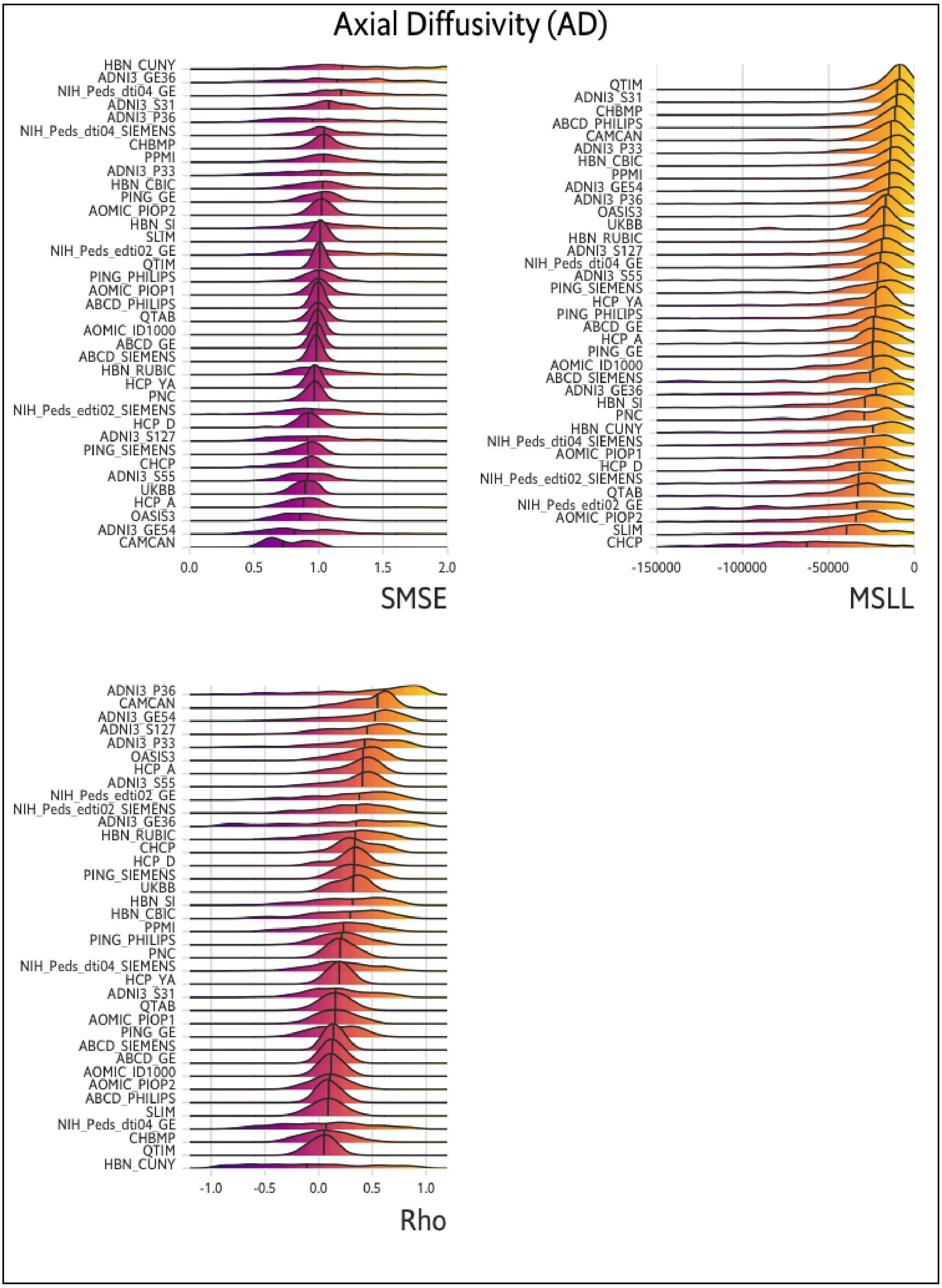

Figure S12

**Model Diagnostics per site.** Model diagnostics metrics (SMSE, MSLL, Rho) for Radial Diffusivity (RD) per site. (37 sites, 10 experimental iterations).

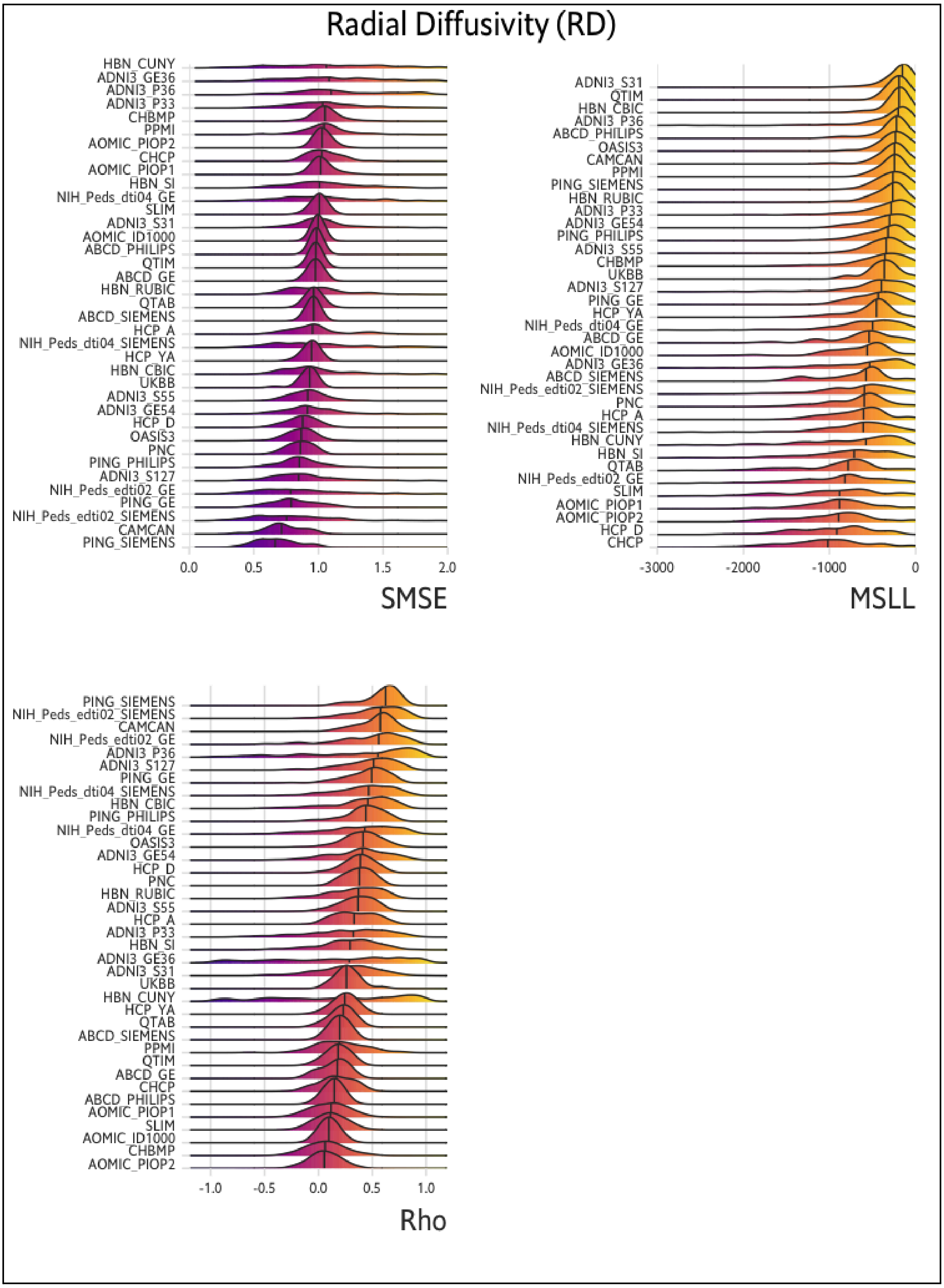

Figure S13.

**QQ_plots for Fractional Anisotropy.** QQ_plot for each WM region of the test subjects for all 10 experimental repetitions run at training. Z-scores are derived from the test set subjects (n=10,917).

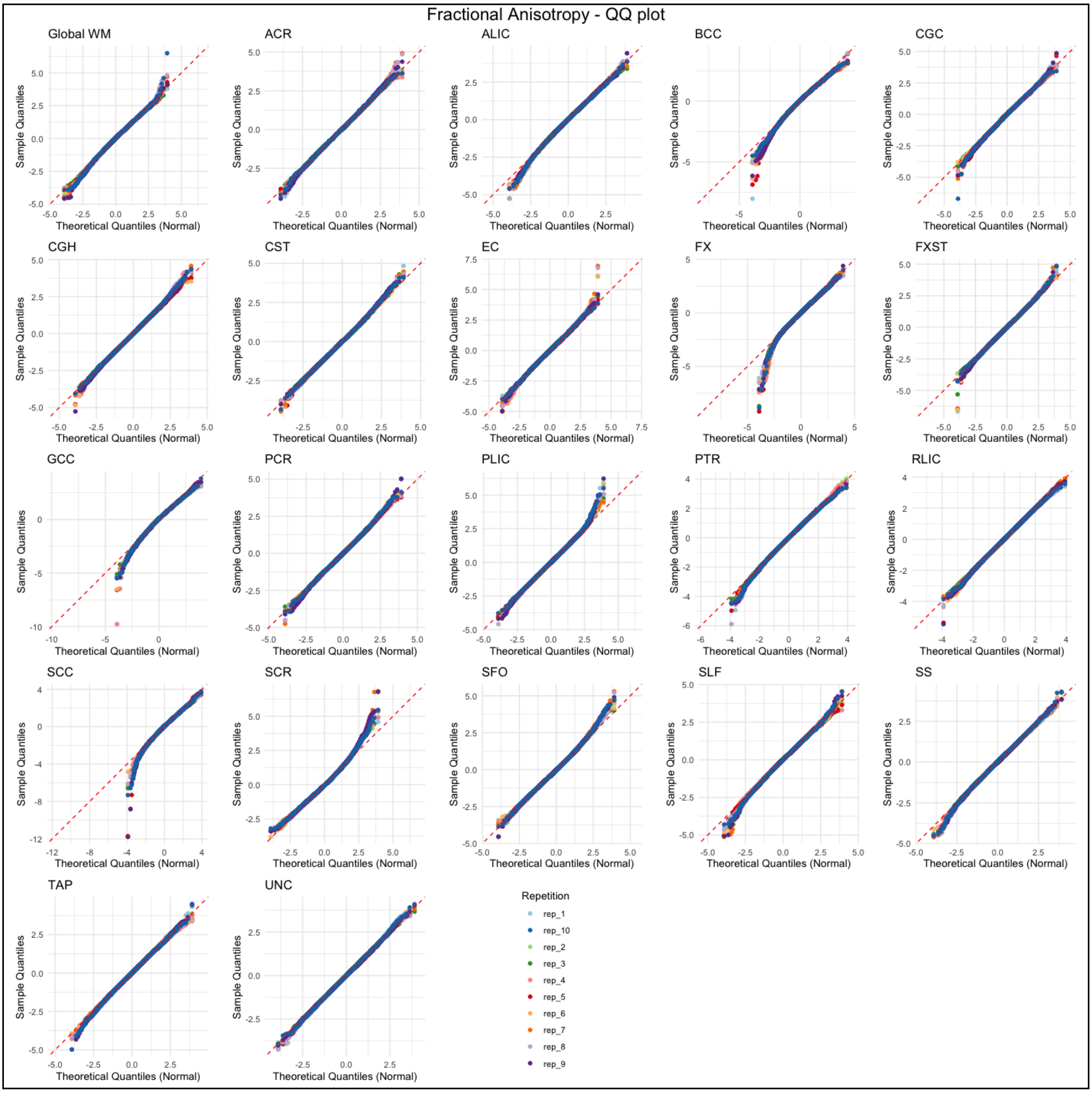

Figure S14.

**QQ_plots for Mean Diffusivity**. QQ_plot for each WM region of the test subjects for all 10 experimental repetitions run at training. Z-scores are derived from the test set subjects (n=10,917).

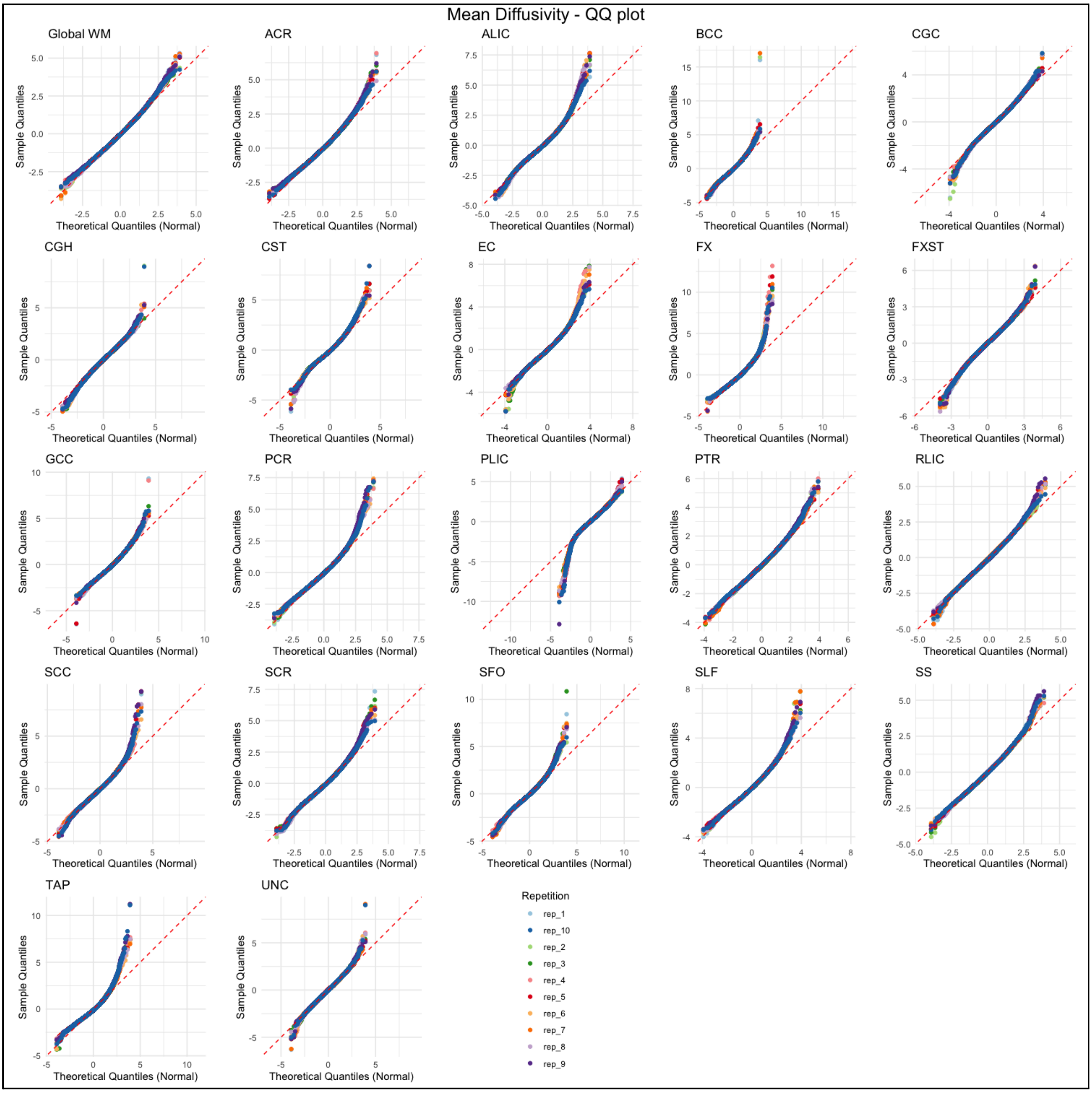

Figure S15.

**QQ_plots for Axial Diffusivity.** QQ_plot for each WM region of the test subjects for all 10 experimental repetitions run at training. Z-scores are derived from the test set subjects (n=10,917).

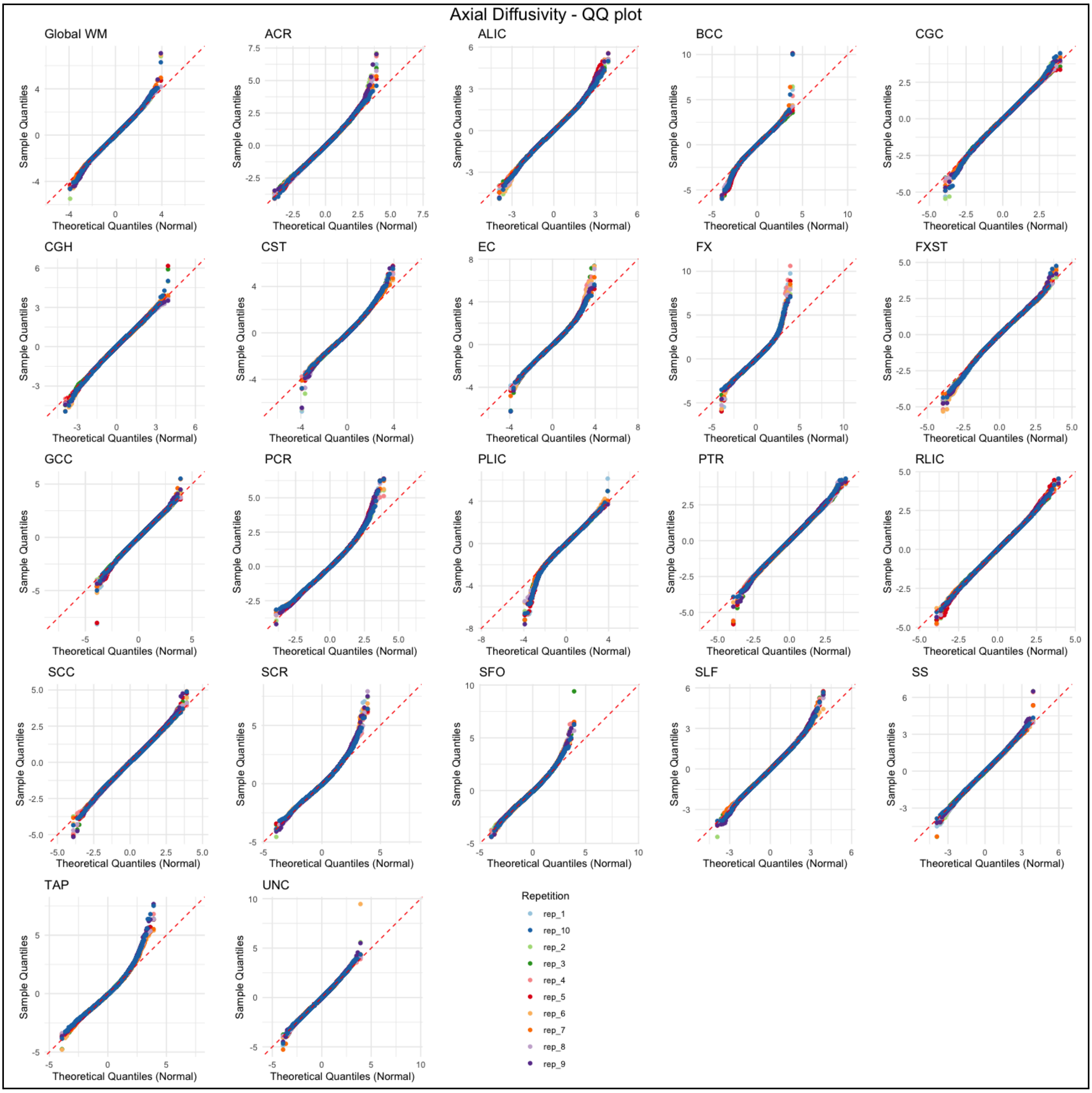

Figure S16.

**QQ_plots for Radial Diffusivity.** QQ_plot for each WM region of the test subjects for all 10 experimental repetitions run at training. Z-scores are derived from the test set subjects (n=10,917).

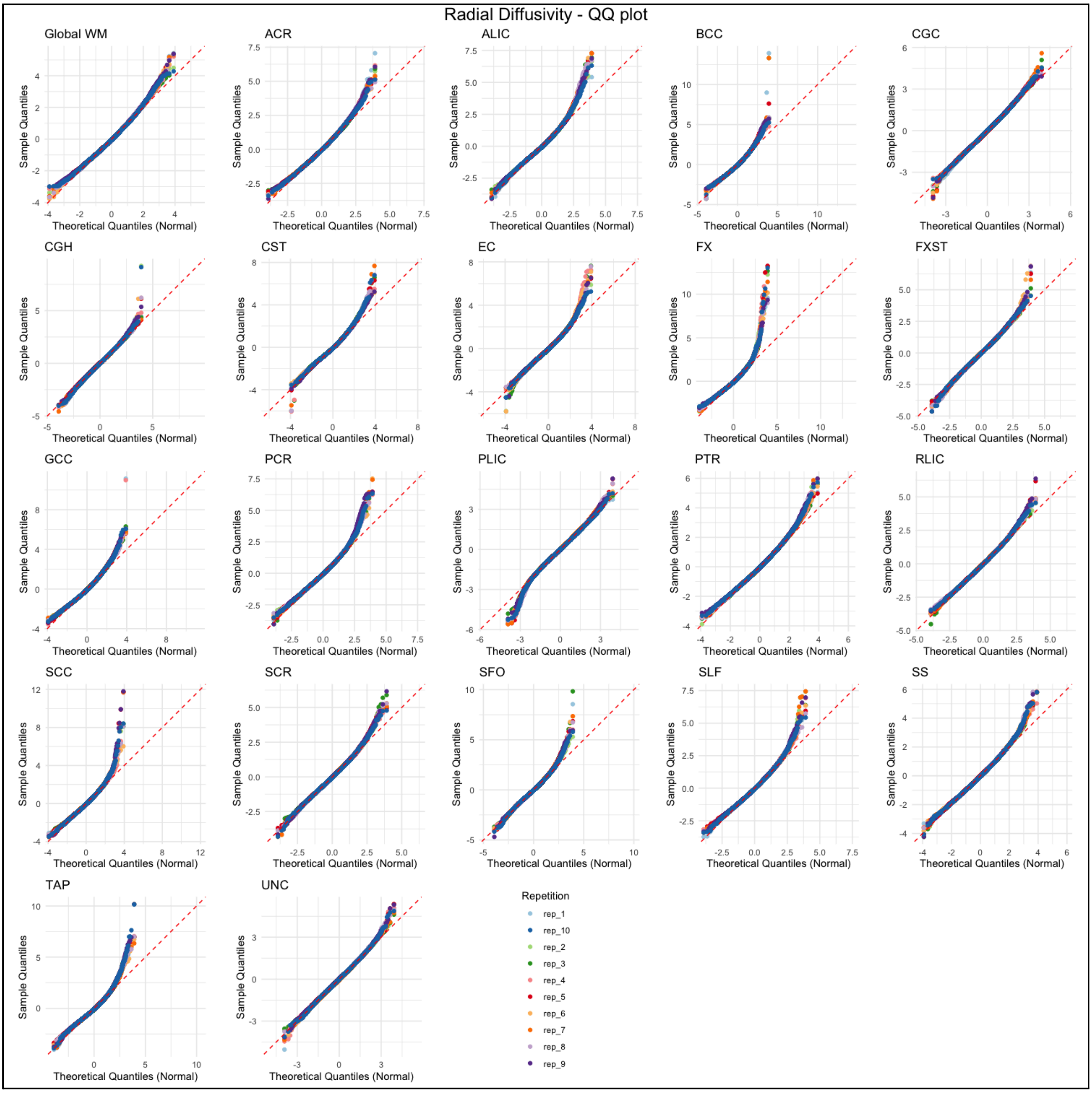

Figure S17.

**Gain-predicts-loss:** Null distributions of generated "spun" Spearman correlations and corresponding permutation p-values, and the correlation coefficients obtained from the empirical correlation between the percent change of FA and MD from 4 years to peak and the percent change from peak to 91 years. Results were significant at $p < 0.05$ under a one-sided spin permutation test (95th percentile of the null distribution).

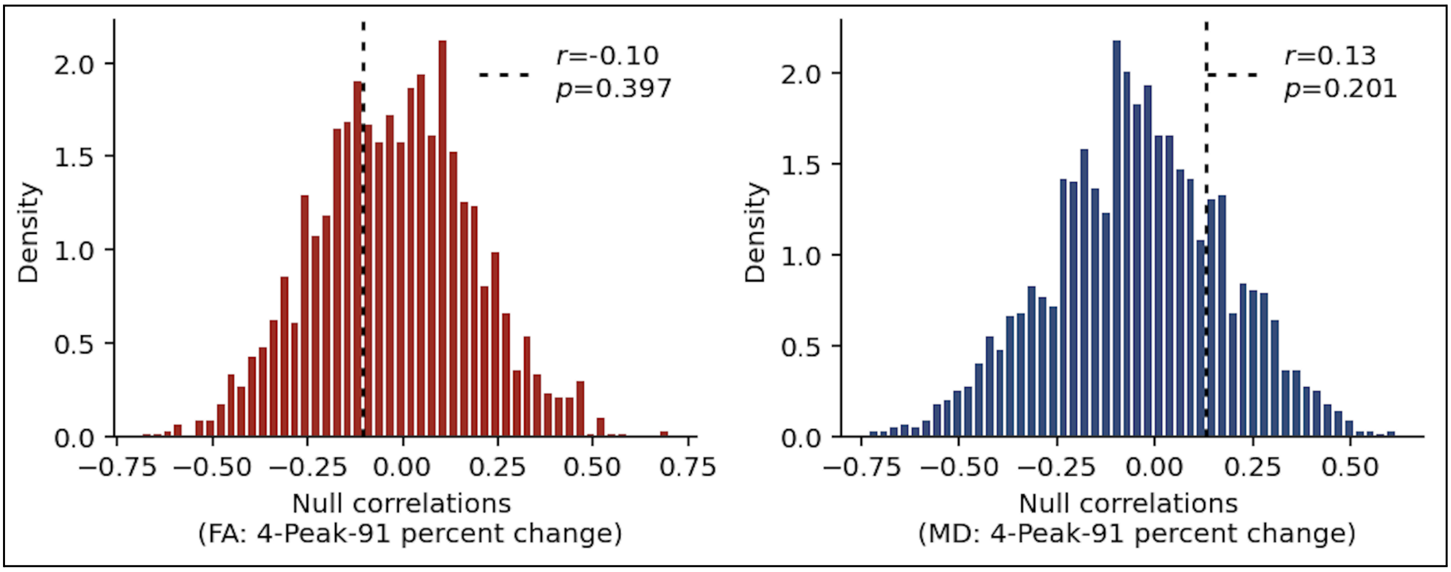


Figure S18.

**Gain-predicts-loss:** Null distributions of generated "spun" Spearman correlations and corresponding permutation p-values, and the correlation coefficients obtained from the empirical correlation between the percent change of AD and RD from 4 years to peak and the percent change from peak to 91 years. Results were significant at $p < 0.05$ under a one-sided spin permutation test (95th percentile of the null distribution).

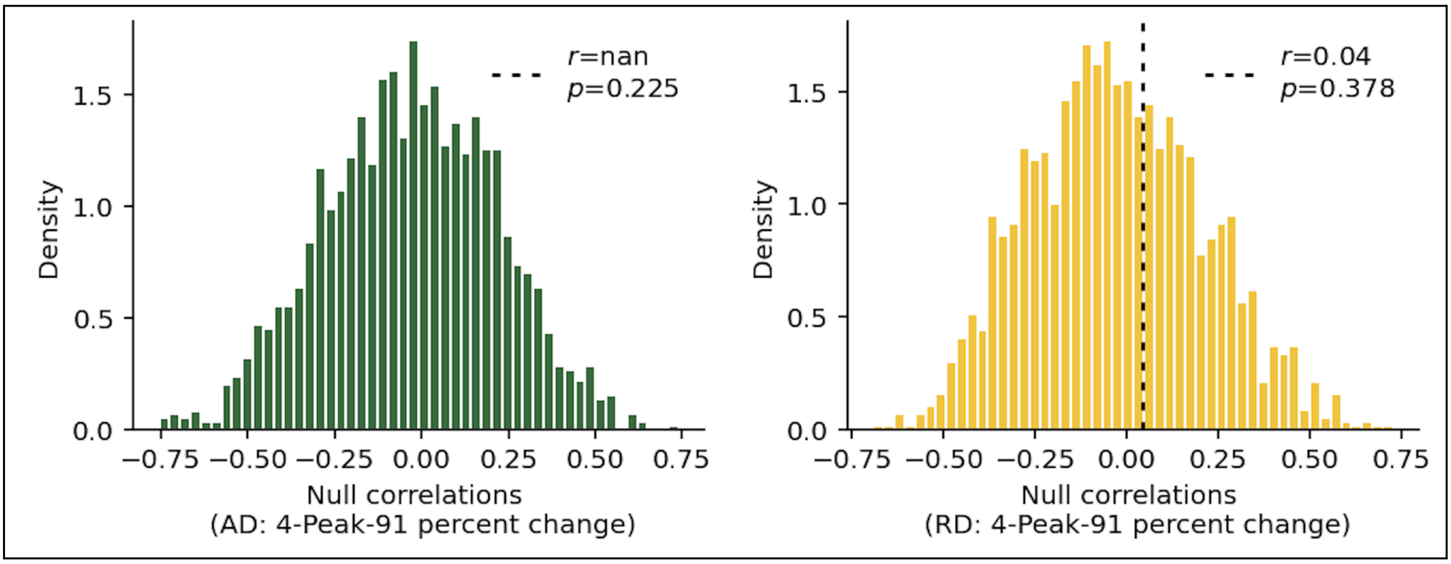

Figure S19.

**Last-in, first-out hypothesis for FA:** Null distributions of generated “spun” Spearman correlations and corresponding permutation p-values, and correlation coefficients obtained from the empirical correlation between the age of peak of FA and the percent change between 4 to 14 years, 75 to 84 years and 85 to 91 years. Results were significant at p < 0.05 under a one-sided spin permutation test (95th percentile of the null distribution).

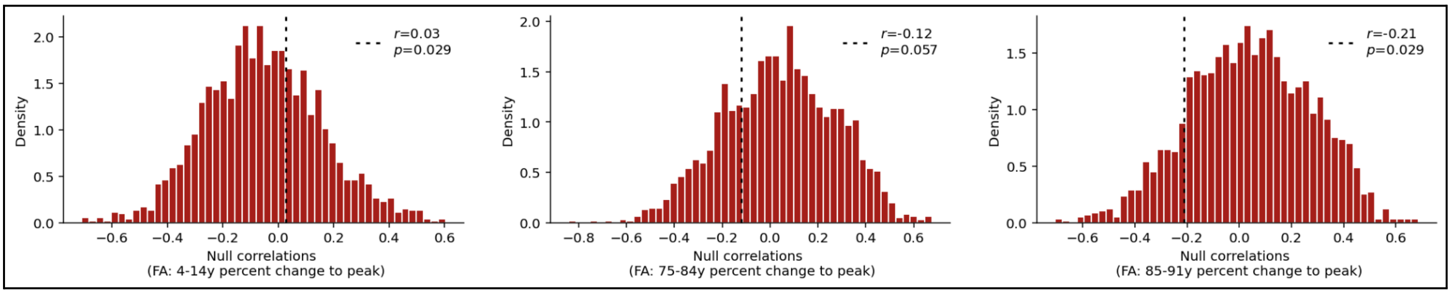


Figure S20.

**Last-in, first-out hypothesis for MD:** Null distributions of generated “spun” Spearman correlations and corresponding permutation p-values, and the correlation coefficients obtained from the empirical correlation between the age of peak of MD and the percent change between 4 to 14 years, 75 to 84 years and 85 to 91 years. Results were significant at p < 0.05 under a one-sided spin permutation test (95th percentile of the null distribution).

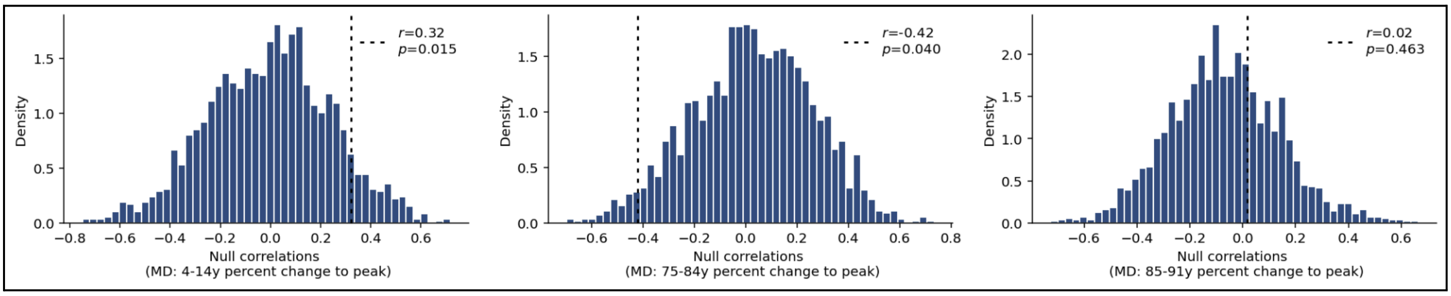


Figure S21.

**Last-in, first-out hypothesis for AD:** Null distributions of generated “spun” Spearman correlations and corresponding permutation p-values, and the correlation coefficients obtained from the empirical correlation between the age of peak of AD and the percent change between 4 to 14 years, 75 to 84 years and 85 to 91 years. Results were significant at p < 0.05 under a one-sided spin permutation test (95th percentile of the null distribution).

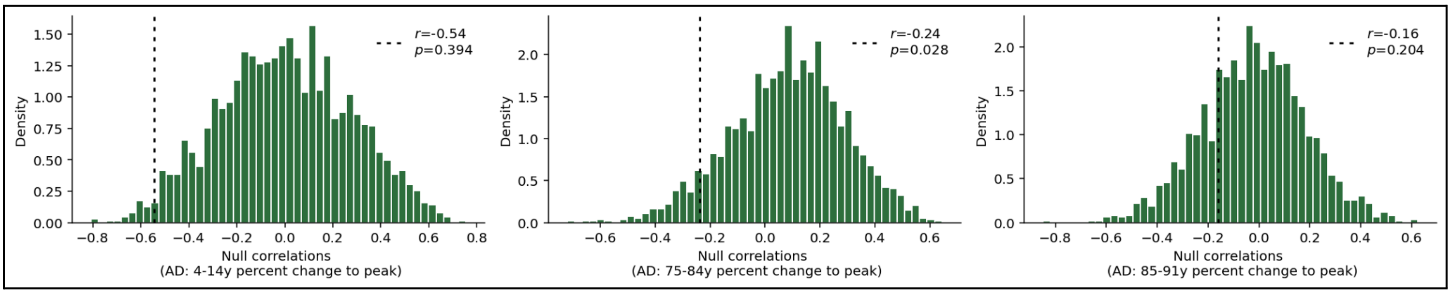

Figure S22.

**Last-in, first-out hypothesis for RD:** Null distributions of generated “spun” Spearman correlations and corresponding permutation p-values, and the correlation coefficients obtained from the empirical correlation between the age of peak of RD and the percent change between 4 to 14 years, 75 to 84 years and 85 to 91 years. Results were significant at $p < 0.05$ under a one-sided spin permutation test (95th percentile of the null distribution).

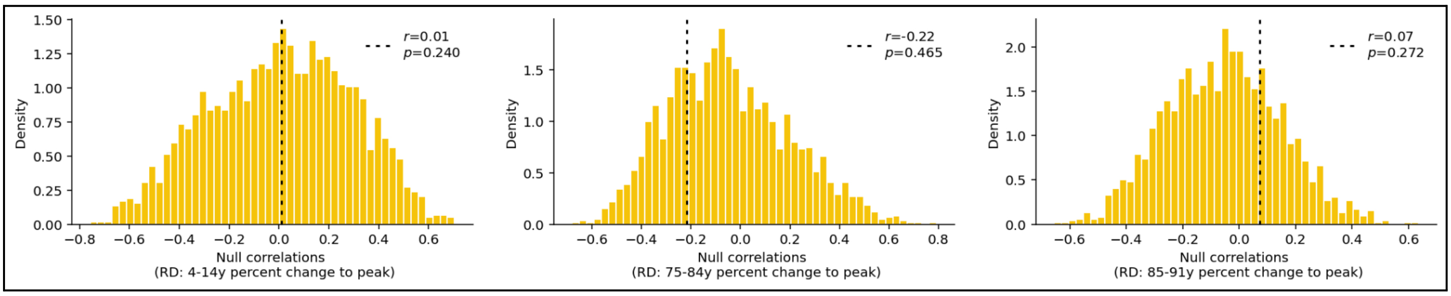

Figure S23.

**Manual QC:** Results of the t-tests comparing the kurtosis values of the Z-scores between manually and non-manually QCed datasets across 10 experimental iterations for FA, for each ROI. Multiple comparisons correction: FDR. Box: 25th - 75th percentile (IQR: Interquartile Range), Center Line: Median, Whiskers: Q1/Q3±1.5xIQR.

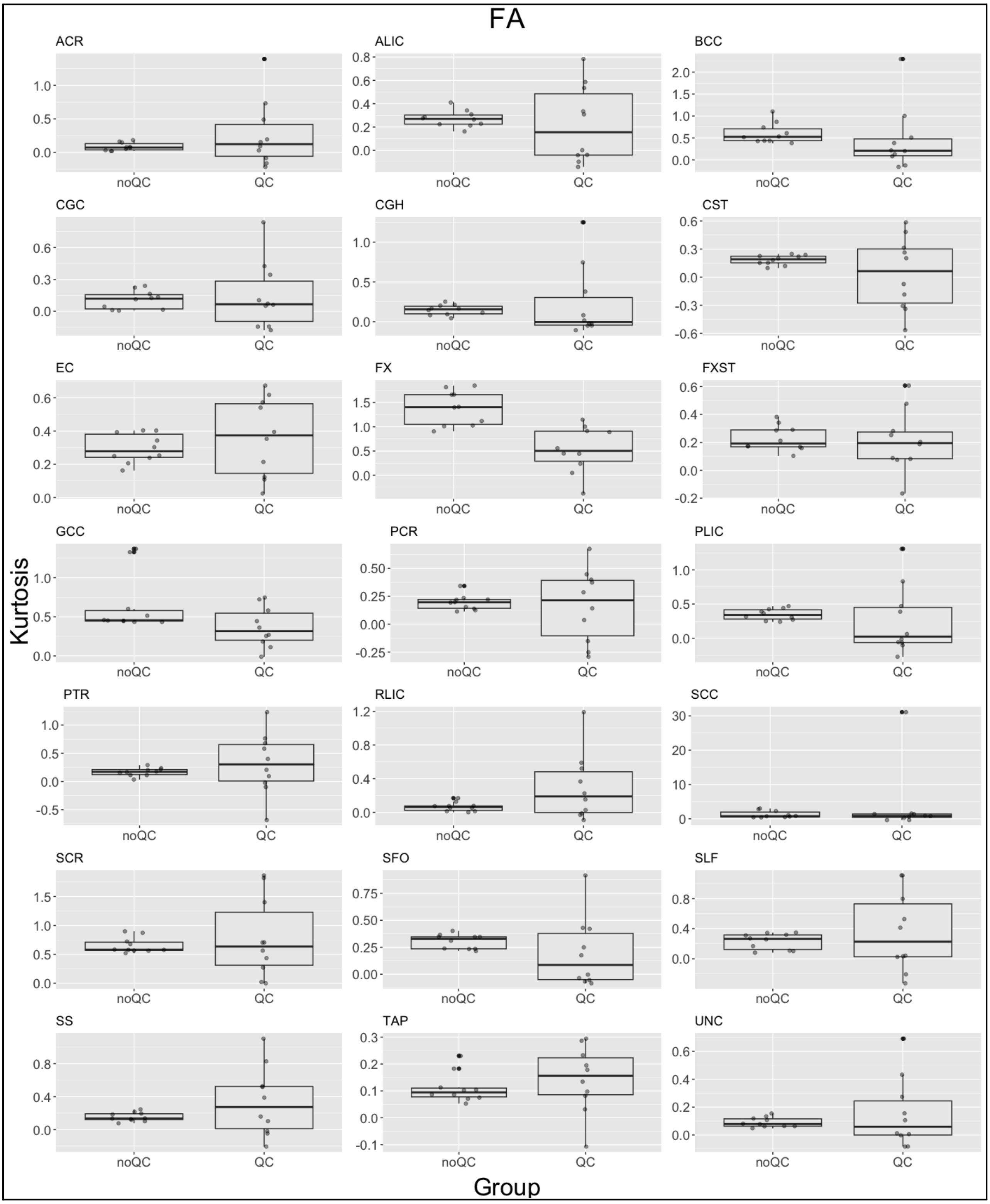

Figure S24.

**Manual QC:** Results of the t-tests comparing the kurtosis values of the Z-scores between manually and non-manually QCed datasets across 10 experimental iterations for MD, for each ROI. Multiple comparisons correction: FDR. Box: 25th - 75th percentile (IQR: Interquartile Range), Center Line: Median, Whiskers: Q1/Q3±1.5xIQR.

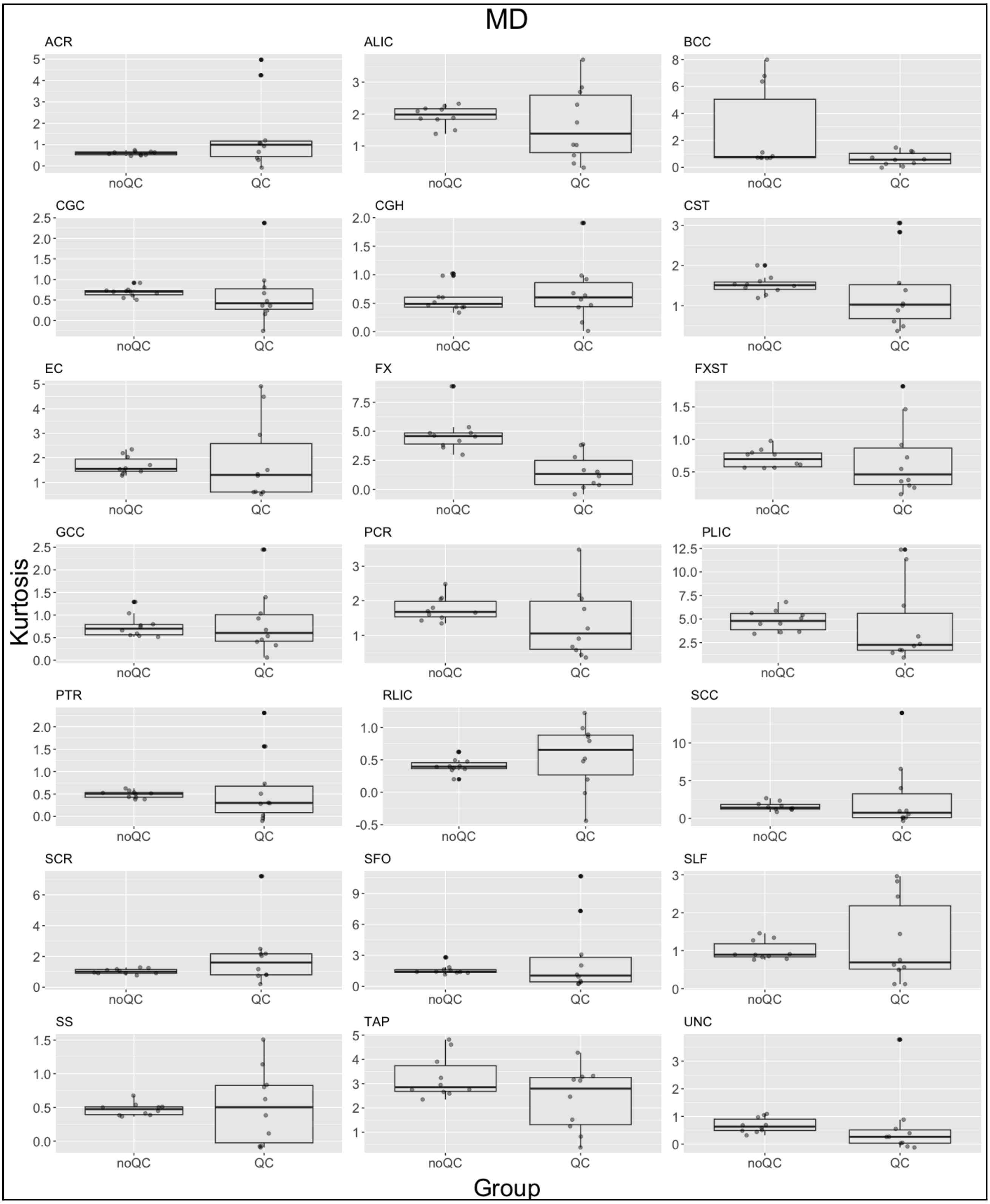

Figure S25.

**Manual QC:** Results of the t-tests comparing the kurtosis values of the Z-scores between manually and non-manually QCed datasets across 10 experimental iterations for AD, for each ROI. Multiple comparisons correction: FDR. Box: 25th - 75th percentile (IQR: Interquartile Range), Center Line: Median, Whiskers: Q1/Q3±1.5xIQR.

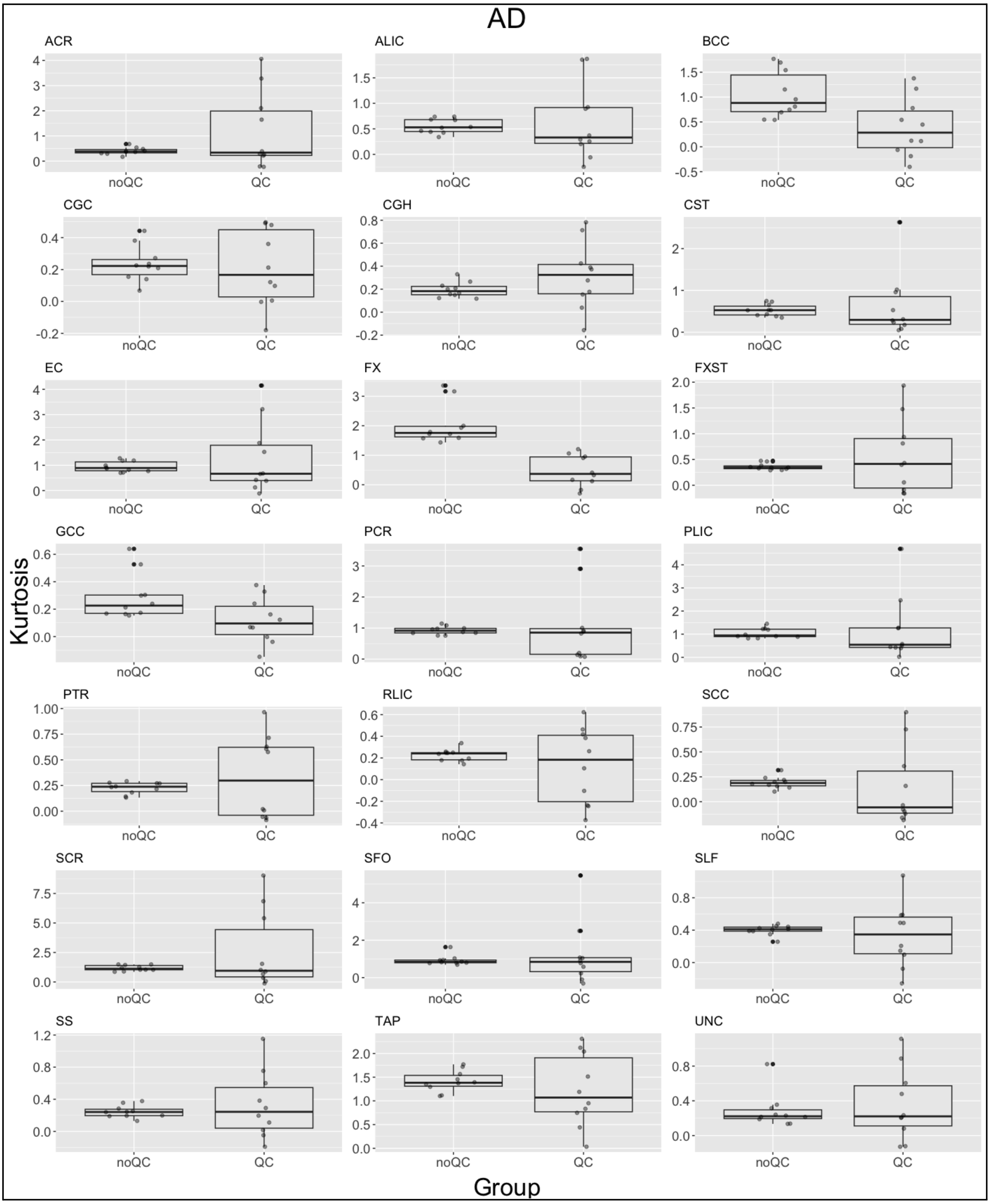

Figure S26.

**Manual QC:** Results of the t-tests comparing the kurtosis values of the Z-scores between manually and non-manually QCed datasets across 10 experimental iterations for RD, for each ROI. Multiple comparisons correction: FDR. Box: 25th - 75th percentile (IQR: Interquartile Range), Center Line: Median, Whiskers: Q1/Q3±1.5xIQR.

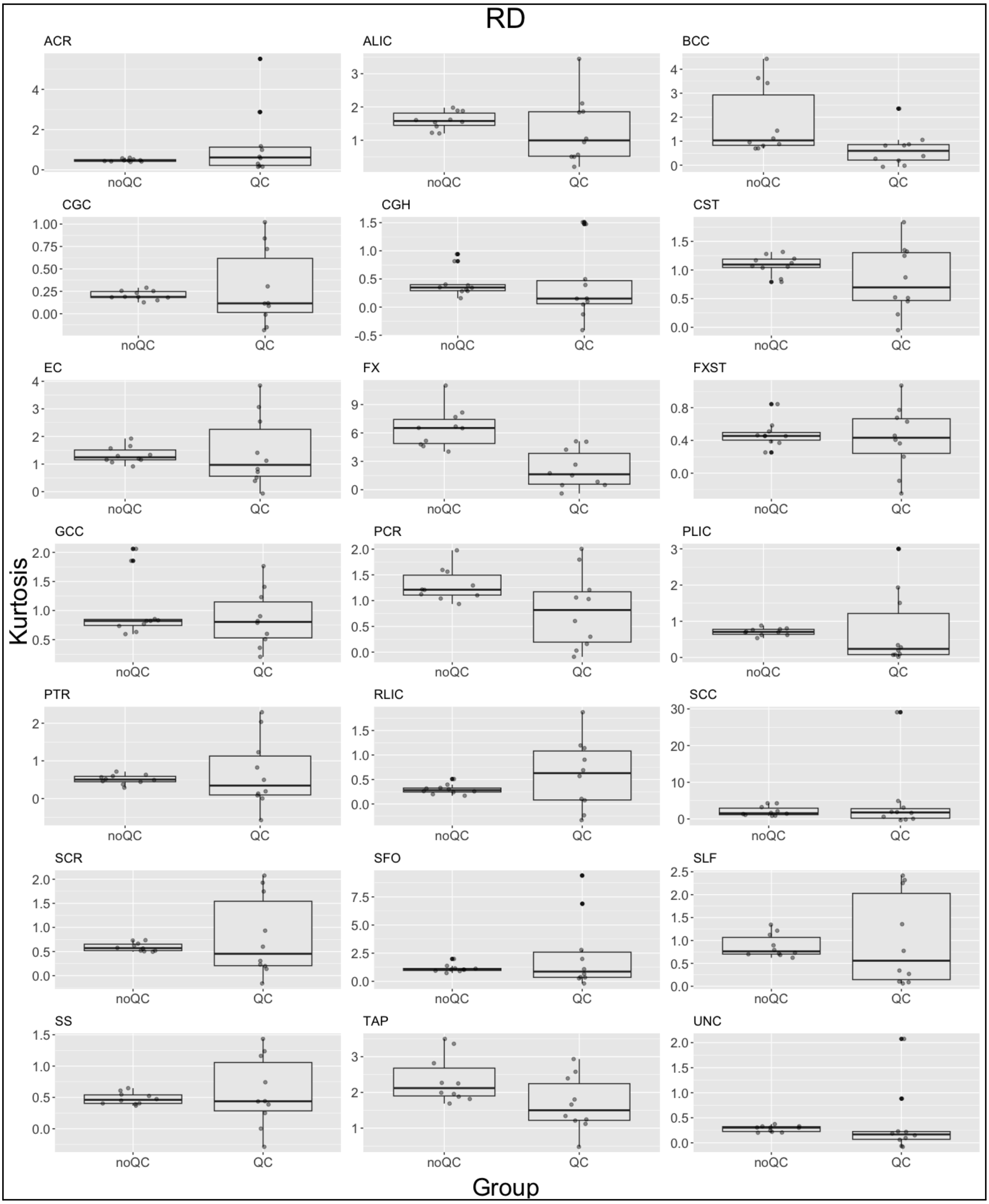

Table S1

List of full names for each of the regions of interest of the white matter according to the JHU-ENIGMA-DTI template space. There are 21 white matter regions and the “Global-WM” (including all WM in a single metric for a total of 22 total regions of interest included in the analyses.

| Acronym | Region of Interest (ROI) |
|---|---|
| Global-WM | Mean across the entire ENIGMA-DTI white matter skeleton. |
| ACR | Anterior corona radiata |
| ALIC | Anterior limb of internal capsule |
| BCC | Body of corpus callosum |
| CGC | Cingulum of the cingulate gyrus |
| CGH | Cingulum of the hippocampus |
| CST | Cortico-spinal tract |
| EC | External/extreme capsule |
| FX | Fornix |
| FXST | Crus of the fornix/stria terminalis |
| GCC | Genu of the corpus callosum |
| PCR | Posterior corona radiata |
| PLIC | Posterior limb of internal capsule |
| PTR | Posterior thalamic radiation |
| RLIC | Retrolenticular part of internal capsule |
| SCC | Splenium of the corpus callosum |
| SCR | Superior corona radiata |
| SFO | Superior fronto-occipital fasciculus |
| SLF | Superior longitudinal fasciculus |
| SS | Sagittal stratum |
| TAP | Tapetum of the corpus callosum |
| UNC | Uncinate fasciculus |

Table S2.

Comparison of age of maturation between DTI metrics (peak for FA, and minimum for MD, AD, RD). Method: repeated measures ANOVA design using a percentile bootstrap method on trimmed means (R package *WRS*[1], function wmcppb). Tests the hypothesis that the difference scores of DTI metrics across ROIs do not differ and are all equal to zero (Omnibus test). Degrees of freedom are not applicable to empirical bootstrap distributions. p-values were corrected for multiple comparisons with Hochberg's method.

| | critical p-value | CI (lower) | CI (upper) | psi.hat |
|---|---|---|---|---|
| FA vs. MD | 0.010 | -17.357143 | -9.142857 | -13.000 |
| FA vs. AD | 0.016 | -27.071429 | -10.071429 | -20.142 |
| FA vs. RD | 0.012 | -11.214286 | -4.928571 | -7.142 |
| MD vs. AD | 0.050 | -10.071429 | -3.071429 | -6.857 |
| MD vs. RD | 0.008 | 3.642857 | 6.714286 | 5.071 |
| AD vs. RD | 0.025 | 8.071429 | 16.642857 | 12.642 |

Table S3.

Weighted Least Square (WLS) regression results for comparing the original HBR model and model 1. Using WLS regression with site-clustered robust standard errors, we found no significant global differences in SMSE, Rho and MSLL (two-sided Wald tests, all p>0.05). Two-sided tests were used, as there was no a priori assumption that harmonization would necessarily improve or degrade performance for all metrics. No multiple comparison correction was required, because a single global hypothesis was tested per evaluation metric. mean_delta=mean difference of score across ROIs, ci_low=lower confidence interval, ci_high=higher confidence interval.

| | | **mean_delta** | **ci_low** | **ci_high** | **z** | **p** |
|---|---|---|---|---|---|---|
| FA | SMSE Δ | -0.0009 | -0.0041 | 0.0024 | -0.5230 | 0.6010 |
| | Rho Δ | -0.0033 | -0.0088 | 0.0022 | -1.1807 | 0.2377 |
| | MSLL Δ | 29.9303 | -23.1695 | 83.0301 | 1.1048 | 0.2693 |
| MD | SMSE Δ | -0.0074 | -0.0173 | 0.0024 | -1.4757 | 0.1400 |
| | Rho Δ | 0.0032 | -0.0172 | 0.0236 | 0.3085 | 0.7577 |
| | MSLL Δ | 48.2450 | -75.5843 | 172.0743 | 0.7636 | 0.4451 |
| AD | SMSE Δ | -0.0015 | -0.0040 | 0.0010 | -1.1720 | 0.2412 |
| | Rho Δ | 0.0002 | -0.0038 | 0.0043 | 0.1047 | 0.9166 |
| | MSLL Δ | 873.1640 | -913.8350 | 2,660.1629 | 0.9577 | 0.3382 |
| RD | SMSE Δ | -0.0077 | -0.0159 | 0.0004 | -1.8518 | 0.0641 |
| | Rho Δ | 0.0022 | -0.0171 | 0.0214 | 0.2222 | 0.8242 |
| | MSLL Δ | 41.2503 | -59.9701 | 142.4707 | 0.7987 | 0.4244 |

Table S4.

Weighted Least Square (WLS) regression results for comparing the original HBR model and model 2. Using WLS regression with site-clustered robust standard errors, we found no significant global differences in SMSE, Rho and MSLL (two-sided Wald tests, all p>0.05). Two-sided tests were used, as there was no a priori assumption that harmonization would necessarily improve or degrade performance for all metrics. No multiple comparison correction was required, because a single global hypothesis was tested per evaluation metric. mean_delta=mean difference of score across ROIs, ci_low=lower confidence interval, ci_high=higher confidence interval.

| | | mean_delta | ci_low | ci_high | z | p |
|---|---|---|---|---|---|---|
| FA | SMSE Δ | -0.0024 | -0.0069 | 0.0021 | -1.0338 | 0.3012 |
| | Rho Δ | 0.0001 | -0.0058 | 0.0060 | 0.0227 | 0.9819 |
| | MSLL Δ | 29.9294 | -23.1702 | 83.0290 | 1.1047 | 0.2693 |
| MD | SMSE Δ | -0.0065 | -0.0173 | 0.0044 | -1.1709 | 0.2420 |
| | Rho Δ | -0.0014 | -0.0272 | 0.0245 | -0.1043 | 0.9170 |
| | MSLL Δ | 48.2456 | -75.5837 | 172.0750 | 0.7636 | 0.4450 |
| AD | SMSE Δ | -0.0006 | -0.0035 | 0.0022 | -0.4343 | 0.6641 |
| | Rho Δ | -0.0019 | -0.0092 | 0.0053 | -0.5235 | 0.6006 |
| | MSLL Δ | 873.1642 | -913.8348 | 2,660.1632 | 0.9577 | 0.3382 |
| RD | SMSE Δ | -0.0081 | -0.0166 | 0.0003 | -1.8879 | 0.0590 |
| | Rho Δ | 0.0024 | -0.0177 | 0.0226 | 0.2348 | 0.8144 |
| | MSLL Δ | 41.2503 | -59.9704 | 142.4709 | 0.7987 | 0.4244 |

Table S5

Residual site effects for the LMM (Linear Mixed Model) for the Z-scores of FA. We used a joint Wald test to evaluate if all site coefficients were equal to zero. We corrected the resulting p-values across ROIs using FDR. For all ROIs, the number of sites is 37, and the number of subjects is 48,668.

| **roi_id** | **p_raw** | **p_fdr** |
|---|---|---|
| ACR | 1.0000000 | 1.0000000 |
| ALIC | 0.9999997 | 1.0000000 |
| Average | 1.0000000 | 1.0000000 |
| BCC | 0.9999370 | 1.0000000 |
| CGC | 1.0000000 | 1.0000000 |
| CGH | 0.9999995 | 1.0000000 |
| CST | 1.0000000 | 1.0000000 |
| EC | 1.0000000 | 1.0000000 |
| FX | 1.0000000 | 1.0000000 |
| FXST | 0.9999988 | 1.0000000 |
| GCC | 0.9999979 | 1.0000000 |
| PCR | 1.0000000 | 1.0000000 |
| PLIC | 1.0000000 | 1.0000000 |
| PTR | 0.9999998 | 1.0000000 |
| RLIC | 1.0000000 | 1.0000000 |
| SCC | 0.9999904 | 1.0000000 |
| SCR | 1.0000000 | 1.0000000 |
| SFO | 0.9999999 | 1.0000000 |
| SLF | 0.9999994 | 1.0000000 |
| SS | 1.0000000 | 1.0000000 |
| TAP | 0.9999681 | 1.0000000 |
| UNC | 1.0000000 | 1.0000000 |

Table S6

Residual site effects for the LMM (Linear Mixed Model) for the Z-scores of MD. We used a joint Wald test to evaluate if all site coefficients were equal to zero. We corrected the resulting p-values across ROIs using FDR. For all ROIs, the number of sites is 37, and the number of subjects is 48,668.

| **roi_id** | **p_raw** | **p_fdr** |
|---|---|---|
| TAP | 0.98212856 | 0.99999998 |
| ALIC | 0.98945941 | 0.99999998 |
| SCC | 0.99884178 | 0.99999998 |
| SFO | 0.99971873 | 0.99999998 |
| EC | 0.99984165 | 0.99999998 |
| BCC | 0.99986008 | 0.99999998 |
| UNC | 0.99990202 | 0.99999998 |
| PCR | 0.99991092 | 0.99999998 |
| SCR | 0.99998027 | 0.99999998 |
| FX | 0.99998524 | 0.99999998 |
| FXST | 0.99998782 | 0.99999998 |
| GCC | 0.99999297 | 0.99999998 |
| PTR | 0.99999473 | 0.99999998 |
| ACR | 0.99999489 | 0.99999998 |
| SLF | 0.99999491 | 0.99999998 |
| Average | 0.99999861 | 0.99999998 |
| CGC | 0.99999892 | 0.99999998 |
| PLIC | 0.99999954 | 0.99999998 |
| CST | 0.99999962 | 0.99999998 |
| RLIC | 0.99999967 | 0.99999998 |
| CGH | 0.99999988 | 0.99999998 |
| SS | 0.99999998 | 0.99999998 |

Table S7

Residual site effects for the LMM (Linear Mixed Model) for the Z-scores of RD. We used a joint Wald test to evaluate if all site coefficients were equal to zero. We corrected the resulting p-values across ROIs using FDR. For all ROIs, the number of sites is 37, and the number of subjects is 48,668.

| **roi_id** | **p_raw** | **p_fdr** |
|---|---|---|
| TAP | 0.98966554 | 1 |
| ALIC | 0.99510668 | 1 |
| BCC | 0.99884173 | 1 |
| SCC | 0.9998607 | 1 |
| GCC | 0.99995684 | 1 |
| FX | 0.99996411 | 1 |
| ACR | 0.99999278 | 1 |
| PCR | 0.99999514 | 1 |
| SLF | 0.99999526 | 1 |
| EC | 0.99999634 | 1 |
| SFO | 0.99999634 | 1 |
| UNC | 0.99999667 | 1 |
| PTR | 0.99999944 | 1 |
| FXST | 0.99999976 | 1 |
| SCR | 0.99999983 | 1 |
| Average | 0.99999991 | 1 |
| PLIC | 0.99999996 | 1 |
| CST | 0.99999996 | 1 |
| CGC | 0.99999998 | 1 |
| RLIC | 1 | 1 |
| CGH | 1 | 1 |
| SS | 1 | 1 |

Table S8

Residual site effects for the LMM (Linear Mixed Model) for the Z-scores of AD. We used a joint Wald test to evaluate if all site coefficients were equal to zero. We corrected the resulting p-values across ROIs using FDR. For all ROIs, the number of sites is 37, and the number of subjects is 48,668.

| **roi_id** | **p_raw** | **p_fdr** |
|---|---|---|
| TAP | 0.99986892 | 0.99999999 |
| SLF | 0.99988829 | 0.99999999 |
| SFO | 0.99991711 | 0.99999999 |
| SCC | 0.99996602 | 0.99999999 |
| FXST | 0.99996853 | 0.99999999 |
| EC | 0.99998205 | 0.99999999 |
| PTR | 0.99998212 | 0.99999999 |
| Average | 0.9999841 | 0.99999999 |
| PCR | 0.99998571 | 0.99999999 |
| RLIC | 0.99999571 | 0.99999999 |
| CGH | 0.99999635 | 0.99999999 |
| SCR | 0.99999638 | 0.99999999 |
| ALIC | 0.99999834 | 0.99999999 |
| CST | 0.99999878 | 0.99999999 |
| UNC | 0.99999906 | 0.99999999 |
| SS | 0.99999923 | 0.99999999 |
| FX | 0.99999943 | 0.99999999 |
| ACR | 0.99999987 | 0.99999999 |
| BCC | 0.99999989 | 0.99999999 |
| CGC | 0.99999996 | 0.99999999 |
| PLIC | 0.99999999 | 0.99999999 |
| GCC | 0.99999999 | 0.99999999 |

Table S9.

**Areas Under the ROC Curve (AUC)** for all four DTI metrics for **mild cognitive impairment (MCI)**. Numbers indicate the average AUC (across 10 experimental repetitions) and number of passing tests after multiple comparisons correction (FDR_sum) for each DTI metric per ROI. Yellow cells indicate the ROIs that passed multiple comparison testing (FDR) in at least 9 out of 10 experimental repetitions.

| ROI | FA | | MD | | RD | | AD | |
|---|---|---|---|---|---|---|---|---|
| | FDR_sum | AUC | FDR_sum | AUC | FDR_sum | AUC | FDR_sum | AUC |
| ACR | 7 | 0.54 | 7 | 0.54 | 0 | 0.52 | 8 | 0.54 |
| ALIC | 9 | 0.55 | 0 | 0.48 | 0 | 0.51 | 0 | 0.53 |
| BCC | 10 | 0.57 | 10 | 0.58 | 10 | 0.58 | 10 | 0.59 |
| CGC | 10 | 0.56 | 10 | 0.55 | 9 | 0.55 | 8 | 0.54 |
| CGH | 10 | 0.56 | 9 | 0.55 | 10 | 0.56 | 0 | 0.48 |
| CST | 0 | 0.47 | 0 | 0.50 | 0 | 0.50 | 0 | 0.49 |
| EC | 10 | 0.56 | 0 | 0.49 | 0 | 0.51 | 0 | 0.52 |
| FX | 10 | 0.58 | 10 | 0.58 | 10 | 0.59 | 6 | 0.54 |
| FXST | 4 | 0.53 | 0 | 0.51 | 0 | 0.51 | 0 | 0.52 |
| GCC | 8 | 0.55 | 10 | 0.56 | 8 | 0.55 | 10 | 0.58 |
| UNC | 2 | 0.53 | 10 | 0.57 | 6 | 0.54 | 10 | 0.57 |
| PCR | 0 | 0.53 | 2 | 0.53 | 0 | 0.51 | 10 | 0.57 |
| PLIC | 0 | 0.47 | 0 | 0.52 | 0 | 0.49 | 0 | 0.50 |
| PTR | 10 | 0.55 | 10 | 0.55 | 10 | 0.57 | 0 | 0.52 |
| RLIC | 0 | 0.52 | 10 | 0.55 | 5 | 0.54 | 3 | 0.54 |
| SCC | 10 | 0.55 | 10 | 0.60 | 10 | 0.57 | 10 | 0.56 |
| SCR | 4 | 0.53 | 7 | 0.54 | 1 | 0.53 | 10 | 0.57 |
| SFO | 3 | 0.53 | 0 | 0.50 | 0 | 0.50 | 0 | 0.52 |
| SLF | 2 | 0.53 | 10 | 0.56 | 3 | 0.54 | 10 | 0.57 |
| SS | 0 | 0.53 | 10 | 0.57 | 7 | 0.55 | 9 | 0.54 |
| TAP | 0 | 0.51 | 10 | 0.55 | 8 | 0.54 | 0 | 0.52 |

Table S10.

**Areas Under the ROC Curve (AUC)** for all four DTI metrics for **dementia**. Numbers indicate the average AUC (across 10 experimental repetitions) and number of passing tests after multiple comparisons correction (FDR_sum) for each DTI metric per ROI. Yellow cells indicate the ROIs that passed multiple comparison testing (FDR) in at least 9 out of 10 experimental repetitions.

| ROI | FA | | MD | | RD | | AD | |
|---|---|---|---|---|---|---|---|---|
| | FDR_sum | AUC | FDR_sum | AUC | FDR_sum | AUC | FDR_sum | AUC |
| ACR | 0 | 0.49 | 10 | 0.61 | 10 | 0.58 | 10 | 0.60 |
| ALIC | 0 | 0.52 | 0 | 0.53 | 0 | 0.50 | 0 | 0.52 |
| BCC | 8 | 0.57 | 10 | 0.65 | 10 | 0.63 | 10 | 0.67 |
| CGC | 10 | 0.62 | 10 | 0.64 | 10 | 0.68 | 0 | 0.51 |
| CGH | 10 | 0.65 | 10 | 0.70 | 10 | 0.74 | 0 | 0.54 |
| CST | 0 | 0.50 | 0 | 0.53 | 0 | 0.53 | 0 | 0.50 |
| EC | 0 | 0.54 | 0 | 0.48 | 0 | 0.52 | 0 | 0.51 |
| FX | 10 | 0.64 | 10 | 0.66 | 10 | 0.68 | 8 | 0.57 |
| FXST | 1 | 0.55 | 10 | 0.62 | 10 | 0.60 | 10 | 0.61 |
| GCC | 4 | 0.56 | 10 | 0.62 | 10 | 0.60 | 10 | 0.61 |
| UNC | 10 | 0.60 | 10 | 0.61 | 10 | 0.63 | 10 | 0.60 |
| PCR | 0 | 0.51 | 0 | 0.53 | 0 | 0.52 | 3 | 0.57 |
| PLIC | 0 | 0.53 | 0 | 0.53 | 0 | 0.54 | 0 | 0.52 |
| PTR | 8 | 0.58 | 10 | 0.58 | 10 | 0.61 | 0 | 0.50 |
| RLIC | 0 | 0.54 | 0 | 0.55 | 2 | 0.55 | 0 | 0.55 |
| SCC | 10 | 0.60 | 10 | 0.68 | 10 | 0.63 | 10 | 0.65 |
| SCR | 5 | 0.56 | 0 | 0.55 | 0 | 0.53 | 0 | 0.53 |
| SFO | 0 | 0.54 | 0 | 0.51 | 0 | 0.48 | 7 | 0.57 |
| SLF | 10 | 0.59 | 10 | 0.58 | 10 | 0.60 | 10 | 0.60 |
| SS | 0 | 0.52 | 10 | 0.59 | 10 | 0.59 | 0 | 0.54 |
| TAP | 0 | 0.46 | 9 | 0.57 | 0 | 0.52 | 0 | 0.55 |

Table S11.

**Extreme deviations** for all four DTI metrics for ***mild cognitive impairment (MCI)*. ** Numbers indicate the percentage of subjects with MCI (ADNI3 and OASIS3) with extreme deviations for each DTI metric per ROI. Yellow cells indicate if the ROI passed multiple comparisons correction (FDR) in at least 9 out of 10 experimental repetitions. Also see supplementary Table 2 for the AUCs.

| ROI | FA | | MD | | RD | | AD | |
|---|---|---|---|---|---|---|---|---|
| | Z < -2 | Z > 2 | Z < -2 | Z > 2 | Z < -2 | Z > 2 | Z < -2 | Z > 2 |
| ACR | 5.24 | 3.56 | 1.11 | 9.82 | 1.20 | 10.40 | 1.60 | 10.44 |
| ALIC | 5.87 | 3.42 | 0.00 | 8.89 | 0.18 | 8.09 | 0.84 | 9.24 |
| BCC | 7.78 | 4.36 | 1.51 | 11.20 | 1.64 | 9.69 | 2.58 | 8.62 |
| CGC | 6.62 | 2.89 | 2.84 | 6.04 | 2.98 | 6.80 | 3.51 | 2.67 |
| CGH | 4.44 | 3.11 | 0.44 | 6.36 | 1.33 | 10.76 | 1.11 | 3.96 |
| CST | 1.91 | 0.00 | 3.20 | 6.27 | 0.58 | 4.93 | 1.51 | 3.11 |
| EC | 5.47 | 1.73 | 0.49 | 7.38 | 0.09 | 8.58 | 0.89 | 6.53 |
| FX | 6.31 | 2.44 | 1.33 | 9.96 | 1.38 | 10.58 | 1.60 | 6.31 |
| FXST | 4.71 | 1.20 | 1.24 | 3.73 | 1.16 | 5.11 | 5.07 | 1.82 |
| GCC | 8.44 | 1.69 | 0.98 | 13.20 | 0.93 | 13.47 | 2.58 | 9.33 |
| UNC | 7.33 | 1.87 | 1.78 | 10.49 | 0.62 | 10.04 | 4.13 | 5.69 |
| PCR | 4.93 | 2.27 | 0.44 | 7.73 | 0.04 | 8.31 | 1.11 | 10.00 |
| PLIC | 1.42 | 2.40 | 0.93 | 5.64 | 1.96 | 2.31 | 0.84 | 4.67 |
| PTR | 6.44 | 2.93 | 0.93 | 7.96 | 1.56 | 10.49 | 1.73 | 4.67 |
| RLIC | 4.76 | 1.56 | 1.16 | 7.64 | 0.89 | 7.82 | 1.16 | 4.22 |
| SCC | 13.16 | 0.58 | 1.29 | 15.42 | 0.00 | 16.53 | 3.96 | 5.29 |
| SCR | 1.16 | 4.49 | 0.62 | 11.42 | 0.36 | 6.67 | 1.20 | 10.09 |
| SFO | 3.47 | 2.36 | 0.09 | 8.53 | 0.80 | 7.42 | 0.18 | 8.98 |
| SLF | 4.62 | 3.51 | 0.89 | 8.76 | 1.69 | 8.36 | 1.42 | 6.53 |
| SS | 5.38 | 1.91 | 0.49 | 10.00 | 1.02 | 8.80 | 1.47 | 6.36 |
| TAP | 1.16 | 2.49 | 0.00 | 5.82 | 0.44 | 6.44 | 1.51 | 4.62 |

Table S12.

**Extreme deviations** for all four DTI metrics for ***dementia***. Numbers indicate the percentage of subjects with dementia (ADNI3 and OASIS3) with extreme deviations for each DTI metric per ROI. Yellow cells indicate if the ROI passed multiple comparison testing (FDR) in at least 9 out of 10 experimental repetitions. Also see supplementary Table 3 for the AUCs.

| ROI | FA | | MD | | RD | | AD | |
|---|---|---|---|---|---|---|---|---|
| | Z < -2 | Z > 2 | Z < -2 | Z > 2 | Z < -2 | Z > 2 | Z < -2 | Z > 2 |
| ACR | 7.90 | 0.00 | 0.00 | 18.15 | 0.00 | 18.02 | 0.00 | 12.96 |
| ALIC | 4.44 | 2.47 | 0.12 | 7.41 | 0.00 | 5.56 | 0.00 | 10.49 |
| BCC | 11.85 | 0.12 | 0.25 | 26.91 | 0.12 | 20.12 | 2.47 | 28.52 |
| CGC | 14.94 | 0.00 | 0.00 | 17.90 | 0.00 | 20.62 | 5.80 | 2.47 |
| CGH | 20.37 | 0.00 | 0.00 | 29.88 | 0.00 | 36.67 | 3.46 | 7.28 |
| CST | 4.07 | 0.12 | 0.99 | 3.21 | 0.00 | 5.56 | 0.99 | 2.72 |
| EC | 9.01 | 2.72 | 0.00 | 8.77 | 0.00 | 8.64 | 0.00 | 7.16 |
| FX | 20.99 | 0.00 | 0.00 | 21.48 | 0.00 | 22.22 | 0.00 | 16.91 |
| FXST | 11.73 | 0.00 | 4.69 | 7.28 | 0.00 | 11.23 | 10.00 | 2.84 |
| GCC | 11.36 | 1.11 | 1.11 | 22.35 | 0.00 | 15.93 | 1.60 | 14.94 |
| UNC | 14.81 | 1.85 | 1.23 | 17.90 | 0.00 | 16.91 | 4.57 | 4.44 |
| PCR | 4.69 | 1.23 | 0.00 | 11.85 | 0.00 | 9.63 | 0.00 | 15.56 |
| PLIC | 0.74 | 7.65 | 0.00 | 4.44 | 3.95 | 3.33 | 0.00 | 7.65 |
| PTR | 12.22 | 1.73 | 2.47 | 14.32 | 1.85 | 15.68 | 3.70 | 0.74 |
| RLIC | 1.85 | 1.98 | 1.23 | 7.53 | 0.74 | 6.54 | 0.12 | 6.17 |
| SCC | 16.79 | 0.25 | 1.23 | 25.80 | 0.00 | 24.81 | 2.84 | 18.40 |
| SCR | 1.23 | 11.36 | 0.00 | 12.47 | 0.86 | 6.91 | 0.00 | 13.95 |
| SFO | 0.12 | 8.52 | 0.00 | 7.65 | 0.12 | 4.20 | 0.00 | 12.96 |
| SLF | 8.89 | 5.31 | 1.23 | 16.91 | 1.23 | 16.05 | 0.00 | 13.33 |
| SS | 6.91 | 0.37 | 1.23 | 12.22 | 0.37 | 13.83 | 1.60 | 6.05 |
| TAP | 2.96 | 2.84 | 1.23 | 6.17 | 1.23 | 4.94 | 1.11 | 5.31 |

Table S13.

Model adaptation on the **NIMHANS** dataset. **Extreme deviations** for all four DTI metrics for **mild cognitive impairment (MCI)**. Numbers indicate the percentage of subjects with MCI with extreme deviations for each DTI metric per ROI. Yellow cells indicate the ROIs that passed multiple comparison testing (FDR) in at least 9 out of 10 experimental repetitions. Also see supplementary Table 4 for the AUCs.

| ROI | FA | | MD | | RD | | AD | |
|---|---|---|---|---|---|---|---|---|
| | Z < -2 | Z > 2 | Z < -2 | Z > 2 | Z < -2 | Z > 2 | Z < -2 | Z > 2 |
| ACR | 3.26 | 1.24 | 1.12 | 3.03 | 1.24 | 3.26 | 1.35 | 4.04 |
| ALIC | 1.24 | 2.36 | 0.22 | 3.03 | 1.24 | 2.47 | 1.35 | 1.12 |
| BCC | 3.93 | 0.56 | 0.00 | 6.74 | 0.00 | 6.18 | 0.45 | 5.17 |
| CGC | 3.26 | 1.01 | 0.11 | 4.38 | 0.34 | 3.93 | 0.00 | 3.82 |
| CGH | 4.38 | 1.35 | 0.56 | 3.71 | 1.24 | 3.48 | 0.00 | 4.16 |
| CST | 0.22 | 3.37 | 0.45 | 1.57 | 0.90 | 0.90 | 0.34 | 2.02 |
| EC | 2.02 | 1.57 | 0.67 | 2.70 | 1.01 | 2.36 | 0.67 | 3.48 |
| FX | 7.08 | 0.22 | 0.00 | 6.29 | 0.00 | 8.43 | 0.00 | 3.26 |
| FXST | 7.30 | 0.79 | 0.45 | 4.04 | 0.22 | 4.61 | 1.35 | 2.81 |
| GCC | 2.81 | 1.24 | 0.45 | 5.73 | 1.12 | 5.84 | 0.11 | 4.83 |
| UNC | 3.48 | 1.12 | 0.67 | 10.56 | 1.12 | 12.47 | 0.34 | 3.03 |
| PCR | 5.17 | 4.04 | 1.57 | 5.17 | 1.46 | 5.62 | 1.57 | 6.18 |
| PLIC | 3.37 | 1.35 | 0.79 | 3.37 | 1.12 | 3.37 | 1.12 | 2.25 |
| PTR | 2.13 | 2.70 | 4.83 | 3.93 | 2.92 | 3.82 | 5.39 | 3.26 |
| RLIC | 2.25 | 1.57 | 0.00 | 3.71 | 0.11 | 3.37 | 0.00 | 3.71 |
| SCC | 2.36 | 0.00 | 0.11 | 3.71 | 0.00 | 3.60 | 0.45 | 5.17 |
| SCR | 4.27 | 6.40 | 0.22 | 3.82 | 0.79 | 3.93 | 0.79 | 6.74 |
| SFO | 2.81 | 4.83 | 3.37 | 3.37 | 2.25 | 3.37 | 3.37 | 4.04 |
| SLF | 5.28 | 2.36 | 2.81 | 5.73 | 1.80 | 5.96 | 3.48 | 4.27 |
| SS | 2.58 | 2.25 | 2.02 | 4.72 | 2.13 | 4.61 | 0.79 | 5.73 |
| TAP | 2.36 | 3.37 | 1.35 | 5.84 | 2.02 | 4.61 | 0.45 | 5.62 |

Table S14.

Results for model adaptation to the **NIMHANS** dataset. **Areas Under the ROC Curve (AUC)** for all four DTI metrics for **mild cognitive impairment (MCI)**. Numbers indicate the average AUC (across 10 experimental repetitions) and number of passing tests after multiple comparisons correction (FDR_sum) for each DTI metric per ROI. Yellow cells indicate the ROIs that passed multiple comparison testing (FDR) in at least 9 out of 10 experimental repetitions.

| ROI | FA | | | MD | | | RD | | | AD | | |
|---|---|---|---|---|---|---|---|---|---|---|---|---|
| | FDR_sum | AUC (mean/std) | | FDR_sum | AUC (mean/std) | | FDR_sum | AUC (mean/std) | | FDR_sum | AUC (mean/std) | |
| ACR | 0 | 0.42 | 0.05 | 9 | 0.64 | 0.05 | 9 | 0.61 | 0.05 | 3 | 0.57 | 0.06 |
| ALIC | 0 | 0.54 | 0.04 | 10 | 0.89 | 0.02 | 10 | 0.84 | 0.04 | 9 | 0.69 | 0.04 |
| BCC | 0 | 0.56 | 0.05 | 10 | 0.79 | 0.06 | 10 | 0.72 | 0.05 | 10 | 0.71 | 0.05 |
| CGC | 0 | 0.45 | 0.03 | 10 | 0.72 | 0.05 | 9 | 0.65 | 0.05 | 1 | 0.54 | 0.04 |
| CGH | 0 | 0.50 | 0.04 | 6 | 0.59 | 0.03 | 7 | 0.61 | 0.03 | 0 | 0.46 | 0.05 |
| CST | 0 | 0.55 | 0.02 | 10 | 0.77 | 0.06 | 10 | 0.76 | 0.05 | 10 | 0.64 | 0.02 |
| EC | 0 | 0.47 | 0.03 | 10 | 0.76 | 0.03 | 10 | 0.70 | 0.03 | 10 | 0.65 | 0.03 |
| FX | 0 | 0.55 | 0.06 | 1 | 0.52 | 0.04 | 1 | 0.54 | 0.04 | 1 | 0.51 | 0.04 |
| FXST | 0 | 0.50 | 0.04 | 10 | 0.68 | 0.05 | 7 | 0.61 | 0.05 | 6 | 0.59 | 0.06 |
| GCC | 0 | 0.47 | 0.04 | 9 | 0.68 | 0.05 | 8 | 0.61 | 0.04 | 7 | 0.60 | 0.07 |
| UNC | 0 | 0.50 | 0.04 | 10 | 0.74 | 0.03 | 10 | 0.70 | 0.04 | 4 | 0.59 | 0.03 |
| PCR | 0 | 0.55 | 0.04 | 6 | 0.61 | 0.08 | 6 | 0.60 | 0.07 | 6 | 0.60 | 0.08 |
| PLIC | 2 | 0.59 | 0.04 | 10 | 0.85 | 0.03 | 10 | 0.81 | 0.04 | 10 | 0.69 | 0.04 |
| PTR | 2 | 0.62 | 0.02 | 10 | 0.72 | 0.02 | 10 | 0.71 | 0.02 | 2 | 0.55 | 0.03 |
| RLIC | 0 | 0.51 | 0.03 | 10 | 0.64 | 0.03 | 10 | 0.65 | 0.03 | 0 | 0.52 | 0.04 |
| SCC | 0 | 0.55 | 0.03 | 10 | 0.75 | 0.05 | 10 | 0.70 | 0.05 | 9 | 0.67 | 0.05 |
| SCR | 0 | 0.55 | 0.05 | 10 | 0.80 | 0.03 | 10 | 0.75 | 0.04 | 10 | 0.69 | 0.04 |
| SFO | 0 | 0.53 | 0.05 | 10 | 0.89 | 0.03 | 10 | 0.82 | 0.05 | 10 | 0.74 | 0.04 |
| SLF | 0 | 0.57 | 0.04 | 9 | 0.64 | 0.06 | 9 | 0.66 | 0.05 | 3 | 0.55 | 0.08 |
| SS | 0 | 0.51 | 0.04 | 8 | 0.65 | 0.04 | 8 | 0.63 | 0.05 | 0 | 0.53 | 0.04 |
| TAP | 0 | 0.57 | 0.03 | 1 | 0.53 | 0.04 | 5 | 0.57 | 0.05 | 0 | 0.50 | 0.02 |

Table S15.

Model adaptation on the **NIMHANS** dataset. **Extreme deviations** for all four DTI metrics for **dementia**. Numbers indicate the percentage of subjects with dementia with extreme deviations for each DTI metric per ROI. Yellow cells indicate the ROIs that passed multiple comparison testing (FDR) in at least 9 out of 10 experimental repetitions. Also see supplementary Table 4 for the AUCs.

| ROI | FA | | MD | | RD | | AD | |
|---|---|---|---|---|---|---|---|---|
| | Z < -2 | Z > 2 | Z < -2 | Z > 2 | Z < -2 | Z > 2 | Z < -2 | Z > 2 |
| ACR | 3.56 | 1.89 | 1.33 | 13.33 | 1.00 | 11.11 | 0.78 | 15.11 |
| ALIC | 2.33 | 3.00 | 0.22 | 5.56 | 1.56 | 5.44 | 1.11 | 5.78 |
| BCC | 13.89 | 2.00 | 0.89 | 13.11 | 0.67 | 13.22 | 2.00 | 12.11 |
| CGC | 12.11 | 1.44 | 1.00 | 7.44 | 1.22 | 10.78 | 2.22 | 4.44 |
| CGH | 21.33 | 0.56 | 1.22 | 10.67 | 1.11 | 14.56 | 1.89 | 6.11 |
| CST | 3.78 | 3.78 | 1.67 | 4.33 | 1.44 | 3.89 | 3.67 | 3.44 |
| EC | 7.44 | 3.11 | 1.67 | 9.22 | 2.56 | 9.11 | 1.44 | 10.22 |
| FX | 23.00 | 0.00 | 0.00 | 17.33 | 0.00 | 23.56 | 0.11 | 9.56 |
| FXST | 15.33 | 1.00 | 2.33 | 4.11 | 1.33 | 8.22 | 6.44 | 2.11 |
| GCC | 11.00 | 1.44 | 0.56 | 14.33 | 1.22 | 14.44 | 0.00 | 12.22 |
| UNC | 5.56 | 1.11 | 1.78 | 12.11 | 1.78 | 13.22 | 1.11 | 8.67 |
| PCR | 8.67 | 8.00 | 1.22 | 14.56 | 2.33 | 12.89 | 0.22 | 17.00 |
| PLIC | 4.33 | 3.44 | 1.44 | 4.11 | 2.56 | 4.67 | 1.56 | 3.44 |
| PTR | 5.78 | 4.33 | 2.44 | 7.22 | 2.67 | 8.00 | 3.78 | 6.44 |
| RLIC | 3.22 | 0.33 | 0.00 | 6.11 | 0.00 | 5.67 | 0.00 | 6.44 |
| SCC | 10.78 | 0.22 | 0.00 | 12.22 | 0.11 | 11.33 | 0.00 | 12.44 |
| SCR | 3.33 | 6.89 | 0.67 | 7.33 | 1.22 | 5.33 | 0.44 | 12.89 |
| SFO | 2.78 | 4.11 | 0.44 | 6.44 | 0.78 | 5.67 | 1.11 | 9.22 |
| SLF | 6.56 | 3.56 | 2.78 | 12.00 | 2.56 | 11.44 | 1.67 | 10.78 |
| SS | 7.33 | 2.33 | 1.11 | 9.33 | 1.22 | 9.89 | 2.44 | 6.56 |
| TAP | 6.67 | 2.11 | 1.67 | 15.89 | 1.56 | 14.56 | 0.33 | 15.22 |

Table S16.

Results for model adaptation to the **NIMHANS** dataset. **Areas Under the ROC Curve (AUC)** for all four DTI metrics for **dementia**. Numbers indicate the average AUC (across 10 experimental repetitions) and number of passing tests after multiple comparisons correction (FDR_sum) for each DTI metric per ROI. Yellow cells indicate the ROIs that passed multiple comparison testing (FDR) in at least 9 out of 10 experimental repetitions.

| ROI | FA | | | MD | | | RD | | | AD | | |
|---|---|---|---|---|---|---|---|---|---|---|---|---|
| | FDR_sum | AUC (mean/std) | | FDR_sum | AUC (mean/std) | | FDR_sum | AUC (mean/std) | | FDR_sum | AUC (mean/std) | |
| ACR | 1 | 0.50 | 0.05 | 10 | 0.70 | 0.05 | 10 | 0.68 | 0.04 | 9 | 0.66 | 0.06 |
| ALIC | 0 | 0.51 | 0.04 | 10 | 0.83 | 0.02 | 10 | 0.76 | 0.04 | 10 | 0.78 | 0.04 |
| BCC | 8 | 0.65 | 0.05 | 10 | 0.81 | 0.04 | 10 | 0.77 | 0.04 | 10 | 0.81 | 0.05 |
| CGC | 1 | 0.54 | 0.04 | 10 | 0.75 | 0.05 | 10 | 0.68 | 0.03 | 5 | 0.59 | 0.04 |
| CGH | 5 | 0.59 | 0.04 | 10 | 0.69 | 0.03 | 10 | 0.72 | 0.02 | 4 | 0.55 | 0.05 |
| CST | 5 | 0.59 | 0.02 | 10 | 0.81 | 0.03 | 10 | 0.79 | 0.04 | 10 | 0.67 | 0.03 |
| EC | 0 | 0.51 | 0.03 | 10 | 0.71 | 0.03 | 10 | 0.67 | 0.04 | 10 | 0.68 | 0.04 |
| FX | 10 | 0.72 | 0.05 | 10 | 0.67 | 0.04 | 10 | 0.71 | 0.04 | 3 | 0.54 | 0.05 |
| FXST | 2 | 0.56 | 0.03 | 10 | 0.69 | 0.04 | 8 | 0.62 | 0.04 | 8 | 0.65 | 0.06 |
| GCC | 3 | 0.57 | 0.05 | 10 | 0.71 | 0.04 | 10 | 0.68 | 0.03 | 9 | 0.67 | 0.06 |
| UNC | 0 | 0.51 | 0.04 | 10 | 0.76 | 0.03 | 10 | 0.71 | 0.04 | 9 | 0.63 | 0.03 |
| PCR | 6 | 0.61 | 0.04 | 9 | 0.70 | 0.07 | 10 | 0.69 | 0.05 | 9 | 0.67 | 0.06 |
| PLIC | 9 | 0.65 | 0.04 | 10 | 0.83 | 0.04 | 10 | 0.81 | 0.02 | 10 | 0.76 | 0.03 |
| PTR | 10 | 0.70 | 0.02 | 10 | 0.77 | 0.02 | 10 | 0.77 | 0.01 | 9 | 0.62 | 0.03 |
| RLIC | 2 | 0.57 | 0.03 | 10 | 0.67 | 0.05 | 10 | 0.67 | 0.04 | 8 | 0.62 | 0.04 |
| SCC | 9 | 0.66 | 0.03 | 10 | 0.79 | 0.05 | 10 | 0.78 | 0.04 | 10 | 0.69 | 0.04 |
| SCR | 1 | 0.55 | 0.04 | 10 | 0.83 | 0.03 | 10 | 0.77 | 0.03 | 10 | 0.78 | 0.03 |
| SFO | 2 | 0.55 | 0.05 | 10 | 0.85 | 0.03 | 10 | 0.80 | 0.03 | 10 | 0.77 | 0.04 |
| SLF | 7 | 0.60 | 0.03 | 9 | 0.66 | 0.05 | 10 | 0.69 | 0.04 | 8 | 0.65 | 0.07 |
| SS | 6 | 0.61 | 0.04 | 10 | 0.67 | 0.03 | 10 | 0.68 | 0.03 | 2 | 0.56 | 0.03 |
| TAP | 9 | 0.62 | 0.02 | 10 | 0.63 | 0.04 | 10 | 0.68 | 0.04 | 5 | 0.59 | 0.02 |

Table S17.

Model adaptation on the **UCLA** dataset. **Extreme deviations** for all four DTI metrics for **22q11.2 deletion syndrome**. Numbers indicate the percentage of subjects with 22q11.2 deletion syndrome with extreme deviations for each DTI metric per ROI. Yellow cells indicate the ROIs that passed multiple comparison testing (FDR) in at least 9 out of 10 experimental repetitions. Also see supplementary Table 4 for the AUCs.

| ROI | FA | | MD | | RD | | AD | |
|---|---|---|---|---|---|---|---|---|
| | Z < -2 | Z > 2 | Z < -2 | Z > 2 | Z < -2 | Z > 2 | Z < -2 | Z > 2 |
| ACR | 2.39 | 5.21 | 9.15 | 0.14 | 10.28 | 0.14 | 4.23 | 0.42 |
| ALIC | 1.41 | 2.25 | 0.28 | 0.42 | 0.99 | 0.56 | 1.69 | 2.11 |
| BCC | 0.14 | 9.30 | 7.75 | 0.00 | 10.00 | 0.85 | 6.48 | 0.99 |
| CGC | 5.63 | 0.70 | 6.48 | 0.42 | 0.56 | 3.94 | 8.17 | 0.42 |
| CGH | 8.87 | 1.55 | 8.73 | 3.52 | 6.48 | 8.73 | 8.59 | 0.00 |
| EC | 15.35 | 0.00 | 3.38 | 1.41 | 0.00 | 4.65 | 18.17 | 0.00 |
| FX | 25.63 | 1.55 | 0.00 | 25.07 | 0.00 | 29.30 | 0.14 | 16.20 |
| FXST | 10.56 | 0.42 | 3.80 | 1.55 | 1.27 | 4.79 | 7.89 | 0.00 |
| GCC | 0.14 | 6.06 | 7.75 | 1.41 | 8.87 | 1.83 | 3.38 | 2.68 |
| UNC | 3.38 | 6.90 | 9.15 | 0.85 | 8.59 | 2.11 | 10.28 | 4.93 |
| PCR | 2.25 | 10.14 | 21.55 | 0.56 | 18.59 | 0.42 | 11.13 | 0.00 |
| PLIC | 1.41 | 6.34 | 2.11 | 0.28 | 3.24 | 0.56 | 0.56 | 7.46 |
| PTR | 0.14 | 1.41 | 19.15 | 0.14 | 1.83 | 0.42 | 22.11 | 0.00 |
| RLIC | 0.14 | 1.41 | 10.85 | 0.56 | 5.49 | 0.42 | 7.18 | 0.00 |
| SCC | 0.00 | 10.28 | 4.79 | 0.00 | 14.51 | 0.00 | 2.25 | 0.00 |
| SCR | 4.51 | 14.65 | 7.32 | 0.00 | 13.52 | 0.70 | 5.21 | 3.38 |
| SFO | 1.55 | 1.83 | 4.65 | 0.00 | 4.79 | 0.28 | 3.10 | 0.00 |
| SLF | 4.37 | 0.14 | 11.13 | 0.42 | 3.24 | 1.13 | 17.61 | 0.00 |
| SS | 3.94 | 2.82 | 12.39 | 1.41 | 5.63 | 3.24 | 10.85 | 0.00 |
| TAP | 0.00 | 12.82 | 17.04 | 2.96 | 14.23 | 2.11 | 1.83 | 0.00 |

Table S18.

Results for model adaptation to the **UCLA** dataset. **Areas Under the ROC Curve (AUC)** for all four DTI metrics for **22q11.2 Deletion Syndrome.** Numbers indicate the average AUC (across 10 experimental repetitions) and number of passing tests after multiple comparisons correction (FDR_sum) for each DTI metric per ROI. Yellow cells indicate the ROIs that passed multiple comparison testing (FDR) in at least 9 out of 10 experimental repetitions.

| ROI | FA | | | MD | | | RD | | | AD | | |
|---|---|---|---|---|---|---|---|---|---|---|---|---|
| | FDR_sum | AUC (mean/std) | | FDR_sum | AUC (mean/std) | | FDR_sum | AUC (mean/std) | | FDR_sum | AUC (mean/std) | |
| ACR | 0 | 0.39 | 0.08 | 10 | 0.72 | 0.04 | 5 | 0.63 | 0.05 | 4 | 0.63 | 0.04 |
| ALIC | 0 | 0.42 | 0.07 | 1 | 0.56 | 0.03 | 0 | 0.49 | 0.05 | 6 | 0.62 | 0.04 |
| BCC | 0 | 0.50 | 0.07 | 7 | 0.65 | 0.05 | 0 | 0.48 | 0.04 | 9 | 0.68 | 0.04 |
| CGC | 0 | 0.50 | 0.07 | 9 | 0.69 | 0.06 | 4 | 0.61 | 0.08 | 8 | 0.65 | 0.04 |
| CGH | 1 | 0.59 | 0.05 | 5 | 0.62 | 0.07 | 5 | 0.65 | 0.06 | 0 | 0.48 | 0.05 |
| EC | 0 | 0.51 | 0.05 | 0 | 0.54 | 0.04 | 0 | 0.54 | 0.05 | 10 | 0.76 | 0.03 |
| FX | 0 | 0.49 | 0.09 | 0 | 0.45 | 0.04 | 0 | 0.38 | 0.03 | 1 | 0.57 | 0.04 |
| FXST | 1 | 0.52 | 0.07 | 6 | 0.63 | 0.05 | 6 | 0.65 | 0.04 | 1 | 0.51 | 0.06 |
| GCC | 0 | 0.49 | 0.04 | 10 | 0.70 | 0.05 | 0 | 0.43 | 0.06 | 10 | 0.74 | 0.05 |
| UNC | 0 | 0.51 | 0.04 | 0 | 0.56 | 0.03 | 0 | 0.52 | 0.06 | 3 | 0.56 | 0.07 |
| PCR | 0 | 0.46 | 0.06 | 9 | 0.72 | 0.06 | 5 | 0.64 | 0.04 | 5 | 0.62 | 0.08 |
| PLIC | 0 | 0.41 | 0.06 | 8 | 0.65 | 0.03 | 0 | 0.49 | 0.05 | 10 | 0.73 | 0.02 |
| PTR | 0 | 0.32 | 0.05 | 8 | 0.65 | 0.04 | 0 | 0.43 | 0.07 | 7 | 0.66 | 0.06 |
| RLIC | 0 | 0.40 | 0.06 | 9 | 0.68 | 0.05 | 1 | 0.51 | 0.06 | 7 | 0.64 | 0.04 |
| SCC | 4 | 0.67 | 0.04 | 1 | 0.55 | 0.06 | 4 | 0.64 | 0.04 | 0 | 0.48 | 0.04 |
| SCR | 0 | 0.47 | 0.06 | 10 | 0.67 | 0.02 | 4 | 0.63 | 0.03 | 2 | 0.57 | 0.05 |
| SFO | 0 | 0.39 | 0.05 | 6 | 0.62 | 0.04 | 0 | 0.52 | 0.05 | 0 | 0.56 | 0.05 |
| SLF | 0 | 0.40 | 0.06 | 10 | 0.76 | 0.04 | 4 | 0.62 | 0.04 | 10 | 0.75 | 0.04 |
| SS | 0 | 0.49 | 0.05 | 8 | 0.67 | 0.05 | 3 | 0.60 | 0.06 | 7 | 0.64 | 0.04 |
| TAP | 0 | 0.56 | 0.04 | 0 | 0.46 | 0.05 | 0 | 0.50 | 0.05 | 0 | 0.37 | 0.07 |

| Dataset | Scanner Manufacturer | Magnetic Field Strength (T) | TR (ms) | TE (ms) | Diffusion-weighted b-value (s/mm$^2$) | No. of diffusion-weighted dirs. | b-values (volumes) | No. of b0 volumes | Voxel size (mm) | In-plane resolution (voxel) | Slices |
|---|---|---|---|---|---|---|---|---|---|---|---|
| **ABCD_Siemens** | Siemens | 3 | 4100 | 88 | 1000 | 15 | 1000 (15) | 7 | 1.70 x 1.70 x 1.70 | 140 x 140 | 81 |
| **ABCD_GE** | GE | 3 | 4100 | 81.9 | 1000 | 15 | 1000 (15) | 7 | 1.70 x 1.70 x 1.70 | 140 x 140 | 81 |
| **ABCD_Philips** | Philips | 3 | 5300 | 89 | 1000 | 15 | 1000 (15) | 7 | 1.70 x 1.70 x 1.70 | 140 x 140 | 81 |
| **ADNI3_GE36** | GE | 3 | 9000-16030 | 55.5-75.7 | 1000 | 32 | 1000 (32) | 4 | 0.91 x 0.91 x 2.00 | 256 x 256 (zero-filled) | 80 |
| **ADNI3_GE54** | GE | 3 | 7800-9000 | 54.7-76.0 | 1000 | 48 | 1000 (48) | 6 | 0.91 x 0.91 x 2.00 | 256 x 256 (zero-filled) | 80 |
| **ADNI3_P33** | Philips | 3 | 9916-10860 | 85.7-100 | 1000 | 32 | 1000 (32) | 1 | 2.00 x 2.00 x 2.00 | 128 x 128 | 80 |
| **ADNI3_P36** | Philips | 3 | 9953-11200 | 86.8-101 | 1000 | 32 | 1000 (32) | 4 | 2.00 x 2.00 x 2.00 | 128 x 128 | 80 |
| **ADNI3_S127** | Siemens | 3 | 3400-4200 | 71-99 | 1000 | 48 | 1000 (48) | 13 | 2.00 x 2.00 x 2.00 | 116 x 116 | 81 |
| **ADNI3_S31** | Siemens | 3 | 12400-16700 | 95-105 | 1000 | 30 | 1000 (30) | 1 | 2.00 x 2.00 x 2.00 | 116 x 116 | 80 |
| **ADNI3_S55** | Siemens | 3 | 7200-10100 | 56-82 | 1000 | 48 | 1000 (48) | 7 | 2.00 x 2.00 x 2.00 | 116 x 116 | 80 |
| **AOMIC_ID1000** | Philips | 3 | 6312 | 74 | 1000 | 32 (x3) | 1000 (96) | 3 | 2.00 x 2.00 x 2.00 | 112 x 112 | 60 |
| **AOMIC_PIOP1** | Philips | 3 | 7387 | 86 | 1000 | 32 | 1000 (32) | 1 | 2.00 x 2.00 x 2.00 | 112 x 112 | 60 |
| **AOMIC_PIOP2** | Philips | 3 | 7387 | 86 | 1000 | 32 | 1000 (32) | 1 | 2.00 x 2.00 x 2.00 | 112 x 112 | 60 |
| **CAMCAN** | Siemens | 3 | 9100 | 104 | 1000 | 30 | 1000 (30) | 3 | 2.00 x 2.00 x 2.00 | 96 x 96 | 66 |
| **CHBMP** | Siemens | 1.5 | 7000 | 160 | 1200 | 12 | 1200 (12) | 1 | 2.00 x 2.00 x 3.00 | 128 x 128 | 50 |
| **CHCP** | Siemens | 3 | 3500 | 86 | 1000 | 46 (x2 AP/PA) | 1000 (46) | 7 | 1.50 x 1.50 x 1.50 | 140 x 140 | 100 |
| **HCP_A** | Siemens | 3 | 3230 | 89.2 | 1500 | 92 (x2 AP/PA) | 1500 (92) | 28 | 1.50 x 1.50 x 1.50 | 140 x 140 | 92 |
| **HCP_D** | Siemens | 3 | 3230 | 89.2 | 1500 | 92 (x2 AP/PA) | 1500 (92) | 28 | 1.50 x 1.50 x 1.50 | 140 x 140 | 92 |
| **HCP_YA** | Siemens | 3 | 5520 | 89.5 | 1000 | 90 (x2 LR/RL) | 1000 (90) | 12 | 1.25 X1.25 x 1.25 | 168 x 144 | 111 |
| **NIMHANS_Philips** | Philips | 3 | 7441 | 85 | 1000 | 64 | 1000 (64) | 2 | 2.00 x 2.00 x 2.00 | 112 x 112 | 64 |
| **NIMHANS_Siemens** | Siemens | 3 | 8400 | 91 | 1000 | 64 | 1000 (64) | 1 | 1.97 x 1.97 x 2.00 | 122 x 122 | 61 |
| **OASIS3** | Siemens | 3 | 11000 | 87 | 1000 | 64 | 1000 (64) | 1 | 2.50 x 2.50 x 2.50 | 96 x 96 | 64 |
| **PedsDTI_GE** | GE | 1.5 | 6000-17000 | 75-102 | 1000 (Objective 1) | 6 (x4) | 1000 (24) | 4 | 3.00 x 3.00 x 3.00 | variable per subject | 48-60 |
| **PedsDTI_Siemens** | Siemens | 1.5 | 6000-17000 | 75-102 | 1000 (Objective 1) | 6 (x4) | 1000 (24) | 4 | 3.00 x 3.00 x 3.00 | variable per subject | 48-60 |
| **PedsDTI_eDTI_GE** | GE | 1.5 | 8900-17060 | 69-94 | 100, 300, 500, 800, 1100 | 110 | 100 (10), 300 (10), 500 (10), 800 (30), 1100 (50) | 9 | 2.50 x 2.50 x 2.50 | 96x96 | 60 |
| **PedsDTI_eDTI_Siemens** | Siemens | 1.5 | 8900-17060 | 69-94 | 100, 300, 500, 800, 1100 | 110 | 100 (10), 300 (10), 500 (10), 800 (30), 1100 (50) | 10 | 2.50 x 2.50 x 2.50 | 96x96 | 60 |
| **PING_GE** | GE | 3 | 13600 | 83 | 1000 | 30 | 1000 (30) | 2 | 2.50 x 2.50 x 2.50 | 96 x 96 | 51 |
| **PING_Siemens** | Siemens | 3 | 20000 | 94 | 1000 | 30 | 1000 (30) | 2 | 2.50 x 2.50 x 2.50 | 96 x 96 | 64 |
| **PING_Philips** | Philips | 3 | 9000 | 91 | 1000 | 30 | 1000 (30) | 2 | 2.50 x 2.50 x 2.50 | 96 x 96 | 60 |
| **PNC** | Siemens | 3 | 8100 | 82 | 1000 | 64 | 1000 (64) | 7 | 1.88 x 1.88 x 2.00 | 128 x 128 | 70 |
| **PPMI** | Siemens | 3 | 9000 | 88 | 1000 | 64 | 1000 (64) | 1 | 1.98 x 1.98 x 2.00 | 116 x116 | 72 |
| **QTAB** | Siemens | 3 | 3800 | 70 | 1000 | 20 | 1000 (20) | 12 | 2.00 x 2.00 x 2.00 | 122 x 122 | 68 |
| **QTIM** | Bruker | 4 | 6090 | 91.7 | 1149 | 94 | 1149 (94) | 11 | 1.79 x 1.79 x 2.00 | 128 x 128 | 55 |
| **SLIM** | Siemens | 3 | 11000 | 98 | 1000 | 30 (x3) | 1000 (90) | 3 | 2.00 x 2.00 x 2.00 | 128 x 128 | 60 |
| **UKBB** | Siemens | 3 | 3600 | 92 | 1000 | 50 | 1000 (50) | 5 | 2.01 x 2.01 x 2.01 | 104 x 104 | 72 |
| **HBN_CBIC** | Siemens | 3 | 3320 | 100.2 | 1000 | 64 | 1000 (64) | 1 | 1.80 x 1.80 x 1.80 | 104 x 104 | 72 |
| **HBN_CUNY** | Siemens | 3 | 3320 | 100.2 | 1000 | 64 | 1000 (64) | 1 | 1.80 x 1.80 x 1.80 | 104 x 104 | 72 |
| **HBN_RU** | Siemens | 3 | 3320 | 100.2 | 1000 | 64 | 1000 (64) | 1 | 1.80 x 1.80 x 1.80 | 104 x 104 | 72 |
| **HBN_SI** | Siemens | 1.5 | 4500 | 93.8 | 1000 | 64 | 1000 (64) | 1 | 2.00 x 2.00 x 2.00 | 96 x 96 | 72 |
| **UCLA_Prisma** | Siemens | 3 | 3230 | 89.2 | 1500 | 92 (x2 AP/PA) | 1500 (92) | 28 | 1.50 x 1.50 x 1.50 | 140 x 140 | 92 |
| **UCLA_Trio** | Siemens | 3 | 7100 | 93 | 1000 | 64 | 1000 (64) | 1 | 1.97 x 1.97 x 2.00 | 96 x 96 | 50 |

**Table S19.** Acquisition Protocols. Diffusion MRI acquisition protocols used throughout all the experiments described in this manuscript. We identified 37 protocols across the studies. For instance, one study - ADNI3 - used seven distinctive protocols. The 37 protocols were modeled as a batch effect for the Hierarchical Bayesian Regression.

Table S20.

FA: Student's unpaired two-sample t-test between kurtosis values of QCed vs. non QCed sites.

| ROI | t | p.value | conf.int.low | conf.int.high |
|---|---|---|---|---|
| Average | -1.194 | 0.2480 | -0.555 | 0.153 |
| ACR | -1.132 | 0.2726 | -0.506 | 0.152 |
| ALIC | 0.452 | 0.6567 | -0.176 | 0.273 |
| BCC | 0.626 | 0.5393 | -0.357 | 0.659 |
| CGC | -0.341 | 0.7370 | -0.252 | 0.182 |
| CGH | -0.515 | 0.6125 | -0.373 | 0.226 |
| CST | 1.186 | 0.2511 | -0.113 | 0.405 |
| EC | -0.838 | 0.4132 | -0.233 | 0.100 |
| FX | 4.560 | 0.0002 | 0.462 | 1.251 |
| FXST | 0.267 | 0.7928 | -0.137 | 0.177 |
| GCC | 1.976 | 0.0636 | -0.018 | 0.582 |
| PCR | 0.262 | 0.7961 | -0.193 | 0.248 |
| PLIC | 0.582 | 0.5675 | -0.242 | 0.427 |
| PTR | -0.850 | 0.4067 | -0.505 | 0.214 |
| RLIC | -1.823 | 0.0850 | -0.492 | 0.035 |
| SCC | -0.805 | 0.4313 | -8.904 | 3.971 |
| SCR | -0.555 | 0.5856 | -0.589 | 0.343 |
| SFO | 1.051 | 0.3070 | -0.109 | 0.328 |
| SLF | -0.717 | 0.4825 | -0.475 | 0.233 |
| SS | -1.372 | 0.1870 | -0.460 | 0.097 |
| TAP | -0.754 | 0.4605 | -0.122 | 0.057 |
| UNC | -0.752 | 0.4620 | -0.228 | 0.108 |

Table S21.

MD: Student’s unpaired two-sample t-test between kurtosis values of QCed vs. non QCed sites.

| ROI | t | p.value | conf.int.low | conf.int.high |
|---|---|---|---|---|
| Average | -1.631 | 0.1202 | -0.862 | 0.108 |
| ACR | -1.625 | 0.1216 | -2.018 | 0.258 |
| ALIC | 0.692 | 0.4977 | -0.532 | 1.055 |
| BCC | 2.073 | 0.0528 | -0.028 | 4.083 |
| CGC | 0.301 | 0.7665 | -0.408 | 0.545 |
| CGH | -0.520 | 0.6096 | -0.478 | 0.288 |
| CST | 0.631 | 0.5362 | -0.448 | 0.831 |
| EC | -0.352 | 0.7288 | -1.317 | 0.939 |
| FX | 4.640 | 0.0002 | 1.764 | 4.684 |
| FXST | 0.098 | 0.9230 | -0.364 | 0.399 |
| GCC | -0.335 | 0.7415 | -0.563 | 0.408 |
| PCR | 1.207 | 0.2432 | -0.299 | 1.107 |
| PLIC | 0.359 | 0.7237 | -2.418 | 3.415 |
| PTR | -0.399 | 0.6943 | -0.615 | 0.418 |
| RLIC | -0.864 | 0.3989 | -0.490 | 0.204 |
| SCC | -0.774 | 0.4492 | -4.141 | 1.912 |
| SCR | -1.497 | 0.1518 | -2.278 | 0.383 |
| SFO | -0.945 | 0.3569 | -3.435 | 1.303 |
| SLF | -0.649 | 0.5243 | -0.989 | 0.522 |
| SS | -0.242 | 0.8118 | -0.421 | 0.334 |
| TAP | 1.834 | 0.0833 | -0.132 | 1.940 |
| UNC | 0.213 | 0.8339 | -0.710 | 0.870 |

Table S22.

AD: Student’s unpaired two-sample t-test between kurtosis values of QCed vs. non QCed sites.

| ROI | t | p.value | conf.int.low | conf.int.high |
|---|---|---|---|---|
| Average | 1.368 | 0.18823 | -0.093 | 0.439 |
| ACR | -1.611 | 0.12462 | -1.797 | 0.237 |
| ALIC | -0.336 | 0.74091 | -0.582 | 0.421 |
| BCC | 2.765 | 0.01277 | 0.157 | 1.153 |
| CGC | 0.320 | 0.75302 | -0.149 | 0.202 |
| CGH | -1.314 | 0.20551 | -0.318 | 0.073 |
| CST | -0.392 | 0.69994 | -0.626 | 0.430 |
| EC | -0.774 | 0.44922 | -1.298 | 0.599 |
| FX | 5.748 | 0.00002 | 0.990 | 2.131 |
| FXST | -0.894 | 0.38318 | -0.686 | 0.276 |
| GCC | 2.307 | 0.03318 | 0.015 | 0.326 |
| PCR | -0.350 | 0.73058 | -0.945 | 0.675 |
| PLIC | -0.360 | 0.72290 | -1.109 | 0.785 |
| PTR | -0.817 | 0.42461 | -0.374 | 0.165 |
| RLIC | 0.868 | 0.39656 | -0.138 | 0.333 |
| SCC | 0.364 | 0.72039 | -0.216 | 0.307 |
| SCR | -1.342 | 0.19615 | -3.573 | 0.787 |
| SFO | -0.538 | 0.59737 | -1.420 | 0.842 |
| SLF | 0.530 | 0.60242 | -0.195 | 0.326 |
| SS | -0.606 | 0.55210 | -0.357 | 0.197 |
| TAP | 0.780 | 0.44534 | -0.332 | 0.723 |
| UNC | -0.488 | 0.63119 | -0.376 | 0.234 |

Table S23.

RD: Student's unpaired two-sample t-test between kurtosis values of QCed vs. non QCed sites.

| ROI | t | p.value | conf.int.low | conf.int.high |
|---|---|---|---|---|
| Average | -1.481 | 0.1559 | -1.025 | 0.177 |
| ACR | -1.454 | 0.1633 | -1.917 | 0.349 |
| ALIC | 0.880 | 0.3906 | -0.401 | 0.980 |
| BCC | 2.253 | 0.0370 | 0.077 | 2.203 |
| CGC | -0.601 | 0.5552 | -0.367 | 0.204 |
| CGH | 0.228 | 0.8225 | -0.405 | 0.504 |
| CST | 1.327 | 0.2012 | -0.151 | 0.669 |
| EC | -0.260 | 0.7976 | -0.988 | 0.770 |
| FX | 4.751 | 0.0002 | 2.419 | 6.255 |
| FXST | 0.396 | 0.6970 | -0.230 | 0.337 |
| GCC | 0.620 | 0.5427 | -0.332 | 0.610 |
| PCR | 1.965 | 0.0651 | -0.034 | 1.025 |
| PLIC | -0.123 | 0.9032 | -0.730 | 0.649 |
| PTR | -0.553 | 0.5872 | -0.784 | 0.458 |
| RLIC | -1.341 | 0.1967 | -0.773 | 0.171 |
| SCC | -0.760 | 0.4573 | -8.118 | 3.806 |
| SCR | -0.786 | 0.4419 | -0.759 | 0.346 |
| SFO | -1.192 | 0.2489 | -3.380 | 0.934 |
| SLF | -0.357 | 0.7249 | -0.801 | 0.568 |
| SS | -0.549 | 0.5896 | -0.474 | 0.278 |
| TAP | 2.143 | 0.0460 | 0.013 | 1.343 |
| UNC | -0.451 | 0.6577 | -0.531 | 0.343 |

Supplementary Methods.

1. Preprocessing pipeline used for each dataset

| Study/Dataset | Denoising | Gibbs correction | EPI induced distortion correction | Eddy correction | DTI fitting software |
|---|---|---|---|---|---|
| ABCD | *Preprocessed by the ABCD Consortium** | | | | FSL |
| AOMIC | *Preprocessed by AOMIC (U. of Amsterdam)** | | | | DiPy |
| CAMCAN | DiPy MPPCA | DiPy | TOPUP-SynB0 | FSL-EDDY | DiPy |
| CHBMP | DiPy MPPCA | DiPy | TOPUP | TORTOISE | DiPy |
| CHCP | DiPy MPPCA | DiPy | TOPUP | FSL-EDDY | DiPy |
| HBN | DiPy MPPCA | Not done | TOPUP | FSL-EDDY | FSL |
| HCP-A | DiPy MPPCA | DiPy | TOPUP | FSL-EDDY | DiPy |
| HCP-D | DiPy MPPCA | DiPy | TOPUP | FSL-EDDY | DiPy |
| HCP-YA | DiPy Patch2Self | *Preprocessed by the HCP Consortium**** | | | DiPy |
| PedsDTI | *Preprocessed by the NIH** | | | | |
| PING | Not done | Not done | T1 warp | FSL eddy_correct | FSL |
| PNC | DiPy MPPCA | DiPy | TOPUP-SynB0 | FSL-EDDY | DiPy |
| QTAB | DiPy MPPCA | DiPy | TOPUP | FSL-EDDY | DiPy |
| QTIM | Not done | Not done | T1 warp | FSL eddy_correct | FSL |
| SLIM | Not done | Not done | T1 warp | FSL eddy_correct | FSL |
| UKBB | *Preprocessed by the UKBB** | | | | |
| ADNI3 | DiPy LPCA/MPPCA** | MRtrix3 | T1 warp | FSL-EDDY | FSL |
| OASIS3 | DiPy MPPCA | MRtrix3 | TOPUP-SynB0 | FSL-EDDY | FSL |
| PPMI | Not done | Not done | TOPUP-SynB0 | FSL-EDDY | FSL |
| NIMHANS | DiPy LPCA | DiPy | TOPUP-SynB0 | FSL-EDDY | DiPy |
| UCLA | DiPy LPCA | DiPy | TOPUP | FSL-EDDY | DiPy |

** Please check the corresponding reference in the 'References' section of the paper.*
*** Depending on scanner vendor, one or the other was used according to Thomopoulos et al. 2021[23].*
**** Denoising performed on the downloaded preprocessed dMRI data.*

2. Hierarchical Bayesian Regression specific parameters

This approach differs from the complete pooling approach used by ComBat[4]. ComBat-based methods have recently been introduced for cross site[4] data harmonization in DTI with respect to a previously trained harmonized template[5]. Complete pooling harmonizes the individual data by adjusting for multiplicative and additive batch effects and then feeding the corrected data as an input to estimate the normative model. Normative modeling with HBR adjusts the Z-scores instead of the input data points. In a pilot multi-site study, we used this approach to detect extreme deviations from the norm in small samples of participants with neurogenetic disorders[6]. Thus, NM of multisite microstructural brain metrics holds potential for clinical application in psychiatry and neurology. In this paper we used the following options for the HBR algorithm available in PCNtoolkit:

- Batch effect modeling for the site (n=37) and sex (n=2).
- Random intercept for each batch effect: site and sex.
- Standardization of the output response, i.e., the ROI per DTI metric.

**3.** Outlier removal with isolation forests

"Isolation Forest is an algorithm for data anomaly detection using binary trees. It was developed by Fei Tony Liu in 2008[7]. It has a linear time complexity and a low memory use, which works well for high-volume data." (taken from Wikipedia). In this paper we used the implementation of ScikitLearn: https://scikit-learn.org/stable/modules/generated/sklearn.ensemble.IsolationForest.html.

The goal of using an automatic tool such as the isolation forests is to automate the quality check procedure and reduce the effects of inter-rater variability which arise when relying solely on manual QC. One may opt to train a set of initial normative models to find the outliers, remove them, and then retrain the normative models without the outliers[8].

**4.** Handling of site differences

We also acknowledge though that the so-called "site" effect in neuroimaging multi-center studies is an aggregate effect arising from many variables, including scanner acquisition parameters, preprocessing softwares and study specific inclusion and exclusion criteria, among others. In this paper we selected the scanning parameters as the determining factor for considering a group of scans a different "site" in the model (see Supplementary **Table S19)**. The scanning parameters are also multivariate by nature, including the scanner vendor, magnetic field strength, echo time, repetition time, voxel size, number of gradient directions, b-value, number of 'b-zero' scans and software changes. Importantly, some preprocessing steps are deliberately adapted to handle these specific acquisition parameters and thus these tend to be correlated. One particular combination of the above parameters makes one "site" in our modeling approach for HBR, and by following this method we identified 37 sites. To check the model fit for each of these sites, we added four figures showing evaluation metrics (SMSE, MSLL and Rho) per site for each DTI metric (**Supplementary figures S5, S6, S7, S8**).

If a site using the proposed model adaptation method were concerned that the Z-scores might reflect artifacts of acquisition and processing inconsistencies, it is possible for that site to undertake a calibration test on their own healthy controls. As noted throughout, the calibrated assignment of Z-scores to any new sample can be tested using QQ-plots, pinball loss (to detect mean and maximum errors across centiles) and even formal tests of the statistical moments of the resulting distribution, that would flag any departures from normality.

5. Modeling sex

We chose to include sex as a "batch" or "group" effect, as this is a stable approach to model sex for training normative models of other image-derived phenotypes (IDPs), such as FreeSurfer metrics derived from T1-weighted structural images[9]. In hierarchical Bayesian models, group-level effects are used to model hierarchical or multilevel data - that is, data where observations are nested within groups (e.g., students within schools, patients within hospitals, or measures across different sexes, regions or sites). Here, in the context of HBR, "group" refers to any non-ordinal categorical variable, such as sex or scanner, that may introduce batch effects into the data. For clarification, we use the terms 'batch' or 'group' interchangeably to refer to data that are collected with different acquisition protocols (see dMRI sequence specifics in **Table S19**). However, the HBR formulation for site can also be applied to other biologically relevant group-effects (e.g., sex or ethnicity). Group-level effects allow the model to estimate parameters that vary by group, while also sharing information across groups through a common distribution, known as "partial pooling". As we included a "random intercept" option when training the models, the trajectories for each sex across the lifespan had a different intercept, as depicted in **Figure 2B**. The same is true for the site effect, as shown in **Figure 1B**.

Kia et al.[9] compared two approaches for a multi-site normative modeling scenario: modeling site as a fixed versus a batch-effect. In terms of model fitting performances, the HBR and fixed-effect modeling showed equivalent regression performances. Both were also better when compared to approaches including harmonization of the data before running normative modeling (e.g., ComBat). Taking these considerations into account, we modeled sex and site as group effects in the HBR model. We modeled the difference across group effects by estimating random intercepts across groups (i.e., same curve with different offsets in metrics). An additional possibility is to estimate variable slopes (i.e., using 'random-slope' option to estimate different curves) for each sex on top of variable intercepts in our HBR models. This would add extra complexity to the model as the total number of slope parameters is multiplied by the number of sites and sexes. The total number of parameters would be related to the number of knots of the B-spline and is proportional (depending on the likelihood) to **(n_covariate * n_knots) + n_groups** in the random intercept case, and **(n_covariate * n_knots + 1) * n_groups** in the random slope case (i.e., the effect of the number of groups (n_groups) on the total number of parameters becomes multiplicative). As can be seen, this option comes at a cost, which is the increase in the number of parameters. We are using B-splines with 5 knots plus age as a covariate, and sex and site as group effects.

#### Supplementary Discussion

We observed a peculiar lifespan trajectory for a few WM regions. This aspect is characterized by a 'double peak', around age 50 to 60 years. For instance, the SCC for FA had a second peak at age 73 years of age. Similarly, the PLIC, RLIC, SCR, and SFO also showed similar trajectories for FA, and the CST, FX, CGH, and PTR for diffusivity metrics. This has been reported before for DTI FA: Zhu et al. reported a similar pattern for FA in SCC[5]. Bouhrara et al.[10] reported that FA displayed a more heterogeneous pattern–where some regions, such as the SCR, had higher values with age. Burzynska et al.[11] also found no decrease in FA for the splenium with age. Westlye et al. reported that cingulum bundles showed a less clear-cut or 'non-U-shaped' trajectory[12]. Sullivan et al.[13] did not find significant linear age effects for CGH and the splenium in men. Similar results have been reported in longitudinal studies: in Storsve et al.[14], the *forceps major*, a tract running through the SCC, showed no longitudinally significant changes for FA, similar to the left cingulum angular bundle and left ILF. However, the magnitude of this double bump trajectory is much lower than the estimated variance in the sample. That is to say, that the overall, global variance of the estimated model is larger than the local variance of a specific age range or even for the variance explaining the sex batch effects. This can be observed in Supplementary **Figures S1-S4.** Finally, from the dMRI modeling perspective, DTI is a non-specific microstructural model. FA is a metric sensitive to a combination of multiple tissue properties, including axonal density, myelination, axonal orientation dispersion, crossing fibers, and free-water. Free-water may predominate in many voxels as atrophy and tissue shrinkage occur after age ~50 years and could cause these local effects on the age trajectory of the WM region. Other microstructural properties, such as dispersion, cannot be ruled out - a key topic for future work.
